# Interference-Free Propagation: Achieving Reliable Signal Propagation in Cortical Networks with Areal-Specific Local Dynamics

**DOI:** 10.64898/2026.05.28.728604

**Authors:** Guanchun Li, Zhenyuan Jin, Xiao-Jing Wang, Songting Li

**Author notes:** Contributing authors.

## Abstract

In cortical networks, how to simultaneously achieve both reliable inter-areal signal propagation and areal-specific local dynamics remains largely unclear. In general, strong connections between areas enhance signal propagation, but blend timescales of area-specific neuronal activity, whereas weak connections have the opposite effect. Here we identify a novel dynamical regime termed “interference-free propagation” (IFP) that reconciles the two contrasting demands in the cortex. In the IFP regime, mean signals from upstream areas can propagate reliably, but fluctuations indicative of upstream areas’ timescales are filtered out. This result provides new insights into the operational regime of the cortex, leading to the coexistence of reliable signal propagation and the distinct property of local temporal integration of information in the cortical network.

## 1 Introduction

Brain computation is increasingly recognized as an emergent property of large-scale, multi-areal cortical networks. The cortex comprises many interacting areas with distinct anatomical and physiological characteristics (areal heterogeneity) [1, 2], embedded within dense recurrent connectivity. This heterogeneity and interdependence pose fundamental challenges for understanding how the brain processes and propagates information, highlighting the need for new conceptual frameworks that integrate brain-wide communication with local circuit specialization.

Two key phenomena observed in the cortex underscore this challenge: timescale localization and signal propagation. A hierarchy of intrinsic timescales across cortical areas is predicted by a connectome-based macaque monkey cortex model [3]: from rapid millisecond-scale fluctuations in lower-order sensory areas to more prolonged patterns spanning seconds in higher-order association areas. This timescale hierarchy has been found experimentally in multiple species, including mice [4, 5], macaque monkeys [6–11], and humans [12–19]. The universality of timescale hierarchy is fundamental for cognitive processes, allowing the cortex to integrate information hierarchically over diverse timescales.

Equally important, reliable inter-areal propagation of neural signals ensures that distributed cortical areas can communicate and work in concert. Long-range signal propagation in the cortex is largely determined by the axonal fibers [20–22] and the geometry of the brain [23–25]. In addition, the signal pathway can be shaped by delicate cortical features including near-minimal path length and high clustering [26, 27], assortative community structure [28], a densely interconnected core in the cortex [29], and the hierarchical organization of the brain [30, 31]. Signal propagation in the cortex has often been investigated in simplified scenarios such as feedforward networks [32–36]. However, signal propagation in the highly recurrent cortical network with the abundance of feedback loops [37–39] and areal heterogeneity [40, 41] has not been fully investigated.

The phenomena of timescale hierarchy and reliable signal propagation raise a fundamental theoretical puzzle: How can neural signals reliably propagate through a recurrently connected cortical network without disrupting each area’s distinctive temporal behavior? Given the dense reciprocal connectivity in cortex, one might anticipate that slow fluctuations from higher-order areas would interfere lower-order areas (or vice versa), homogenizing timescales across the network. Yet in reality, cortical areas robustly preserve their distinct temporal characteristics despite substantial inter-areal interaction. Resolving how the cortex can be simultaneously segregated in timescale and integrated in communication across areas is critical for understanding cortical function.

Previous computational models have advanced our understanding of cortical timescale localization and signal propagation, yet also illustrate the inherent tradeoffs between the two. A large-scale model by Chaudhuri et al. (2015) [3] incorporated realistic anatomical connectivity of the macaque cortex and demonstrated that a macroscopic gradient of recurrent excitation across cortical areas naturally gives rise to a hierarchy of timescales. Further model analysis revealed that both the macroscopic gradient of excitation and the detailed excitation-inhibition (E-I) balance are important to achieve localized timescale for each area [2, 3, 42]. In particular, the E-I balance condition necessitates that excitatory inputs from source areas need to cancel with inhibitory inputs within the target area [42]. Consequently, the modeled cortical areas exhibit effectively weak interactions, fostering the emergence of hierarchical timescale localization in a densely connected network [3, 42]. However, the detailed E-I balance condition poses a challenge for reliable signal propagation.

In contrast, a subsequent study by Joglekar et al. (2018) [31] directly addressed the propagation issue. They found that signals face significantly less attenuation when the network operates within a different dynamical regime known as “global balanced amplification”, originating from the concept of balanced amplification that selectively amplifies inputs when feedback inhibition stabilizes strong excitation [43]. However, this regime pushes the system toward greater excitability and loses the timescale hierarchy. Thus, existing theoretical frameworks have either emphasized timescale localization at the expense of effective propagation or vice versa. A unifying theoretical framework satisfying both constraints has remained elusive.

Here we propose a novel operating regime of cortical circuits, interference-free propagation (IFP), which resolves the trade-off between timescale localization and reliable inter-areal transmission. In the IFP regime, a balance of feedforward excitation and inhibition separates upstream inputs into mean and fluctuating components, such that only the mean is propagated while upstream temporal fluctuations containing upstream timescale are suppressed. We formalize this principle using two theoretical metrics—*M*_*SP*_ capturing mean signal propagation and *M*_*T L*_ capturing timescale localization—and show that IFP consistently yields large *M*_*SP*_ with negligible *M*_*T L*_. Through minimal models, analytical theory, large-scale cortical network models and experimental data analysis, we demonstrate that IFP emerges as a robust and biologically plausible mechanism in the cortex. By reconciling timescale hierarchy with inter-areal communication, IFP provides a unifying framework linking local cortical dynamics to large-scale network functionality.

## 2 Results

### 2.1 Purely Excitatory Circuit Mixes Timescales under Strong Coupling

We started by analyzing a simple feedforward model consisting of two connected brain areas labeled *j* (upstream) and *i* (downstream), where each area contains a single excitatory neuronal population. The upstream population drives the downstream population excitatorily (Figure 1a). Mathematically, the dynamics of the excitatory-to-excitatory (*E-to-E*) model can be described as follows:

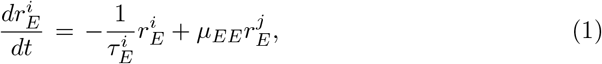

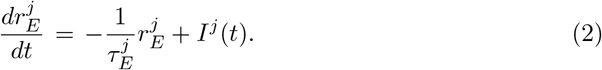

**Fig. 1.**
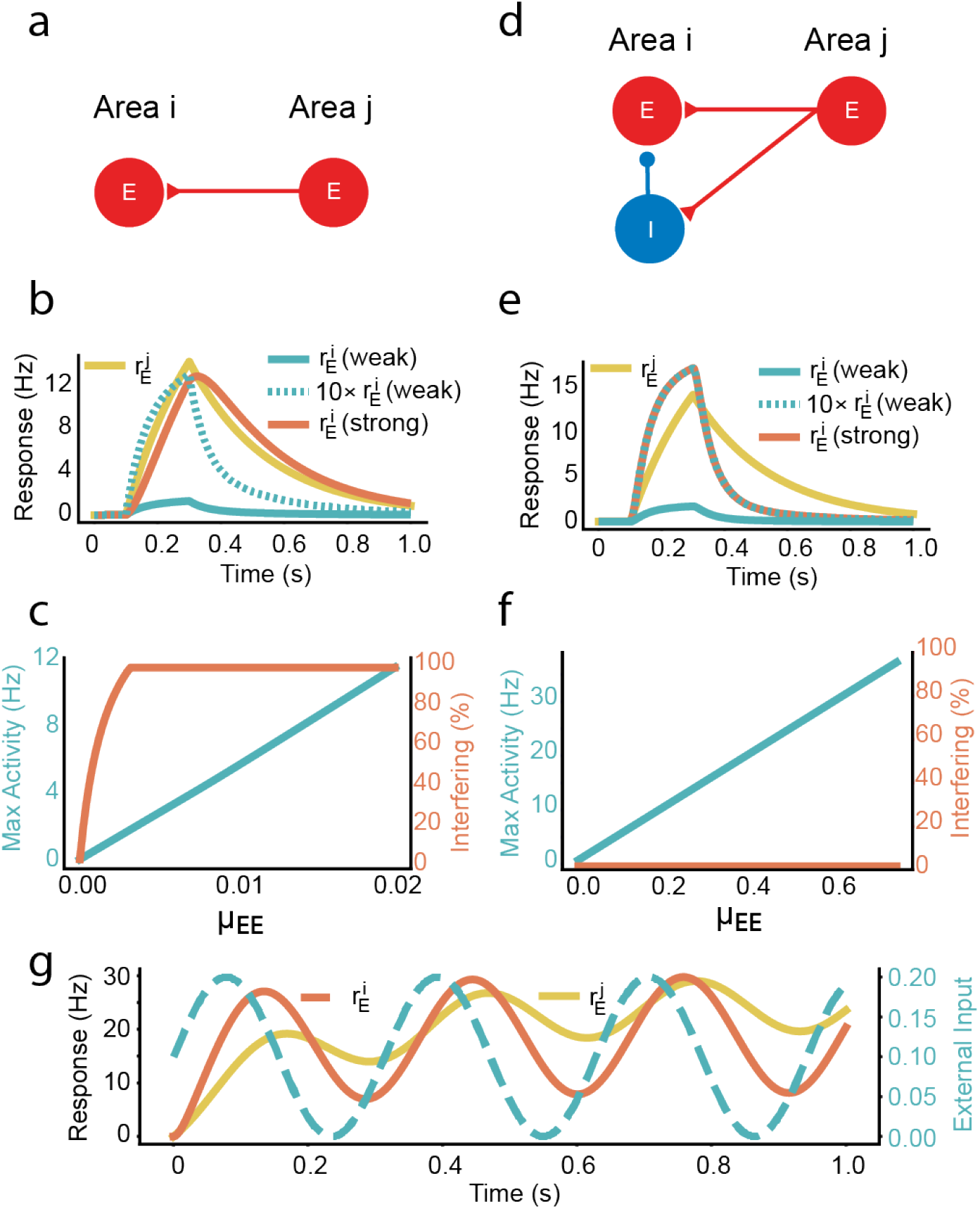
Dynamics of the *E-to-E model* and *E-to-EI model*. (a) Connectivity motif of the *E-to-E model*, consisting of a single excitatory population in each area (red node) with *E* to *E* unidirectional coupling (red arrow). (b) Response of the *E-to-E model* to an brief external pulse input. Yellow: firing rate of upstream population 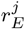. Blue and red: firing rates of downstream population 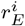 under weak and strong connection strengths. (Dash blue: downstream population 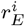 under weak connection, scaled up by a factor of 10.) A small independent external pulse input is applied to the downstream area for visibility. (c) Signal propagation (blue curve, left vertical axis) and timescale interference (red curve, right vertical axis; 100% means downstream timescale fully matches upstream timescale) in the *E-to-E model* with varying connection strengths. (d-f) Same as (a-c) for the *E-to-EI model*. (g) Response of the *E-to-EI model* to oscillatory external input (blue).

Here, 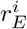 and 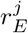 represent the population firing rates of excitatory neuronal populations in brain areas *i* and *j*, respectively. The parameters 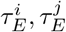 denote the characteristic time constant of the these neuronal populations. We assume distinct time constants in the two areas in order to explore relevant dynamics. In addition, *μ*_*EE*_ specifies the excitatory connection strength from area *j* to *i*, while *I*^*j*^(*t*) is an external input to the upstream area *j*.

This minimal model reveals the interplay between connection strength *μ*_*EE*_, and the dual goals of intra-areal signal integration with distinct timescales and inter-areal signal propagation. As shown in Figure 1b, weak connection allows the downstream area to respond with its own characteristic timescale distinct from the timescale of the upstream area, but yields only a small downstream response (poor signal propagation). In contrast, a strong coupling produces a large downstream response (robust signal propagation) at the expense of the downstream activity inheriting the upstream area’s timescale.

We introduced an interference metric (see Methods) to quantify the degree to which the timescale of the upstream area influences the dynamics of the downstream area. The perfect timescale localization in the downstream is defined as an interference metric of zero. Figure 1c shows that, within the E-to-E model, it is infeasible to simultaneously achieve strong timescale localization (indicated by a low interference metric) and robust signal propagation (indicated by a high peak downstream activity). Additionally, we observed that if the downstream area has a slower characteristic timescale than the upstream area, it could show a large, slow response under strong coupling conditions. However, even in this scenario, the effective downstream timescale remains significantly influenced by the upstream timescale (Figure S1).

### 2.2 Local Inhibition Allows Reliable Propagation without Timescale Mixing

In the cortex, local inhibitory circuits are ubiquitous. Therefore, we extended the two-area model to include an inhibitory neuronal population in the downstream area, yielding an excitatory-to-excitatory-and-inhibitory (*E-to-EI*) model. In this model, the upstream excitatory population in area *j* projects to *both* excitatory and inhibitory populations in downstream area *i*. Figure 1d illustrates this motif. The dynamics can be described as:

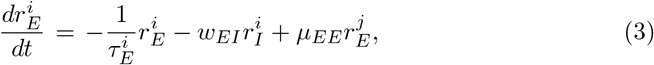

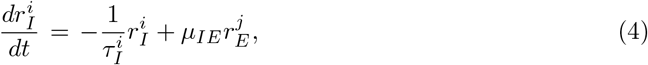

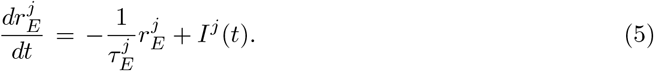

Here the parameter 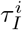 is the characteristic time constant of the inhibitory population of area *i, w*_*EI*_ denotes the intra-areal coupling strength from the inhibitory population to the excitatory population, and *μ*_*IE*_ represents the coupling strength from the upstream excitatory population to the downstream inhibitory population. Other parameters retain their definitions from the *E-to-E* model.

Notably, the *E-to-EI* model is able to simultaneously achieve both timescale localization and reliable signal propagation. As shown in Figure 1e, when the coupling strength increases, the downstream population exhibits a large magnitude of response while it maintains its characteristic timescale that is markedly distinct from the upstream population timescale. Further exploration shows that the model’s performance can be attainable across multiple parameter settings. As illustrated in Figure 1f, the response of the downstream excitatory population increases substantially as the inter-areal coupling strength increases, while the interference of upstream area’s timescale remains negligible. We term this desirable regime “*interference-free propagation*” (*IFP*).

In the IFP regime, a downstream brain area with a short characteristic timescale is even able to track fast oscillatory external inputs, despite that the inputs are received solely from an upstream area with a significantly slower timescale (Figure 1g). This “routing” mechanism effectively bypasses the temporal low-pass filtering imposed by the upstream area’s slower dynamics, ensuring the downstream area to integrate signals locally on its own distinct timescale, while simultaneously enabling robust inter-areal signal propagation. Specifically, the “routing” mechanism enables downstream areas to recover and track the high-frequency components of a driving signal that had been filtered out by the upstream node (detailed analysis and simulations provided in Figure S2 and Supplementary Texs S1).

### 2.3 Balanced Feedforward Excitation–Inhibition Implements IFP

To understand how the *E-to-EI* model achieves the IFP regime, we performed an analytical reduction of the model equations (see Methods and Supplementary Text S1). In brief, using a timescale-separation analysis, we approximated the *E-to-EI* model by an effective one-dimensional equation describing downstream excitatory population activity 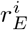. This reduced equation includes an extra term proportional to the time derivative of the upstream activity 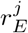, revealing that the downstream excitatory population is driven not only by the upstream firing rate itself (as in the *E-to-E* model), but also by the change in the upstream rate. These two input drives are important for achieving both signal propagation and timescale localization (see details in Methods). The accuracy of the model reduction is also validated numerically in Figure S3.

After model reduction, two *quantitative measures* emerged in the effective model that govern the downstream areal dynamics: *M*_*SP*_ and *M*_*T L*_ (definition in Methods). *M*_*SP*_ quantifies the effective strength of the mean feedforward drive from the upstream to the downstream area, while *M*_*T L*_ quantifies the residual influence of the temporal fluctuations of the upstream area on the downstream area after accounting for the external input into the downstream area. Our theory predicted that the magnitude of the downstream response scales with *M*_*SP*_ and that the degree of interference on the upstream time scale scales with *M*_*T L*_ (detailed analysis provided in Supplementary Text S1). Numerical simulations confirmed these predictions: the peak firing rate of the downstream excitatory population increases with *M*_*SP*_ , and the interference metric (reflecting the influence of upstream area’s timescale on the downstream area’s timescale) increases with *M*_*T L*_ (Figures 2a-b). Consequently, the IFP regime emerged in a parameter range yielding sufficiently large *M*_*SP*_ together with near-zero *M*_*T L*_.

**Fig. 2.**
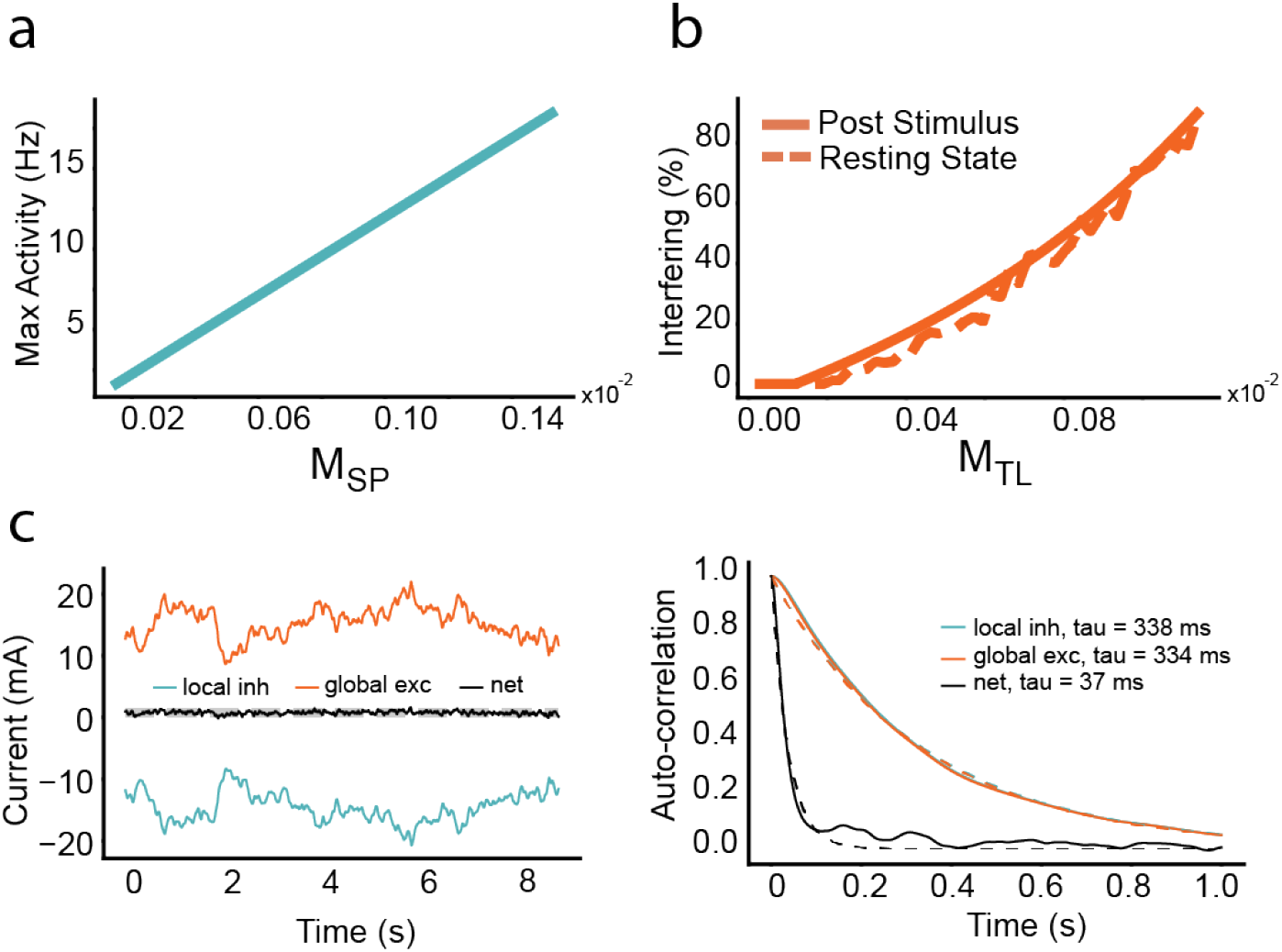
Mechanism of IFP in the *E-to-EI model*. (a) Correlation between the peak downstream activity and *M*_*SP*_ . (b) Correlation between the interference metric and *M*_*T L*_, with interference measured from downstream activity after a pulse (solid line) or during resting state driven by white noise (dashed line). (c) Left: In the IFP regime, input current to the downstream excitatory population from the upstream excitatory population (red) and the local inhibitory population (blue) when the upstream is driven by a stationary Gaussian input. The net current (black) is accurately predicted by 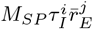 (gray dashed). Right: Auto-correlation functions of the upstream excitatory, local inhibitory, and net currents (solid) and their exponential fits (dash). While the E and I currents show slow timescale induced by the upstream activity, these slow components are canceled in the net current.

To build an intuitive understanding of the IFP mechanism, we examined the two pathways through which the upstream drives the downstream excitatory population in the *E-to-EI* model: a direct excitatory pathway from the upstream excitatory neurons to the downstream excitatory neurons, and an indirect inhibitory pathway from the upstream excitatory neurons to the downstream inhibitory neurons, which in turn inhibit the downstream excitatory neurons. By decomposing the synaptic currents to the downstream excitatory population from both pathways into their mean (steady) components and fluctuating (time-varying) components, we found that the mean component is proportional to *M*_*SP*_ , whereas the fluctuating component related the upstream area’s timescale is proportional to *M*_*T L*_ (see derivation in Methods).

This analysis further indicates that, in the IFP regime, the fluctuating synaptic current involving upstream areas’ timescales from the indirect inhibitory pathway nearly cancels out with that from the direct excitatory pathway, yielding a small *M*_*T L*_; while the mean synaptic currents from the two pathways remain substantially different, yielding a large *M*_*SP*_ . This temporal balance of feedforward excitation and feedforward inhibition enables reliable transmission of the mean signal while filtering out the slow temporal variations of the upstream activity. Figure 2c provides a concrete illustration of this mechanism: in the IFP regime, the upstream-driven excitatory current (red) and the local inhibitory current (blue) are of opposite sign and comparable magnitude in their fluctuating components (with similar intrinsic timescale around 335 ms).

This results in a non-vanishing net current with slow fluctuating components nearly canceled (black), showing its intrinsic timescale of 37 ms. In summary, by tuning the network parameters to achieve large *M*_*SP*_ and minimal *M*_*T L*_, the *E-to-EI* model successfully solves the trade-off present in the *E-to-E* model: it effectively propagates the averaged upstream areal activity while filtering out the upstream area’s intrinsic timescale.

### 2.4 IFP Exists across Diverse Network Motifs

Having established interference-free propagation (IFP) in the minimal two-area circuit, we next investigated whether this mechanism generalizes to more biologically realistic network motifs. Figure 3a illustrates a generalized two-area motif consisting of an upstream excitatory population projecting to both excitatory and inhibitory populations in a downstream area. Unlike the minimal two-area circuit, the motif explicitly incorporates local recurrent connections (E to E, E to I, I to E, and I to I) within each area, while retaining a single directed long-range input pathway from upstream E to downstream E and I (see Methods). This motif is common in cortical networks, providing a testbed to assess whether IFP represents a general cortical network mechanism or merely a special feature of the minimal model.

**Fig. 3.**
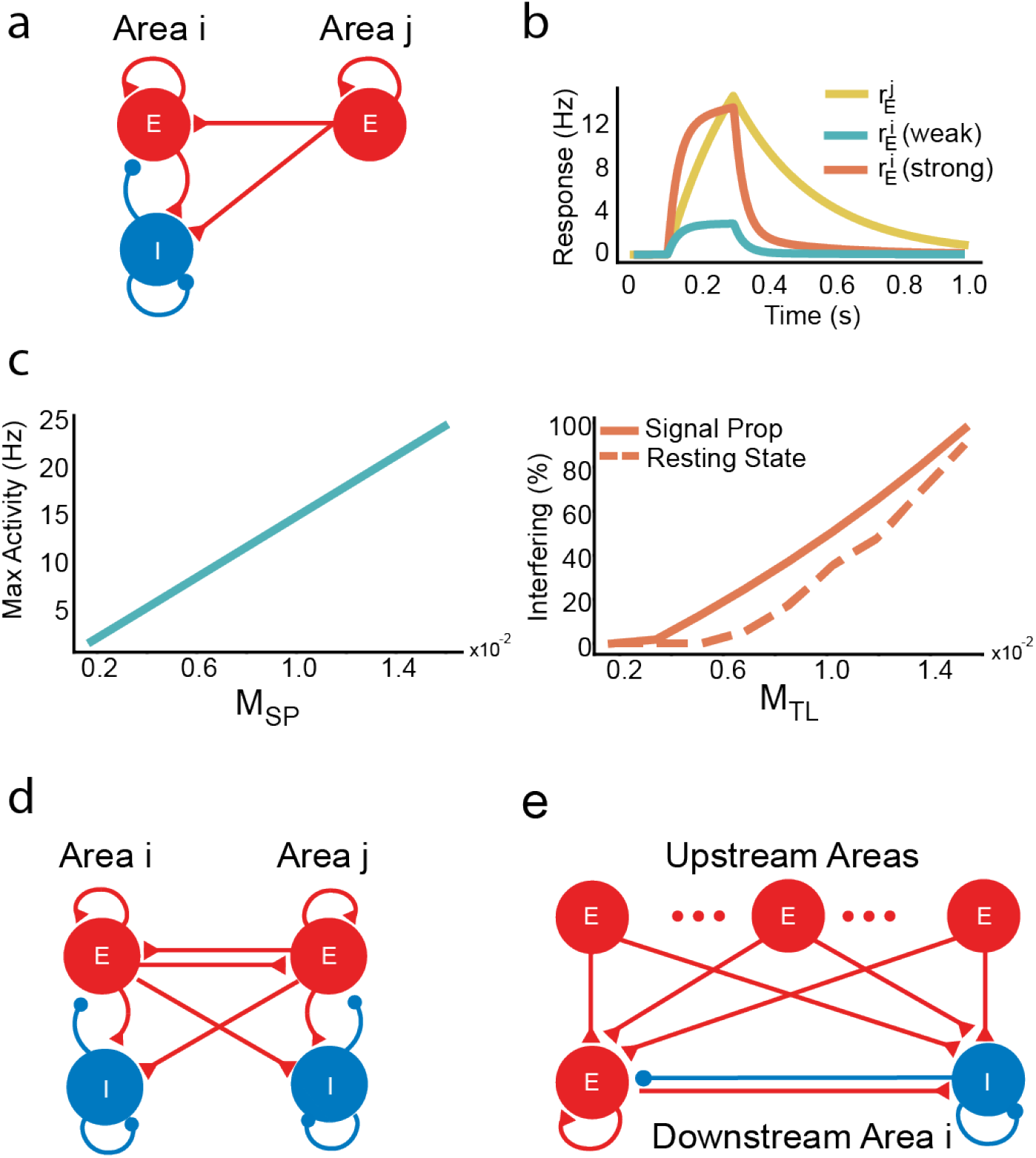
Dynamics of the *E-to-EI model with local recurrent connections*. (a) Scheme of the *E-to-EI* model with local recurrent connections, with the directed connection from area *j* to area *I* shown. (b) Response of the *E-to-EI* model with local recurrent connections to an external pulse input. Yellow: firing rate of upstream population 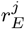. Blue and red: firing rates of downstream population 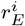 under weak and strong connection strengths. (c) Left panel: Correlation between the max firing rate and *M*_*SP*_ , illustrating its effectiveness as a metric for signal propagation. Right panel: Correlation between the interference metric and *M*_*T L*_, illustrating its effectiveness as a metric for timescale localization. (d) Scheme of the *E-to-EI model with reciprocal connections*. (e) Scheme of the *E-to-EI model with multiple upstream areas*.

Consistent with our findings from the minimal model, a pure E to E feedforward pathway with local E recurrence suffered the same trade-off: weak inter-area coupling preserved timescale localization but yielded poor signal propagation, whereas strong coupling improved signal propagation at the expense of downstream area inheriting the upstream area’s timescale (Figure S4a-c). In contrast, introducing a local inhibitory population downstream resolved this trade-off: as inter-areal coupling increases, the downstream response grows substantially while its intrinsic timescale remains localized (Figure 3b). In addition, fast periodic inputs can be faithfully tracked despite slow upstream dynamics (Figure S4d). Using a similar asymptotic reduction as in the minimal model (see Methods and Supplementary Text S2), we again derived an effective one-dimensional first-order description for the downstream area’s dynamics. The analysis also identified the generalized versions of the two key metrics, *M*_*SP*_ and *M*_*T L*_ (details in Supplementary Text S2). We then numerically confirmed that the peak downstream response scales with *M*_*SP*_ , while interference from the upstream timescale scales with *M*_*T L*_ (Figure 3c). Therefore, the IFP regime with a large *M*_*SP*_ and a vanishing *M*_*T L*_ remains exist in this network motif.

To further probe the generality of IFP, we examined two more biologically realistic motifs: a bidirectionally connected two-area network and a convergent hub motif with multiple upstream inputs converging onto a single downstream area (details in Supplementary Text S3). In the reciprocal two-area motif, both areas could maintain their own intrinsic timescale even under strong coupling, and effectively exchanged persistent signals without homogenizing their dynamics (Figure 3d, Figure S5). To extend our analytical framework to this bidirectional scenario, we defined separate direction-specific *M*_*SP*_ and *M*_*T L*_, which accurately predicted the magnitude of interareal firing-rate and the timescale interference in each direction (Figure S5). Similarly, in the convergent hub motif, we found that the downstream area could preserve its local timescale while integrating the signals from multiple upstream areas, each with distinct timescales (Figure 3e, Figure S6). Increasing the number of upstream inputs produced a near-linear increase in downstream firing-rate response (reflecting an additive increase in *M*_*SP*_), whereas the upstream timescale interference was effectively canceled out as long as each individual input operated within the IFP regime (each had a low *M*_*T L*_; Figure S6).

Together, these findings underscore the theoretical robustness of IFP across a variety of cortical network motifs, suggesting that IFP can preserve distinct timescales even in complex networks involving strong reciprocal coupling and convergent inputs. These motifs and the associated generalized metrics (*M*_*SP*_ , *M*_*T L*_) help bridge the gap between the minimal *E-to-EI model* and large-scale cortical network models, as a network model can be decomposed into these basic motifs in general.

### 2.5 Connectome-based Macaque Cortex Model Achieves IFP

We next examined the IFP regime in an anatomically constrained large-scale model of the macaque cortex with multiple areas. The model was adapted from our previous study [3], which includes 29 sensory and association areas of the macaque cortex (see Methods and Supplementary Text S4). In the model, each cortical area is represented by a local excitatory–inhibitory circuit (analogous to the *E-to-EI* model), and inter-area connections follow the empirically measured directed weights (Figure 4a). We further generalized the previously introduced metrics *M*_*T L*_ and *M*_*SP*_ to quantify timescale localization and signal propagation across multiple areas. By systematically varying the overall strength of inter-area coupling (*μ*_*EE*_, *μ*_*IE*_), we explored distinct dynamical regimes with different degrees of timescale localization and signal propagation.

**Fig. 4.**
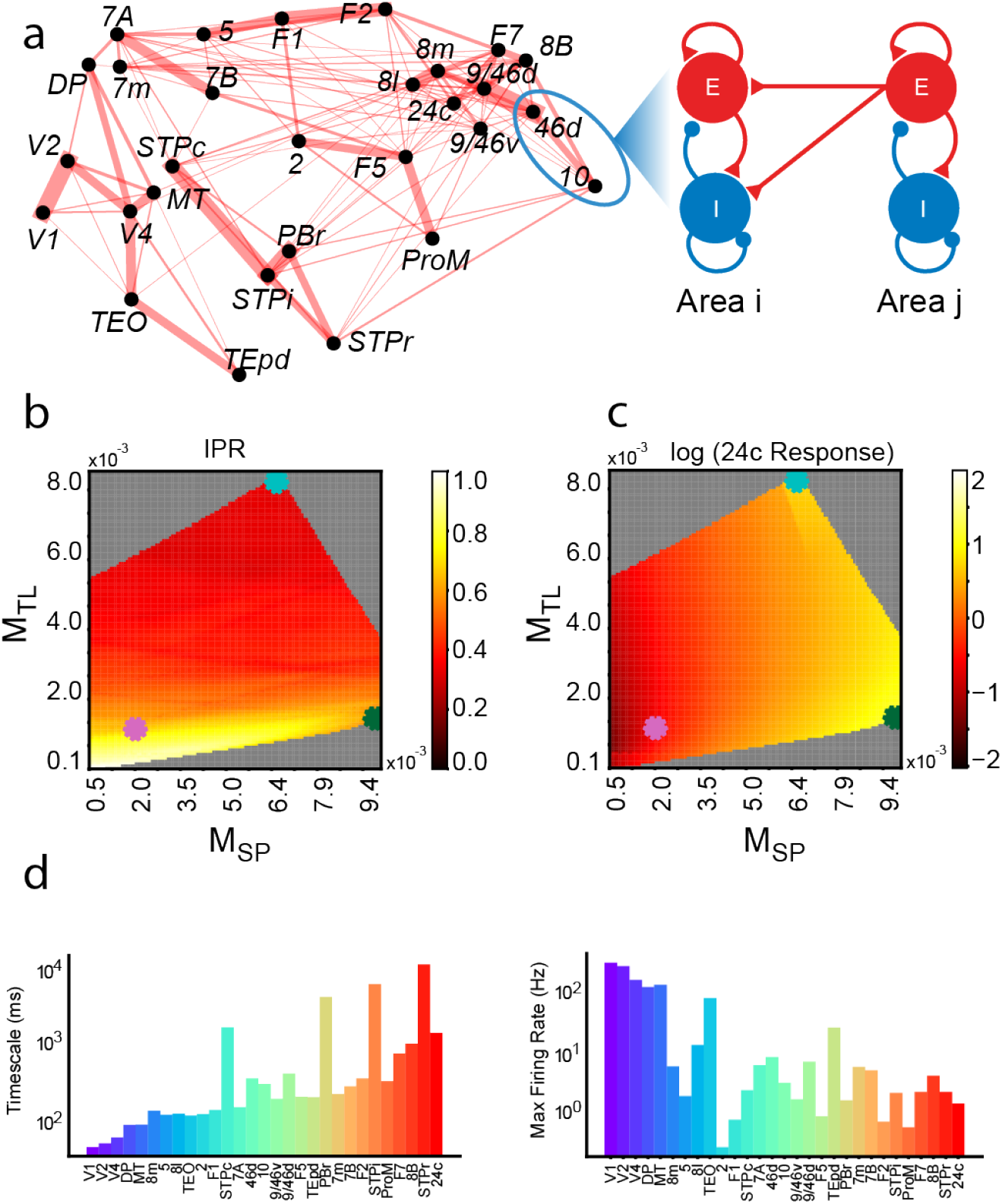
Dynamical regimes of the multi-areal macaque cortex rate model. (a) Left: Graph of macaque cortical areas, with line width representing connection strength (top 25% shown). Right: Model node for each cortical area, consisting of excitatory and inhibitory populations with directed interactions from area *j* to area *i* shown. (b) Heatmap showing timescale localization dependence on *M*_*T L*_ and *M*_*SP*_ , indicated by inverse participation ratio (IPR). (c) Heatmap showing signal propagation dependence on *M*_*T L*_ and *M*_*SP*_ , based on area 24c’s response. Gray areas indicate instability. Parameter values from Chaudhuri et al.[3], Joglekar et al.[31], and the IFP regime are marked by purple, blue, and green dots, respectively. (d) Dynamics in the IFP regime. Left, timescale of each area derived from the auto-correlation function of the simulated resting-state activity. Right, Peak firing rate across areas following a pulse stimulus to V1.

To characterize these dynamical regimes quantitatively, we introduced two summary indicators. For timescale localization, we used Inverse Participation Ratio (IPR) that measures the localization of an eigenvector associated with a particular timescale of the system (see definition in Methods). A localized eigenvector suggests that the corresponding timescale engages only a limited subset of brain areas (substantially nonzero elements in the eigenvector), indicative of localized timescale. For signal propagation, we gave input to the primary visual cortex (V1) which takes the lowest position in the hierarchy in the network, and measured the peak response of area 24c, the area with the highest hierarchical position in the network. Figures 4b and 4c depict phase planes that clearly illustrate the ability of *M*_*SP*_ and *M*_*T L*_ in quantifying timescale localization and signal propagation in the network. Notably, a decrease in *M*_*T L*_ leads to an increased IPR, signifying enhanced timescale localization, regardless of the *M*_*SP*_ value. In contrast, an elevation in *M*_*SP*_ correlates with a heightened response in area 24c, indicating improved signal propagation nearly independent of *M*_*T L*_.

Our findings provided new insights on earlier studies by Chaudhuri et al. [3], and Joglekar et al. [31] (Figure 4b, c and Figure S7). The former observed significant timescale localization and hierarchy but poor signal propagation in a specific regime of detailed balance, resonating with our findings with small *M*_*T L*_ and *M*_*SP*_ (purple star in Figure 4b, c). In contrast, Joglekar et al. [31] spotlighted effective signal propagation in another regime of balanced amplification, mirroring our observations with large *M*_*SP*_ and *M*_*T L*_ that signifies effective signal propagation but weak timescale localization (blue star in Figure 4b, c).

Importantly, our results identified a novel dynamical regime in the phase plane, diverging from earlier studies. As depicted in Figures 4b and 4c, a unique parameter regime around the green star marker is characterized by a large value of *M*_*SP*_ and a small value of *M*_*T L*_, which points to the IFP regime we previously identified. This novel regime uniquely achieves both robust signal propagation (large peak response of highlevel cortical areas) and strong timescale localization (high IPR index). The left panel of Figure 4d shows the characteristic timescale of each brain area in the IFP regime, confirming a clear hierarchy from fast (sensory) to slow (association) areas. The right panel of Figure 4d shows the peak firing rate in each area following a brief stimulus to V1, demonstrating that the stimulus evokes a substantial response in both sensory and high-level association areas (e.g., area 24c). In fact, the IFP regime combines the favorable aspects of the previous two regimes (Figure S8): Similar to Chaudhuri et al. [3] it maintains timescale localization across multiple brain areas, and similar to Joglekar et al. [31] it allows effective signal transmission across brain areas.

### 2.6 Biophysically Detailed Spiking Network Reproduces IFP

We further assessed whether the IFP regime identified in the rate model could be realized in a biologically detailed spiking neuron network. We constructed a multi-area spiking neural network model aligned with the previously described rate model (see Methods; details in Supplementary Text S5). Each cortical area in the spiking model is composed of excitatory and inhibitory leaky integrate-and-fire neurons connected via biologically realistic synapses of AMPA, NMDA, and GABA_*A*_ types, and areas are connected according to the empirically measured macaque connectome. After tuning model parameters to be in the IFP regime predicted by the rate model, we stimulated the network and analyzed its dynamics. Figure 5a presents the hierarchy of timescales across cortical areas (left), and peak firing rate responses to visual stimulation applied to V1 (right). Remarkably, the spiking network qualitatively replicated the key behavior observed in the rate model: each area maintains its own intrinsic timescale while signals propagate reliably through the network. Further quantitative analysis (Figure S9) confirms that the intrinsic timescales and peak firing rates are closely correlated with those in the rate model.

**Fig. 5.**
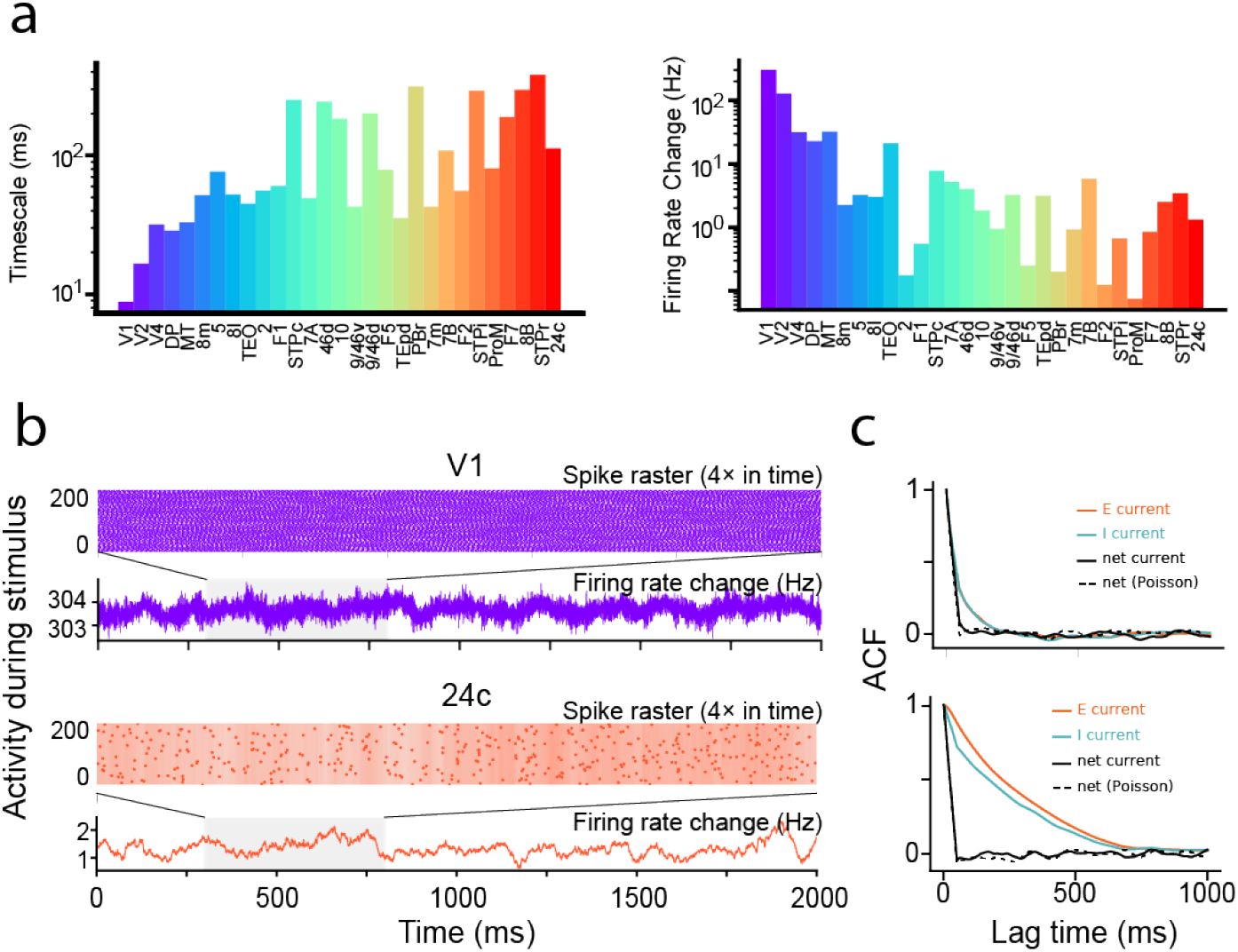
The IFP regime in the multi-areal spiking network model of macaque cortex. (a) Left, the timescale of each area derived from the auto-correlation function of activity during stimulus to V1. Right, Peak firing rate across different brain areas following stimulus to V1. (b) The spike raster and population firing rate change of two areas (V1 and 24c) during stimulus to V1. Their activities exhibit different timescales. (c) Autocorrelation functions for inter-areal excitatory current (red), intra-areal inhibitory current (blue), and net current (solid black) of V1 and 24c, respectively. The dashed black line represents the reference net current with Poisson input replacing inter-areal inputs. Overlap of solid and dashed black curves indicates temporal cancellation of E and I currents.

To better illustrate the network dynamics underlying the IFP regime, Figure 5b shows representative activity from a lower-level visual area (V1) and a higher-level prefrontal area (24c) during stimulation delivered to V1. Spike rasters (upper panels) and population firing rates (lower panels) for V1 (purple) and area 24c (red) reveal distinct intrinsic dynamics: V1 activity fluctuates on a fast timescale (tens of ms) whereas area 24c activity evolves on a much slower timescale (hundreds of ms), consistent with their positions in the hierarchy. Nevertheless, stimulation of V1 robustly drives activity in area 24c (with firing rate increases of ∼ 2 Hz, comparable to the rate model), indicating effective signal propagation from V1 to area 24c.

We then asked whether the same mechanism of temporal cancellation identified in the rate model underlies the IFP regime in this detailed spiking network. Figure 5c plots autocorrelation functions of different synaptic input currents to two representative areas: V1 and area 24c (and more in Figure S10). In both V1 (top panel) and area 24c (bottom panel), the total inter-areal excitatory input current (red curves) exhibits slow temporal correlations, which are mirrored by local intra-areal inhibitory currents (blue curves). However, the net synaptic input current (black solid curves), representing the sum of excitatory and inhibitory inputs, shows substantially reduced temporal correlations, closely matching the reference case where upstream inputs are replaced by temporally uncorrelated Poisson spikes (black dashed curves).

These results suggest that in the spiking network, slow temporal fluctuations of excitatory inputs from other areas are systematically canceled out by locally generated inhibitory currents, leaving only rapid fluctuations in the net current. This cancellation mechanism precisely parallels the one identified in the rate model. Thus, introducing realistic spiking dynamics and synaptic conductances does not abolish the IFP phenomenon. These findings demonstrate the broad applicability of the IFP across different levels of biological detail, strongly supporting its potential as general operating principle of cortical networks, whereby a hierarchical organization of intrinsic timescales can coexist with strong inter-areal communication.

### 2.7 Experimental Evidence of IFP in the Marmoset Cortex

To investigate whether the IFP mechanism exists in real cortical networks, we turned to the marmoset, another primate species with both a well-mapped cortical connectome [39, 44] and publicly available multi-region ECoG datasets [45, 46]. While multi-area recordings in macaques remain limited, the marmoset data offered an ideal testbed for validating our model’s predictions. We constructed a large-scale rate model of the marmoset cortex analogous to our macaque model, incorporating empirical anatomical connectivity and an area-specific gradient of excitation similar to that used in the macaque model (see Methods). Model parameters were calibrated to align each area’s simulated intrinsic timescale with empirical ECoG-derived timescales.

We first evaluated whether the model could reproduce the timescale hierarchy observed experimentally. By simulating a resting state condition (uniform white-noise background input to all areas), our model successfully reproduced the experimentally observed hierarchy of intrinsic timescales across different cortical areas. As shown in Figure 6a-c, lower-level areas resonate at higher frequencies, closely matching the input, while higher-level areas exhibit smoother activity, indicating their longer timescales. The autocorrelation function of each area’s areas revealed exponential decays with significant variability in decay constants, confirming distinct local timescales (Figure 6b). Quantitatively, model-derived timescale closely matched the experimentally measured ECoG timescale (Figure 6c). We note that the absolute values of model-derived timescales were scaled 5-fold larger than the ECoG-derived timescales, consistent with experiment findings [19].

**Fig. 6.**
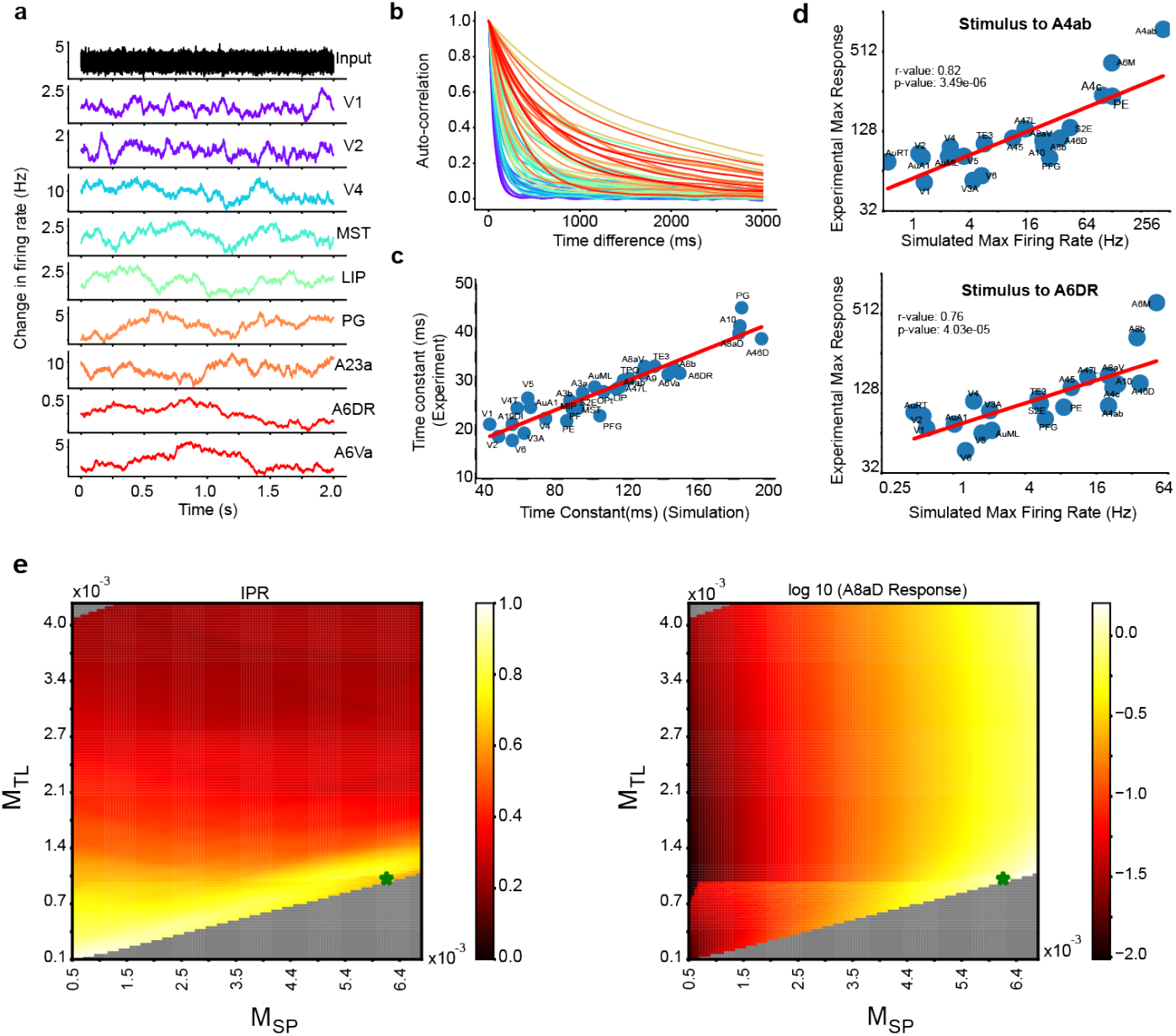
The marmoset model in the IFP regime replicates experiment observations. (a) Simulated resting state activity, with white-noise input applied to all brain areas. Lower-level areas fluctuate at higher frequencies, whereas higher-level areas show more gradual fluctuations. (b) The autocorrelation function of each area’s activity in the resting state. (c) Strong correlation between the timescale of simulated resting-state activity and experimental data (*r* = 0.94, *p* = 6.57 *×* 10^*−*16^). (d) Comparison of the model’s peak response with optogenetic experimental data when stimulating brain areas A4ab (top panel) or A6DR (bottom panel). (e) Evidence that the model resides in the IFP regime. Left panel: Heatmap illustrating the impact of *M*_*T L*_ and *M*_*SP*_ on timescale localization, as indicated by IPR. Right panel: Heatmap illustrating the impact of *M*_*T L*_ and *M*_*SP*_ on signal propagation, measured by the response in area A8aD with the highest hierarchical value. The parameter values of the model determined by experimental data are marked with the green star.

Next, we tested whether the model could capture inter-areal signal propagation patterns observed experimentally. We utilized data from prior optogenetic experiments in which brief LED-driven optogenetic stimulation was applied to specific marmoset cortical areas (e.g., A4ab or A6DR), and resulting activity was measured via multiregion ECoG recordings distributed across the marmoset cortex [45]. Mimicking this stimulation protocol in silico, we applied a 200 ms pulse current input to the corresponding model area and quantified the peak firing rate response in other cortical areas. As shown in Figure 6d, the overall pattern of inter-area signal propagation in our model closely aligns with the experimental data across most cortical areas.

Finally, we assessed whether the experimentally constrained marmoset model operates within the predicted IFP regime defined by our theoretical metrics for signal propagation (*M*_*SP*_) and timescale localization (*M*_*T L*_). Using the same *M*_*SP*_ - *M*_*T L*_ phase plane analysis previously applied to the macaque model (Figure 4), we found a consistent relationship between these metrics and model dynamics: a decrease in *M*_*T L*_ leads to an increased IPR, signaling enhanced timescale localization (independent of *M*_*SP*_), while an elevation in *M*_*SP*_ correlates with a heightened response in area A8aD that has the highest hierarchical value in the network, indicating improved signal propagation (Figure 6e). Importantly, the parameters derived by fitting the marmoset model to experimental data correspond to a point of large *M*_*SP*_ and small *M*_*T L*_ in the phase plane, with values comparable to those identified in the macaque model. These values (green star in Figure 6e) fall within the IFP regime, indicating that the experimentally constrained marmoset model implements the IFP mechanism.

## 3 Discussion

In summary, our study demonstrates that robust inter-areal signal propagation can coexist with area-specific intrinsic dynamics within a novel dynamical regime we term **interference-free propagation (IFP)**. In this IFP regime, cortical networks reliably transmit activity from one area to another without disrupting or homogenizing the characteristic timescale of each area. Each brain area preserves its unique temporal integration properties, from fast, transient responses in early sensory areas to slow, integrative dynamics in higher association areas, even as signals successfully propagate through the cortical hierarchy. These findings provide a new perspective on cortical operation, showing how specialized local temporal processing and effective global communication can simultaneously be achieved within biologically realistic networks.

We identified across multiple levels of analysis that a precisely balanced feedforward excitatory-inhibitory circuit underlies IFP in cortical networks. In minimal two-area circuits, we found that introducing a local inhibitory population (the *E-to-EI* motif) was essential for maintaining the downstream area’s intrinsic timescale despite receiving strong feedforward input from the upstream area. Analytically, we developed two metrics, *M*_*SP*_ and *M*_*T L*_, to quantify the effective strength of signal propagation and the degree of residual upstream temporal interference, respectively. Extending our findings, we verified that IFP generalizes robustly to more biologically plausible and complex network motifs, including local recurrent connections, reciprocal (bidirectional) connectivity, and multiple upstream inputs. Furthermore, in an anatomically constrained large-scale macaque cortical model and the corresponding biophysically detailed spiking network, we demonstrated that networks naturally operate within the IFP regime, preserving a hierarchical timescale structure alongside reliable inter-area signal propagation. Finally, using a data-constrained marmoset cortical model that reproduces empirical ECoG-derived intrinsic timescales and neural activity patterns, we found that the fitted network operates in the IFP regime, suggesting that real cortical networks may indeed exploit this mechanism.

Mechanistically, interference-free propagation works via a precise temporal balance of feedforward excitation and inhibition, intuitively separating an input signal into mean and fluctuating components. By tuning the feedforward excitatory and inhibitory inputs from an upstream area, the downstream area receives a strong drive mainly from the upstream’s mean activity, while any fluctuations related to the upstream area’s timescale are canceled out. Through this selective transmission, each downstream area gains a strong incoming signal without inheriting the upstream area’s temporal interference. Consequently, cortical areas can interact robustly without homogenizing their temporal dynamics, providing a new mechanistic explanation for how the brain achieves both efficient inter-area communication and specialized local temporal processing.

Critically, the IFP regime bridges a gap between previous large-scale models of cortical dynamics. Earlier models successfully captured the empirically observed hierarchy of intrinsic timescales across cortical areas, but struggled to transmit activity through the hierarchy (e.g., stimulus-driven activity dissipated before reaching higherorder areas; see Chaudhuri et al. [3]). Conversely, subsequent models introduced by Joglekar et al. [31] demonstrated that strong inter-areal excitation balanced by local inhibition can enable reliable signal propagation across cortex, but at the cost of homogenizing network dynamics, effectively losing the distinct intrinsic timescales. The IFP regime identified in our study combines the strengths of these prior studies, preserving pronounced local-timescale heterogeneity while allowing stable long-range transmission of signals. In the *M*_*SP*_ −*M*_*T L*_ phase plane summarizing this trade-off (Figure 4b–c), IFP occupies a previously unexplored region where both propagation efficacy and timescale localization are high, whereas the dynamical regimes studied in previous models were constrained to achieve either of the properties. Thus, our findings resolved a longstanding trade-off between areal dynamical specialization and inter-areal communication in cortical network modeling, revealing conditions under which the cortex can accomplish both.

Our study revisits and extends the concept of “balanced amplification” initially proposed by Murphy and Miller [43] that describes the amplification of neural activity patterns in cortical circuits, and later generalized to “global balanced amplification” by Joglekar et al. [31]. Central to both the foundational work of Murphy and Miller [43] and our IFP mechanism is the modulation of signal amplification through a finely tuned balance between excitatory and inhibitory inputs, decoupling it from the inherent timescales of neural activity. This mechanism cannot be achieved without inhibitory population, emphasizing the important role of inhibitory neurons in maintaining the intrinsic timescale of an area.

However, our work sets itself apart by studying anatomically based multi-areal cortical models that explicitly incorporate areal heterogeneity. Each cortical area in the model possesses its own intrinsic timescale. This contrasts with traditional cortical network models, which typically assume homogeneous neuronal properties across areas. Incorporating areal specificity introduces additional complexity in maintaining the functional individuality of cortical areas operating across diverse timescales while ensuring effective inter-area signal propagation. Furthermore, we have developed a novel analytical framework applicable to general neural networks consisting of both excitatory and inhibitory groups. This analytical framework enables us to derive novel quantitative metrics to rigorously characterize and analyze timescale localization and signal propagation (*M*_*T L*_, *M*_*SP*_), which we then apply to the multi-areal models. This unifying analytical framework also provides new insights into the operational conditions required for “balanced amplification”, as detailed further in Supplementary Text S1.

Our investigation underscores the critical role of long-range cortical connections in shaping signal propagation and maintaining area-specific intrinsic dynamics within the cortex. As our current approach primarily assesses average connection strengths between areas, it does not yet explicitly explore the intricate details of connectomic patterns, such as clustering, motifs, or the effects of the exponential distance rule on long-range projections [47]. The analytical and computational methods developed in this study provide promising tools for future exploration of how detailed connectomic structures influence cortical network dynamics. Investigating how specific connection patterns impact the signal propagation and timescale localization represents an important and exciting avenue for future research, potentially revealing additional biological mechanisms that cortical networks use to flexibly manage communication across the hierarchy.

## 4 Methods

### 4.1 The two-area E-to-E model

We started from a motif in which an excitatory neuronal group in one brain area sends input to another excitatory group in a different area. This model, referred to as the *E to E model*, serves as a baseline for exploring inter-area interactions. The model is mathematically formulated as follows:

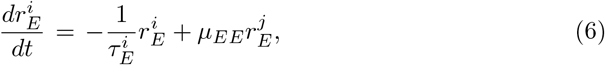

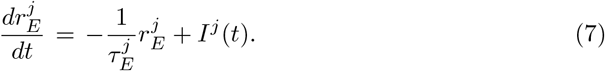

In this formulation, 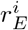 and 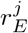 denote the firing rates of excitatory neuron populations in regions *i* and *j*, respectively. The parameter 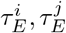 represents the effective time constant of firing rate of neuron population *i* and *j*, respectively, and *μ*_*EE*_ defines the coupling strength of the inter-areal excitatory input from region *j* to region *i*. The 18 external inputs to area *j* is represented as *I*^*j*^(*t*). We set 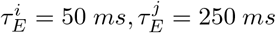. The coupling strength *μ*_*EE*_ is varied depending on the scenarios. For Figure 1b, we choose *μ*_*EE*_ = 0.0005*/ms* for weak connection and *μ*_*EE*_ = 0.02*/ms* for strong connection. For Figure 1c, *μ*_*EE*_ is varied between 0 to 0.02*/ms*.

### 4.2 The two-area E-to-EI model and its asymptotic reductions

Based on the *E-to-E* model described above, we introduced an inhibitory neuron group alongside the excitatory group in area *i*, while area *j* remains exclusively excitatory. The directed connection from area *j* to area *i* contains excitatory projections from area *j* to both excitatory and inhibitory neuron groups in area *i*. The dynamics of this extended model are described by the following equations:

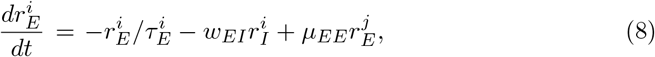

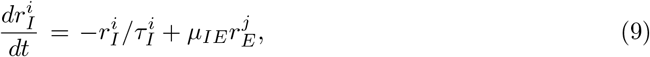

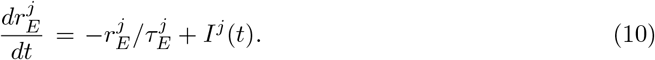

where 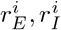 are the firing rates of excitatory and inhibitory groups in area *i*, respectively and 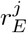 is the firing rate of excitatory group in area *j*. The parameter 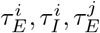 represent the effective time constant for firing rate of excitatory and inhibitory population in the area *i* and *j*, respectively. *w*_*EI*_ denotes the coupling strength from the inhibitory population to the excitatory population within the area *i* and *μ*_*XE*_ represents the coupling strength of the inter-areal input from the excitatory population to the *X* population in a downstream cortical area. Other parameters retain their definitions from the *E-to-E* model. We set 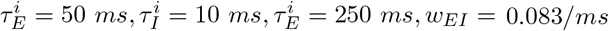. For Figure 1e, we set *μ*_*EE*_ = 0.031*/ms, μ*_*IE*_ = 0.035*/ms* for weak connection and *μ*_*EE*_ = 0.310*/ms, μ*_*IE*_ = 0.355*/ms* for strong connection (also for Figure 1g). For Figure 1f, the *μ*_*EE*_ is varied between 0*/ms* and 0.763*/ms* and *μ*_*IE*_ changes from 0*/ms* to 0.882*/ms*, correspondingly.

In the limit of a large separation between the *E* and *I* timescales, we obtained an approximate first-order system for 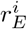 and 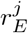. In particular, the downstream dynamics can be written in the form:

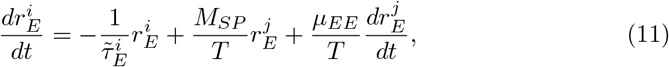

Where 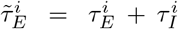 is the effective local timescale, 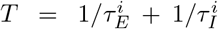 and 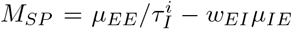. Equation (11) shows that the upstream’s rate of change 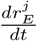 contributes to driving the downstream activity (a direct consequence of the feedforward inhibitory pathway). We then incorporated the upstream population’s dynamic equation (Eq. (10)) into the above. Substituting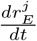from Equation (10) and rearranging terms, we obtained:

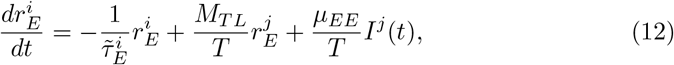

where 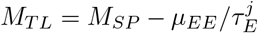is the metric for timescale localization. Equation (11)-(12) are the reduced two-area models used to analyze signal propagation (through the *M*_*SP*_ term) and timescale interference (through *M*_*T L*_). The main text discussion of *M*_*SP*_ and *M*_*T L*_ and their roles in IFP was based on these equations. All further mathematical details, including analyses of the reciprocal and multi-input extensions of the model, can be found in Supplementary Text S1-3.

### 4.3 The E-to-EI model incorporating local recurrent connection

To investigate the effect of local recurrence, we explicitly incorporated local recurrent connections (E to E, E to I, I to E, and I to I) within each area, while retaining a single directed long-range input pathway from upstream E to downstream E and I. The dynamics of the system can then be described as follows:

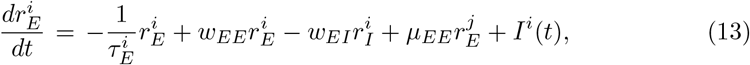

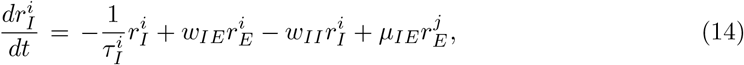

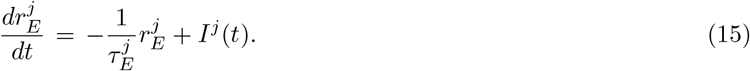

where 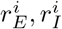 are the firing rates of excitatory and inhibitory groups in area *i*, respectively and 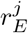 is the firing rate of excitatory group in area *j*, while 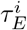 and 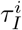 are the corresponding intrinsic time constants, respectively. *w*_*XY*_ symbolizes the coupling strength from *Y* population to *X* population within the area (*X, Y* could be E or I population) and *μ*_*XE*_ represents the coupling strength of the inter-areal input from the excitatory population to the *X* population in a downstream cortical area. Following a previous study [48], we set 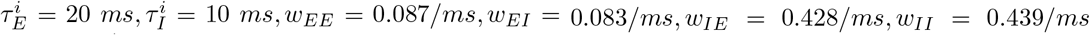, consistent with empirical findings. For Figure 3b, we set *μ*_*EE*_ = 0.400*/ms, μ*_*IE*_ = 2.571*/ms* for weak connection and *μ*_*EE*_ = 1.788*/ms, μ*_*IE*_ = 11.478*/ms* for strong connection (also for Figure S4d,e). For Figure 3c, the *μ*_*EE*_ is varied between 0.672*/ms* and 6.624*/ms* and *μ*_*IE*_ changes from 4.320*/ms* to 42.589*/ms*, correspondingly.

### 4.4 The multi-areal rate model of macaque neocortex

We built the multi-areal macaque cortical model that includes 29 cortical sensory and association areas, adapted from previous works on the modeling of macaque cortical network [3, 31]. Each cortical area is modeled with an excitatory group and an inhibitory group, which are governed by the following dynamics.

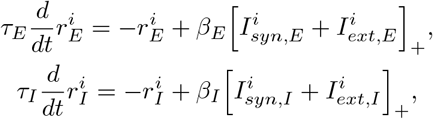

where 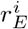 and 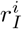 are the firing rates of the excitatory and inhibitory populations in the *i*th area, respectively, *τ*_*E*_ and *τ*_*I*_ are their intrinsic time constants, respectively, and *β*_*E*_ and *β*_*I*_ are the slopes of the f-I curves for the excitatory and inhibitory populations, respectively. The f-I curve takes the form of a rectified linear function with 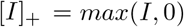. In addition, 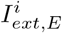and 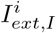 are the external currents, and 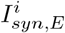 and 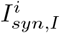 are the synaptic currents that follow

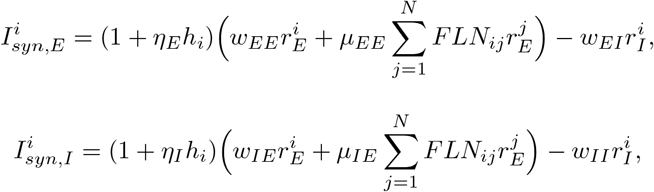

where *w*_*pq*_, *p, q* ∈ {*E, I*} is the local coupling strength from the *q* population to the *p* population within each area. *FLN*_*ij*_ is the fraction of labeled neurons (FLN) from area *j* to area *i* reflecting the strengths of long-range input [38], and *μ*_*EE*_ and *μ*_*IE*_ are scaling parameters that control the strengths of long-range input to the excitatory and inhibitory populations, respectively. Both local and long-range excitatory inputs to an area are scaled by its position in the hierarchy quantified by *h*_*i*_ (a value normalized between 0 and 1), based on the observation that the hierarchical position of an area highly correlates with the number of spines on pyramidal neurons in that area [3, 49]. Constants *η*_*E*_ and *η*_*I*_ map the hierarchy *h*_*i*_ into excitatory connection strengths for the E and I populations, respectively.

Adapted from the parameters chosen from a previous study [48], we set *τ*_*E*_ = 20 *ms, τ*_*I*_ = 10 *ms, β*_*E*_ = 0.066 *Hz/pA, β*_*I*_ = 0.351 *Hz/pA, η*_*E*_ = 0.685, *η*_*I*_ = 0.745, *w*_*EE*_ = 24.4 *pA/Hz, w*_*EI*_ = 19.7 *pA/Hz, w*_*IE*_ = 11.66 *pA/Hz, w*_*II*_ = 12.5 *pA/Hz, μ*_*EE*_ = 132.58 *pA/Hz, μ*_*IE*_ = 98.92 *pA/Hz*.

### 4.5 The multi-areal spiking network model of macaque neocortex

We constructed a large-scale spiking neural network of the macaque cortex comprising 29 cortical areas, following the same areal composition as the rate-based model described above. Each cortical area was modeled as a local excitatory–inhibitory circuit consisting of 800 excitatory (E) and 200 inhibitory (I) spiking neurons, implemented as leaky integrate-and-fire (LIF) units.

Within each area, neurons were sparsely and randomly connected with a 10% connection probability. Excitatory neurons project to both excitatory and inhibitory targets via glutamatergic synapses (AMPAR and NMDAR), while inhibitory neurons project locally to both neuron types via GABA_A_ synapses. Consistent with the rate model architecture, long-range (inter-areal) connections were exclusively excitatory: each area’s excitatory population sent projections to both the excitatory and inhibitory populations of other areas, with the projection strengths scaled according to empirical connectome (*FLN*). To incorporate hierarchical differences across areas, excitatory synaptic weights were modulated by the area-specific factor (1 + *ηh*_*i*_). Each neuron also received independent Poisson background input via AMPA synapses to mimic ongoing spontaneous cortical activity.

The membrane potential dynamics of individual neurons were governed by the LIF equations:

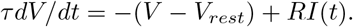

When the membrane potential reached the spike threshold, the neuron generated an action potential and its membrane potential was reset to the reset value, entering a brief refractory period during which no additional spikes could occur. Parameters were consistent across neurons: excitatory cells had a resting potential *V*_*rest*_ = −70*mV* , spike threshold –50 mV, reset potential –55 mV, membrane time constant 20 ms, and refractory period 2 ms. Inhibitory neurons had the same voltage parameters but a shorter membrane time constant (10 ms) and a shorter refractory period (1 ms).

Synaptic interactions were implemented as transient conductance changes with biologically realistic receptor kinetics. Excitatory synapses include both fast AMPAR and slow NMDAR components: AMPAR currents had a rapid decay time constant (2*ms*) with reversal potential 0*mV* , whereas NMDAR currents featured a voltage-dependent *Mg*^2+^ block and exhibited a 2*ms* rise time with a slow decay of 100*ms*, and with reversal potential 0*mV* . Inhibitory synapses are *GABA*_*A*_-mediated, characterized by a reversal potential of −70*mV* and a decay time constant of 5*ms*. Each presynaptic spike triggered an instantaneous conductance increase (delta-function) in synaptic conductance, which subsequently decayed with receptor-specific time constants. All synaptic weights for local and inter-area connections were fixed and tuned specifically to achieve the interference-free propagation (IFP) regime, with detailed parameter values provided in the Supplementary Materials.

We performed all simulations in Python using BrainPy 2.6.0 library [50] with a numerical integration timestep of 0.1*ms*. Each simulation consisted of a 5 seconds pre-stimulus period representing spontaneous activity (background Poisson inputs only), followed by a 150 seconds stimulus period during which excitatory neurons in area V1 received continuous Poisson spike trains at a rate of approximately 9 kHz to mimic prolonged visual input. The stimulation period was followed by a 5 seconds post-stimulus interval. The duration of 150 seconds ensured robust estimation of autocorrelation-based measures of intrinsic timescale. Population firing rates for each brain area were computed by convolving spike outputs with a 50*ms* sliding window. Intrinsic timescales of each area were estimated by fitting an exponential function to the autocorrelation of the area’s steady-state firing-rate response. All further modeling details can be found in Supplementary Text S5.

### 4.6 The multi-areal rate model of marmoset neocortex

We also built a multi-areal marmoset cortical model composed of 55 cortical areas spanning sensory to association areas. Each cortical area was represented by a pair of excitatory and inhibitory neuronal populations, governed by the same set of ordinary differential equations used previously for the macaque neocortex model. Importantly, we also incorporated an area-specific composite gradient factor, *h*_*i*_, which modulates both local intra-areal and long-range inter-areal excitatory inputs. The composite gradient *h*_*i*_ was fitted to match experimental observations of intrinsic timescale gradient across areas and structural hierarchy derived from the fraction of supragranular labeled neurons (SLN) reported by Theodoni et al. (2022) [39]. Details of the fitting are described in Li et al. (2025) [51]. Model parameters were initially adapted from experimental studies [48] and further fine-tuned to reproduce experimental observations, resulting in the following final parameters: *τ*_*E*_ = 20 *ms, τ*_*I*_ = 10 *ms, β*_*E*_ = 0.066 *Hz/pA, β*_*I*_ = 0.351 *Hz/pA, η*_*E*_ = 0.685, *η*_*I*_ = 0.76, *w*_*EE*_ = 24.4 *pA/Hz, w*_*EI*_ = 19.7 *pA/Hz, w*_*IE*_ = 11.66 *pA/Hz, w*_*II*_ = 12.5 *pA/Hz, μ*_*EE*_ = 67.4 *pA/Hz, μ*_*IE*_ = 49.81 *pA/Hz*.

### 4.7 The interference metric

To quantify timescale localization of the downstream area in the two-areal model, we first fit the downstream area’s response following a pulse stimulus, *f* (*t*), with a double exponential decay function *f* (*t*) = *A*_1_ exp(− *t/τ*_1_) + *A*_2_ exp(− *t/τ*_2_), where *τ*_1_ and *τ*_2_ represent the effective characteristic timescales of the upstream and downstream areas, respectively.

The interference metric is then computed as the ratio *A*_1_*/*(*A*_1_ + *A*_2_), which ranges from 0 (minimal interference) to 1 (maximal interference) when the upstream area has a slower intrinsic timescale than the downstream area. The metric can be negative for the other direction. Zero interference metric defines perfect timescale localization for the downstream area.

### 4.8 Inverse participation ratio

We use inverse participation ratio (IPR) to quantify timescale localization of the downstream area in the multi-areal rate model of the macaque and marmoset cortex. Given a normalized eigenvector **v** with unit length, IPR provides a measure of vector localization and is defined as follows:

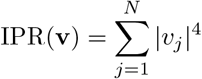

where *v*_*j*_ is the *j*th element of **v** and *N* is the dimensionality of the vector. To evaluate the IPR of the eigenvector matrix, we calculate the mean IPR value for each eigenvector (column) within the eigenmatrix.

### 4.9 Metrics of *M*_*SP*_ and *M*_*T L*_ in the net input current analysis

To provide an intuitive understanding of the IFP mechanism, we compute the input currents received by the downstream *E* population, from which the metrics *M*_*SP*_ and *M*_*T L*_ will naturally emerge. In the *E to E-I* model (Figure 1d), the upstream area influences the downstream excitatory population activity through two pathways: a direct excitation pathway where the upstream activity excites the downstream excitatory population, and an indirect inhibition pathway where the upstream activity first excites the downstream inhibitory population, which then inhibits the downstream excitatory population. By decomposing the neural activity and input into their mean and temporally varying components, i.e., 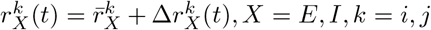, we express the excitatory input to the downstream excitatory population from the direct pathway as

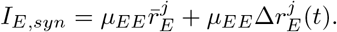

In addition, we calculate the inhibitory input to the downstream excitatory population from the indirect pathway as

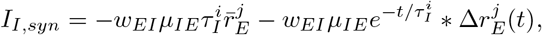

where ∗ denotes temporal convolution.

When the upstream area receives a stationary stochastic input, we have 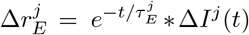, i.e., the time-varying upstream response to the fluctuating component of the input 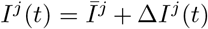. The total synaptic inputs *I*_*total,syn*_ = *I*_*E,syn*_ + *I*_*I,syn*_ takes the form as

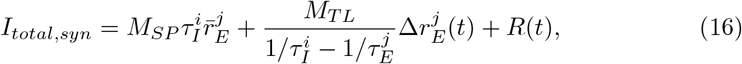

where the remainder term 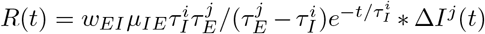 operates only with inhibitory population’s timescale 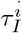, independent of upstream timescale.

As seen in equation (16), *M*_*SP*_ and *M*_*T L*_ appear in the mean and temporal fluctuating components of the total synaptic inputs, respectively. A large value of *M*_*SP*_ reflects a large mean current to the downstream excitatory population, which results from the imbalance between the mean of the excitatory inputs from the direct pathway and that of the inhibitory inputs from the indirect pathway. Therefore, *M*_*SP*_ quantifies the effectiveness of signal propagation through the mean signals. In addition, a small value of *M*_*T L*_ reflects a small fluctuating current component governed by the upstream activity timescale, which results from a substantial cancellation between the temporal fluctuations of the excitatory and inhibitory inputs from the two pathways. Therefore, *M*_*T L*_ quantifies the degree of timescale localization.

### 4.10 Generalized metrics for signal propagation and timescale localization

Expanding the metrics of *M*_*SP*_ and *M*_*T L*_ in the two-area case to a scenario with multiple areas, the formula for the two metrics can be generalized as:

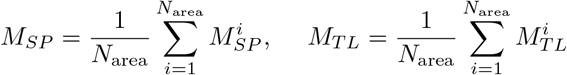

where *N*_area_ is the number of areas, and

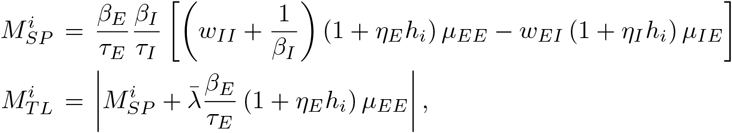

whereas 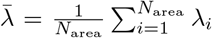, and *λ*_*i*_ is the eigenvalue corresponding to area *i* where all brain areas are isolated (i.e., the inverse of the intrinsic timescale of area *i*). The generalized metrics of *M*_*SP*_ and *M*_*T L*_ describe the average ability of signal propagation and timescale localization of a multi-areal network, respectively.

## Supplementary information

Figures S1-S15 Supplementary Figures

Text S1. The two-area models

Text S2. The two-area models with local recurrent connections

Text S3. Extensive Scenarios for the two-area models

Text S4. The multi-regional network-level model

Text S5. The multi-regional spiking neuron network model

## Acknowledgements

We thank H. Xu and H.P. Zhang for helpful discussions.

## Declarations

### Data availability

Data will be publicly accessible upon publication.

### Code availability

Code will be publicly accessible upon publication.

### Author contribution

Conceptualization, X-J.W., S.L.; methodology, G.L., Z.J and S.L.; investigation, G.L. and S.L.; writing, G.L., X-J.W., S.L.; funding acquisition, X-J.W., S.L.; supervision, X-J.W., S.L.

