## Supplemental Figures and Texts for "Interference-Free Propagation: Achieving Reliable Signal Propagation in Cortical Networks with Areal-Specific Local Dynamics"

#### 1 Supplementary Figures

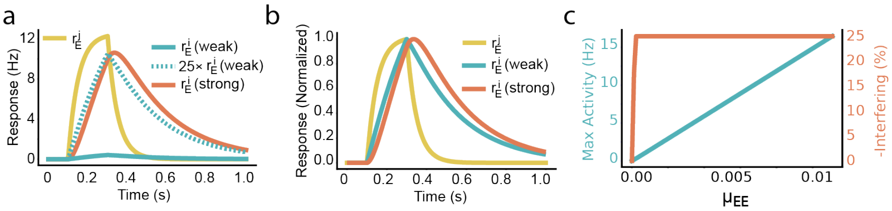


**Fig. S1 Numerical results for the E-E model when the timescale of the downstream node is slower than that of the upstream node.** (a)–(b) Response of the model to an external pulse input, presented unnormalized (a) and normalized (b). Yellow: the firing rate of the upstream population $r_E^j$; Blue and Red: the firing rates of the downstream population $r_E^i$ under weak and strong connection strengths. (c) Signal propagation and timescale interference in the E-E model at varying connection strengths. Blue: The peak firing rate of the downstream population. Red: the interference measure.

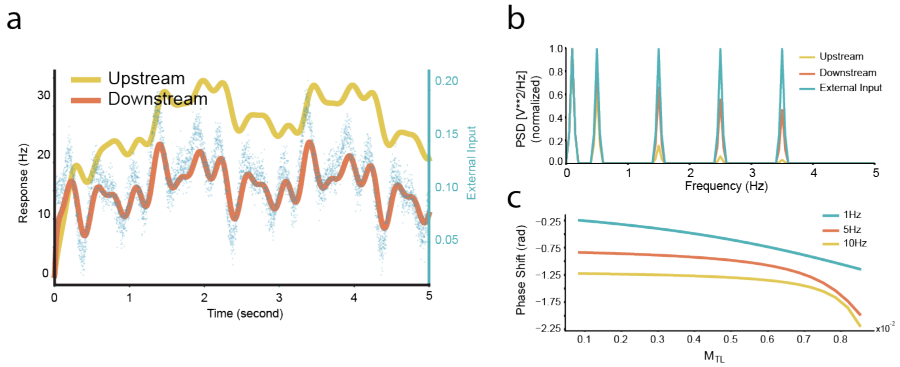


**Fig. S2 Results of the *simplified two-area model receiving oscillatory input.*** (a) Response of the simplified two-area model receiving oscillatory input. The external input (blue dot line) is composed of five sinusoidal functions with identical amplitudes but different frequencies and phases, added by white noise. The downstream area (red line) exhibits larger oscillation amplitudes and tracks the external input more closely than the upstream area (yellow line). (b) Normalized power spectral density of the external input (blue line), upstream area (yellow line) and downstream area (red line). The downstream area reconstructs the high-frequency components filtered by the upstream area. (c) Relationship between the phase shift (temporal lag) of the downstream node with the external input and the timescale localization metric $M_{TL}$, demonstrating that decreasing $M_{TL}$ reduces the temporal lag, a trend that holds across a wide range of input frequencies.

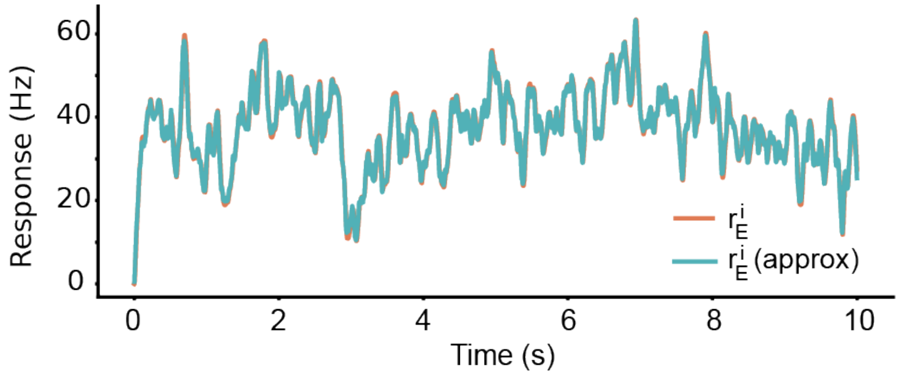


**Fig. S3 The approximation of the *minimal E to E-I* model.** Dynamics of the full model (equations (3)-(5)) versus the approximation model (equation (13)), in response to an external input modeled as filtered white noise (an Ornstein–Uhlenbeck process with timescale of 20 ms). The firing rates of the downstream area, depicted by the red (full model) and blue (approximation model) lines, demonstrate a near-perfect overlap.

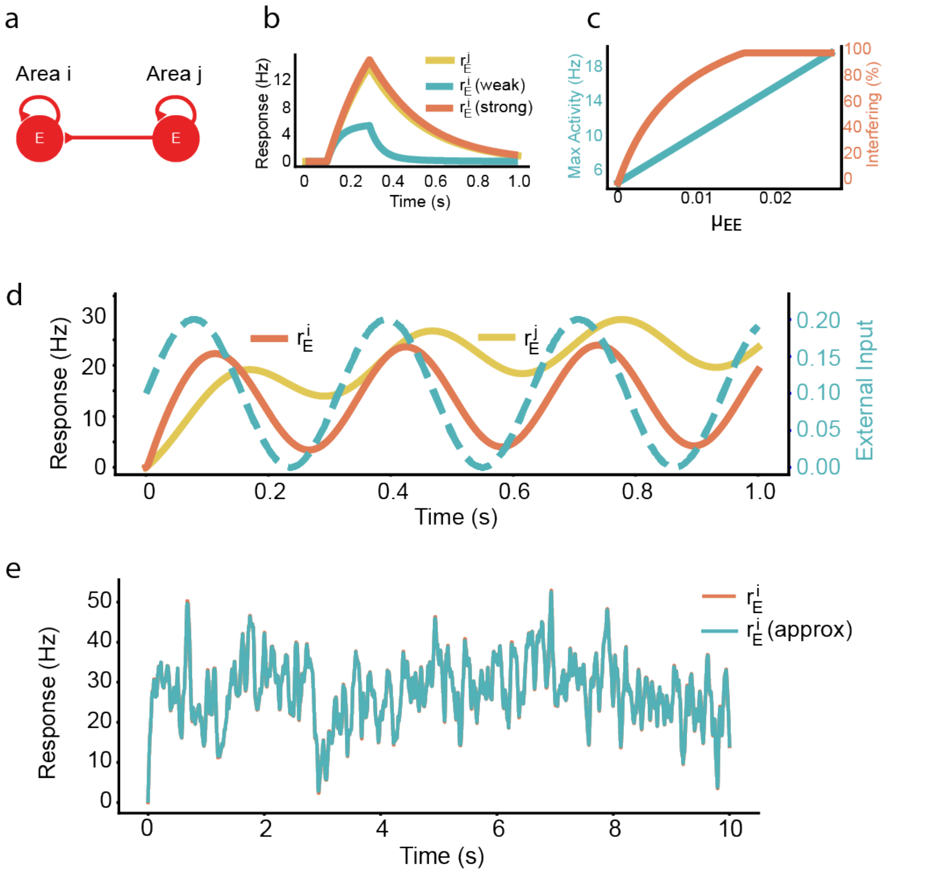


**Fig. S4 Dynamics of the *simplified E to E model* and *simplified E to E-I model* with varied input and connection strengths.** (a) Scheme of the *simplified E to E model*, with the directed interaction from area $j$ to area $i$ shown. (b) Response of the *simplified E to E model* to an external pulse input. Yellow: firing rate of upstream population $r_E^j$. Blue and red: firing rates of downstream population $r_E^i$ under weak and strong connection strengths. Dash blue: scaled downstream response under weak connection. A small independent external input is applied to the downstream area. (c) Signal propagation and timescale interference in the *simplified E to E model* under different connection strengths. Blue line, peak firing rate of the downstream population. Red line, interference measure defined by the proportion of the component from the upstream population to the total (see text). (d) Response of the *E-to-EI* model (with local recurrent connections) to an oscillatory external input (shown in dashed blue). Yellow and red: firing rate of upstream population $r_E^j$ and downstream population $r_E^i$. (e) Dynamics of the full model (equations (38)-(40)) versus the approximation model (equation (76)), in response to an external input modeled as filtered white noise (an Ornstein–Uhlenbeck process with timescale of 20 ms). The firing rates of the downstream area, depicted by the red (full model) and blue (approximation model) lines, demonstrate a near-perfect overlap.

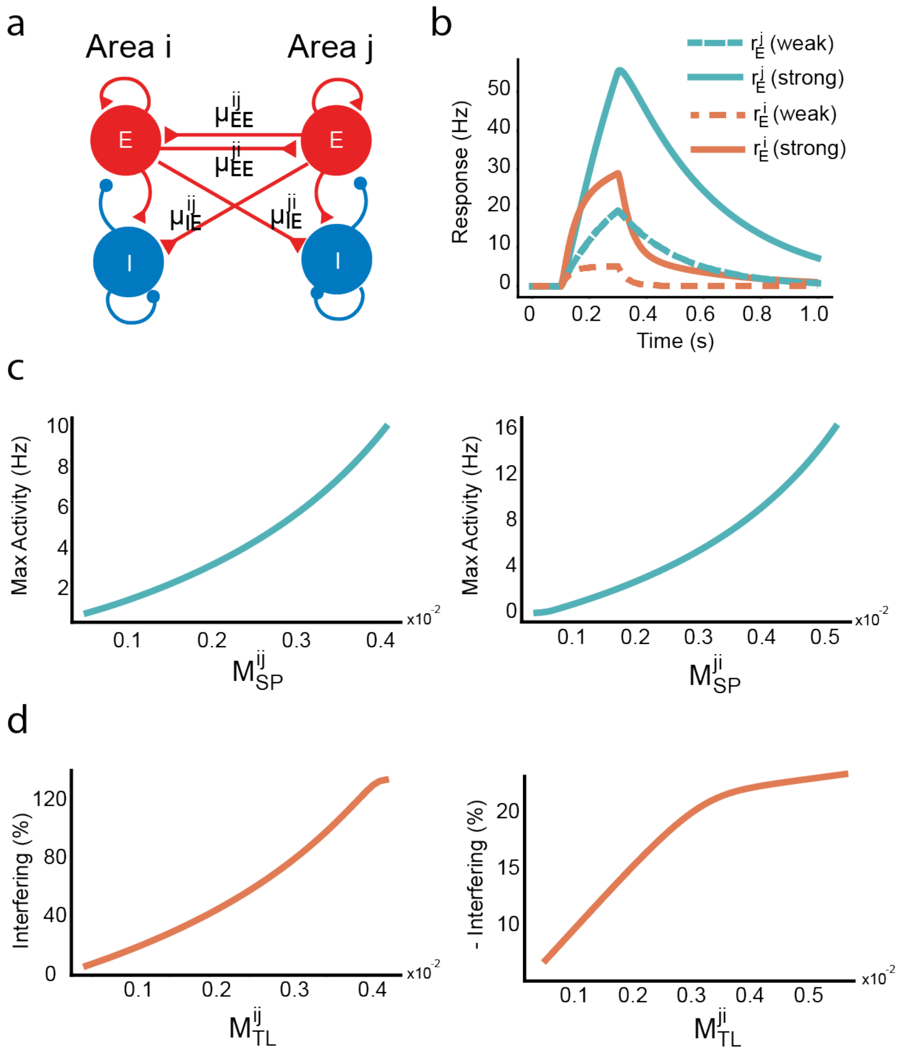


**Fig. S5 Dynamics of the *simplified two-area model with reciprocal connections.*** (a) Scheme of the simplified two-area model with reciprocal connections, showing interactions between area $i$ and area $j$. (b) Response of the model to an external pulse input applied to both areas. Blue and red curves represent the firing rates of the $j$ and $i$ populations, respectively, under weak (dashed lines) and strong (solid lines) connection strengths. (c) Correlation between the max firing rate of areas $i$ ($j$) and their corresponding signal propagation metrics $M_{SP}^{ij}$ ($M_{SP}^{ji}$), illustrating its effectiveness as a metric for signal propagation. (d) Correlation between the interference metric for area $i$ ($j$) and the timescale localization metrics $M_{TL}^{ij}$ ($M_{TL}^{ji}$), illustrating its effectiveness as a metric for timescale localization.

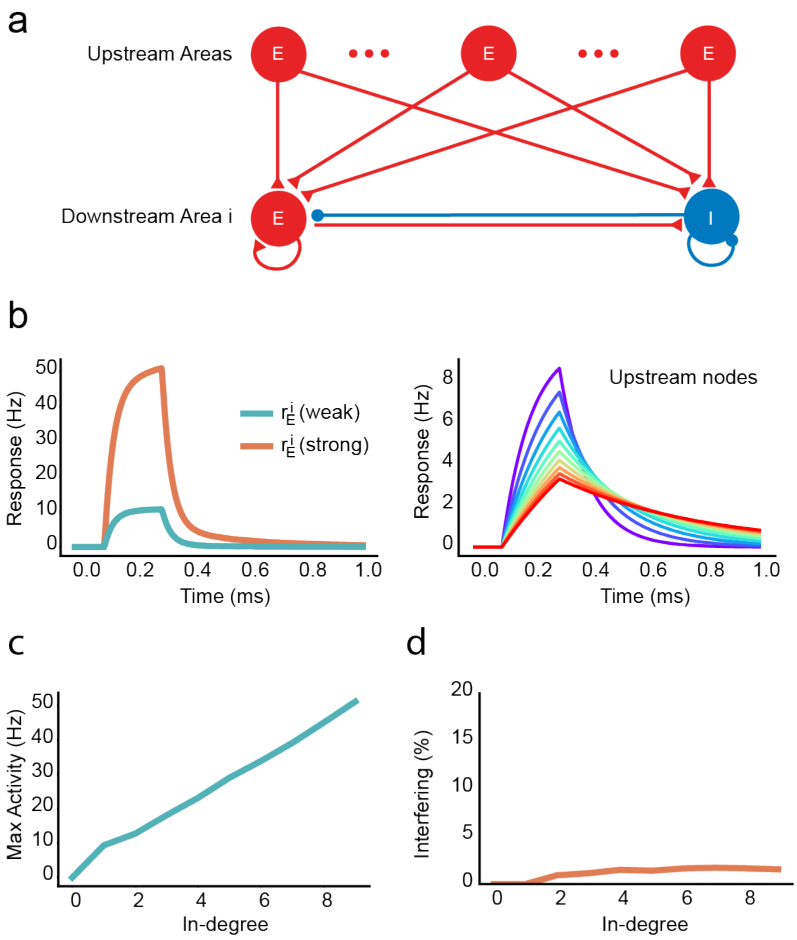


**Fig. S6 Dynamics of the *simplified model with multiple upstream areas.*** (a) Scheme of the simplified model with multiple upstream areas, showing interactions from upstream areas to the downstream area $i$. (b) Response of the model to an external pulse input applied to all upstream areas. Left: Firing rates of the downstream area iii under weak (blue) and strong (red) connection strengths. Right: Firing rates of the upstream areas, with colors representing different areas with distinct timescales. (c) Maximum firing rate of the downstream area $i$ as a function of its in-degree, showing an increase with higher in-degree. (d) The interference metric remains small and is largely independent of the in-degree of the area in the IFP regime.

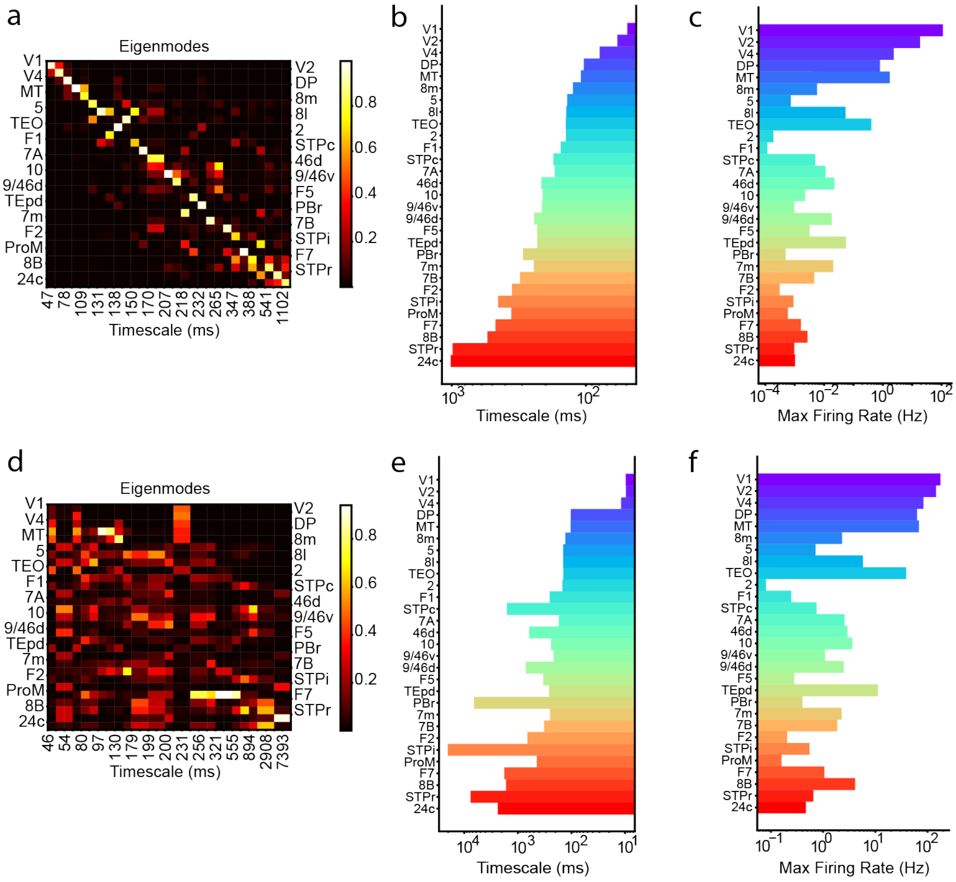


**Fig. S7 Signal dynamics in the multi-regional model of macaque cortex, with parameters falling in different regions.** (a) - (c) Simulation results with parameters in the region of [1]. (a) Visualization of the weight matrix eigenvectors, with associated timescales provided below. (b) The timescale derived from the fitting of auto-correlation function during the resting state. (c) Peak firing rate across different brain areas following a pulse stimulus to V1. (d) - (f) Simulation results with parameters in the region of [2].

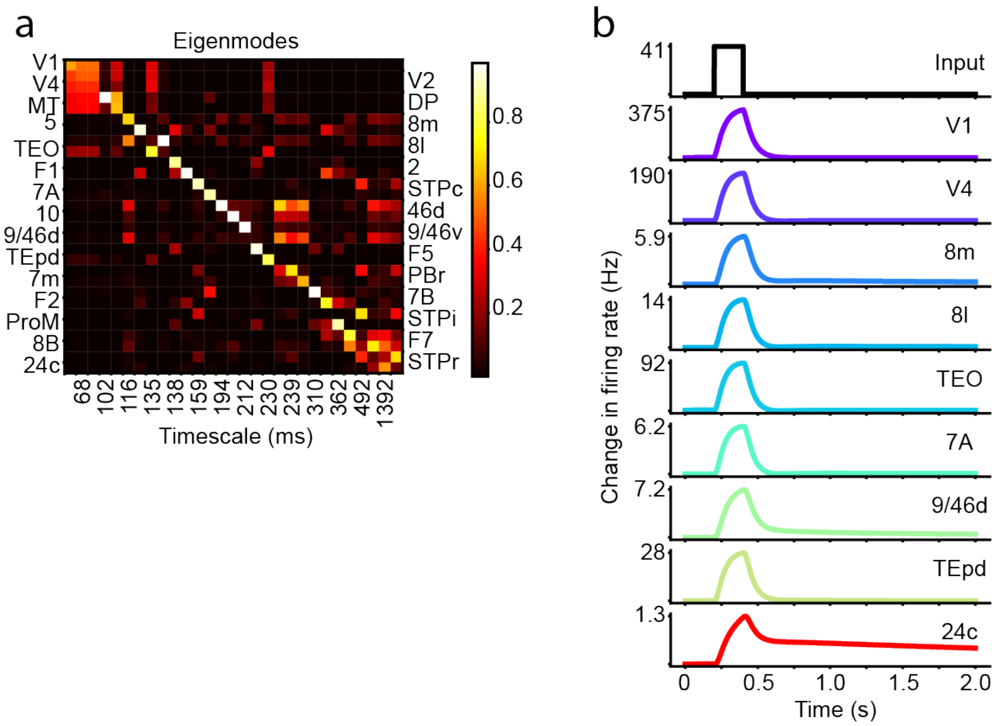


**Fig. S8 Additional results of IFP regime in the multi-areal rate model of macaque cortex.** (a) Visualization of the weight matrix eigenvectors, with associated timescales provided below. (b) The simulated activity of selected areas following a pulse stimulus to V1.

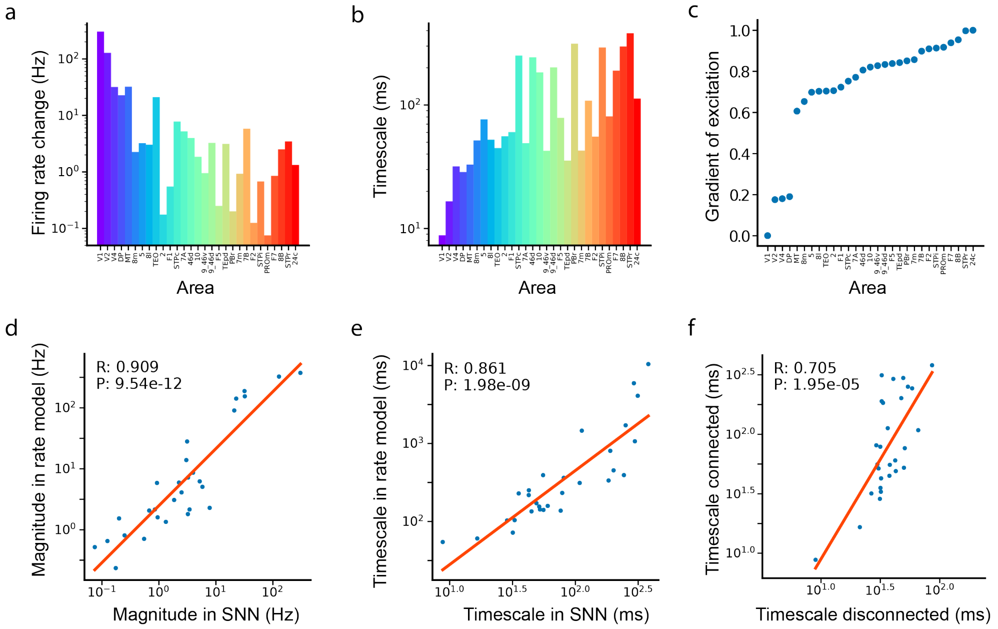


**Fig. S9 The dynamical property of the multi-area spiking network.** (a) Firing rate changes in each area when area V1 receives Poisson inputs. (b) Timescale distribution across areas derived from the auto-correlation functions when each area receives background Poisson inputs. (c) hierarchy value for each area in the SNN. (d) Comparison of mean firing rate changes between the spiking model and the rate model. (e) Comparison of timescales between the spiking model and the rate model. (f) Comparison of timescales between the spiking model and the Poission network model by replacing each upstream neuron activity with a Poisson neuron group exhibiting the same firing rate.

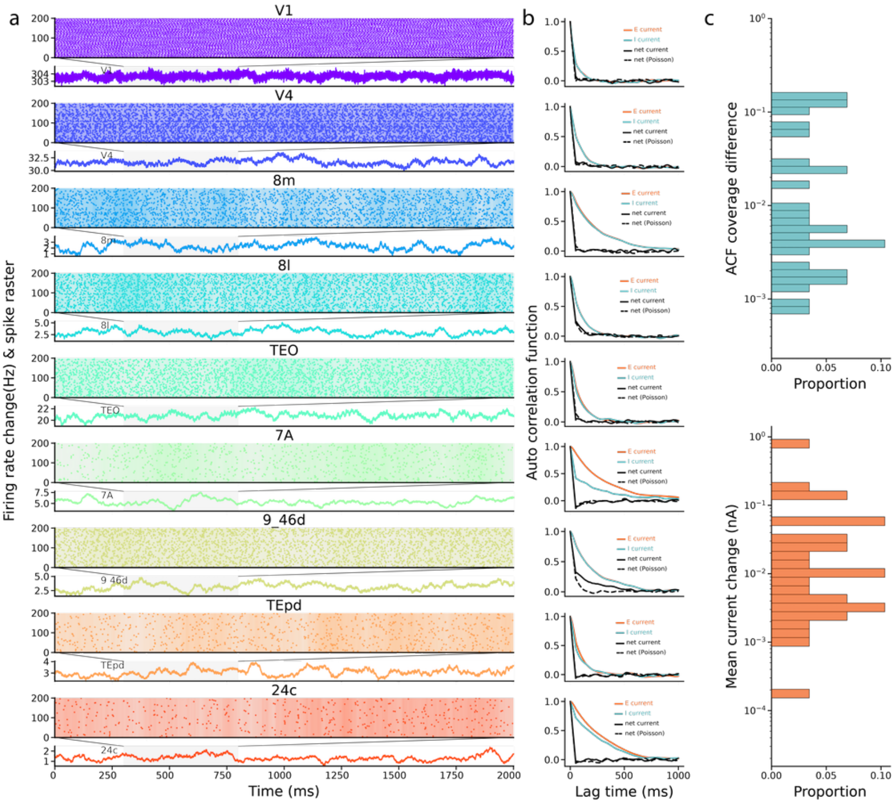


**Fig. S10 Evidence of IFP in the multi-area spiking network.** (a) The spike raster plot of a subgroup excitatory neurons and the population firing rate change, compared with pre-stimulus period, in a time interval during stimulus period of selective brain areas. The areas are arranged along the cortical hierarchy, and different areas exhibit different timescales. (b) Autocorrelation functions for the inter-areal excitatory current from all upstream E populations (red), intra-areal inhibitory current from the I population (blue), and the net current (solid black). The dashed black line represents the autocorrelation function of the reference net current when the inputs from other areas are replaced by Poisson inputs with the same frequency. The overlap between the solid and dashed curves indicates temporal cancellation of the E and I currents. (c) The histogram of the difference between the coverage of auto-correlation function of net input current in the spiking network and the Poisson network (top), and that of the magnitude of the mean total synaptic input (down) across the whole brain.

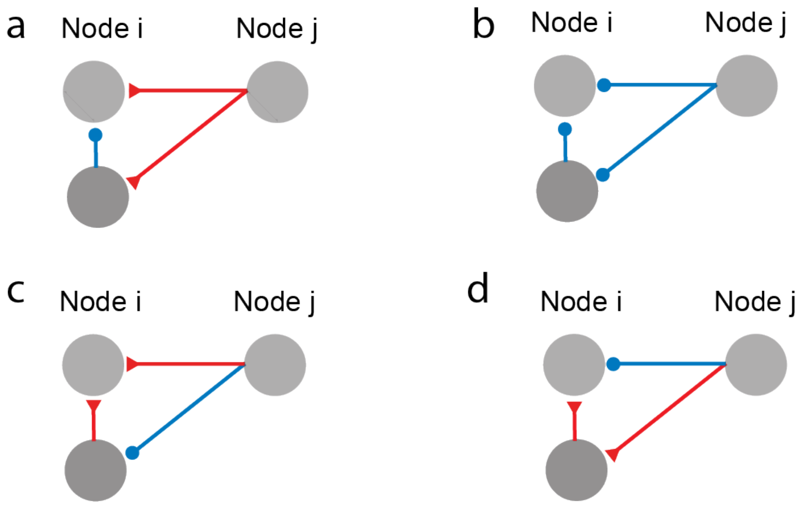


**Fig. S11 Representative 3-node motifs for IFP.** (a) The 3-node motif discussed in detail in this paper. (b–d) Other 3-node motifs capable of implementing IFP. Red and blue arrows represent excitatory and inhibitory projections, respectively. Shared characteristics include: (1) Two components for the downstream node, and (2) one effectively excitatory and one effectively inhibitory pathway from the upstream node to the downstream node.

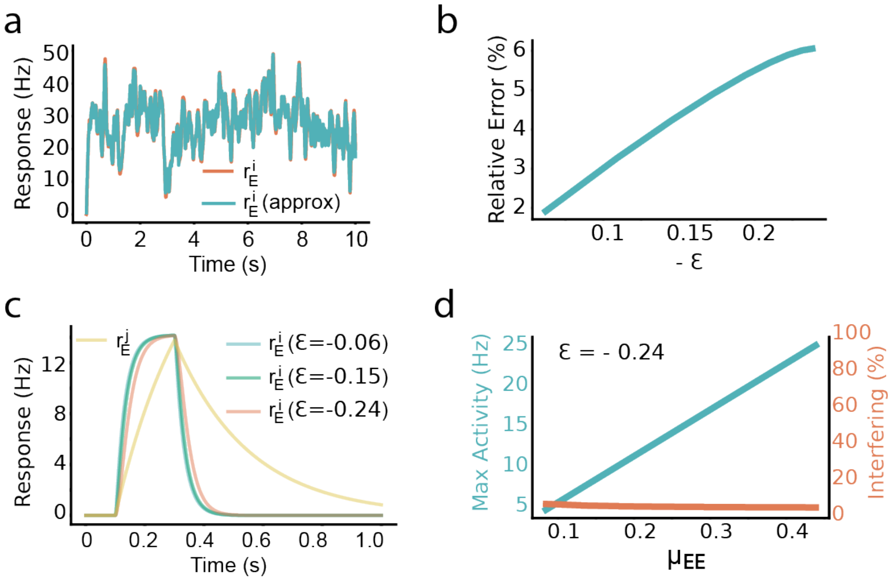


**Fig. S12 Two-areal model approximation and IFP for different epsilon values.** (a) Dynamics of the full model vs. the approximation, in response to an external input modeled as filtered white noise (Ornstein–Uhlenbeck process, 20 ms timescale). Here, asymptotic limit are not met ($\varepsilon$ = -0.24). The downstream area's firing rates (red: full model, blue: approximation) show a small deviation. (b) The relative mean square error of model approximation increases with $\varepsilon$, showing a small but growing error. (c) Model response to a pulse input for different $\varepsilon$ magnitudes. Yellow: upstream firing rate. Blue, green and red: downstream firing rates for small, medium, and large $\varepsilon$ magnitudes, respectively. (d) Signal propagation and timescale interference in the large $\varepsilon$ regime for different connection strengths. Blue: downstream peak firing rate. Red: interference measure, defined as the proportion of upstream influence on the total.

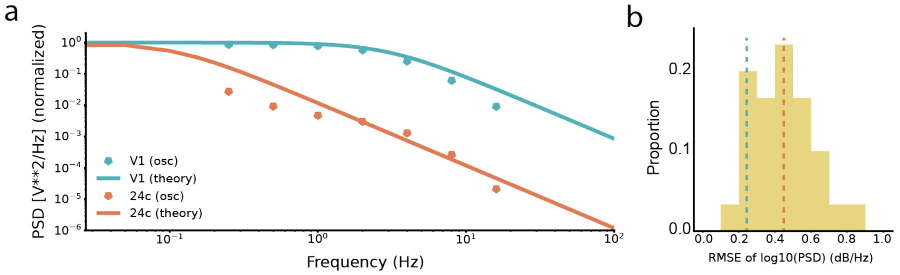


**Fig. S13 Response of the multi-area network model when area V1 receives oscillatory inputs with seven frequencies ranging from 0.25 Hz to 16 Hz.** (a) Dots: the frequency response of area V1 (blue) and 24c (red), Lines: the theoretical frequency response predicted by a low-pass filter with the characteristic timescale of the corresponding brain area measured by the autocorrelation function of population activity under white noise drive (resting state). (b) Histogram of the root mean squared error (RMSE) between the model's frequency response and the theoretical prediction across all areas. Vertical lines are the RMSE for area V1 (blue) and 24c (red), respectively.

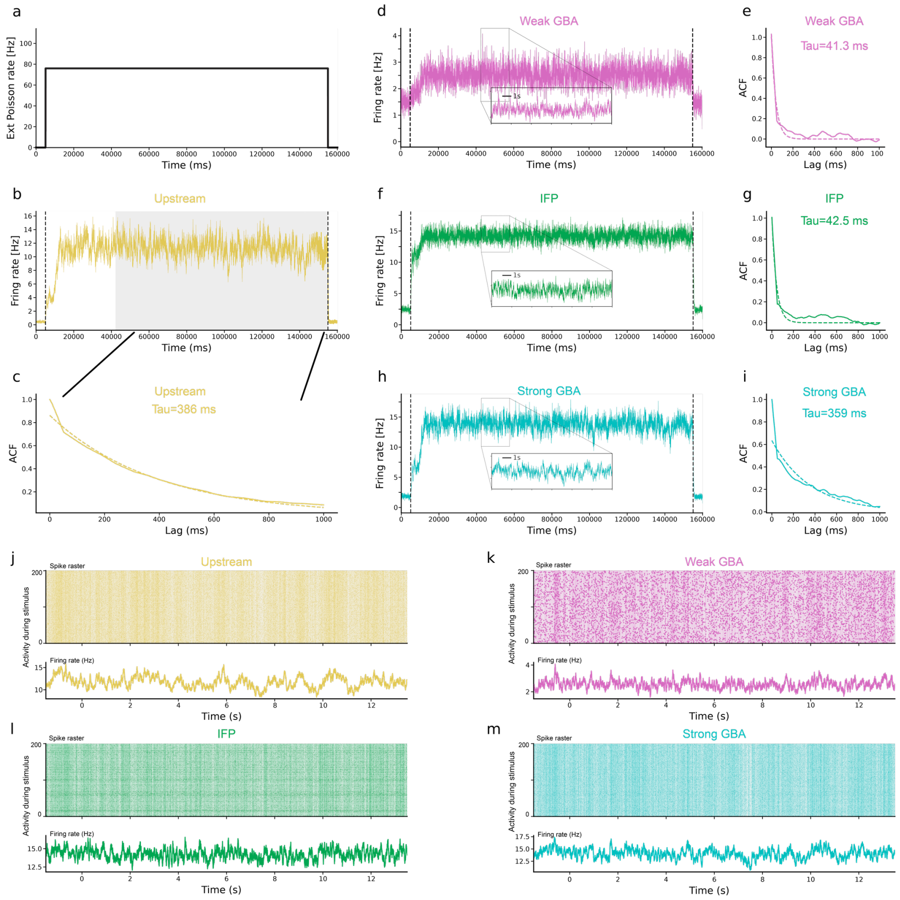


**Fig. S14 Population firing activity of the two-area spiking neuron network.** (a) The rate of external Poisson input to the upstream E neurons. (b) The population firing rate of the upstream area. (c) The autocorrelation function (solid curve) for stable upstream response in the shaded time window and its exponential fit (dash curve) to obtain its timescale about 386 ms. (d-e) In the weak GBA regime, the downstream E neurons show small changes in firing rate ($\sim$1Hz), and their timescale is unaffected by the upstream population (downstream timescale 41.3ms). (f-g) In the IFP regime, signals propagate reliably ($\sim$10Hz), and the timescale remains unaffected (downstream timescale 42.5ms). (h-i) In the strong GBA regime, signals also propagate reliably ($\sim$10Hz), but the timescale of the downstream group is close to that of the upstream group (downstream timescale 359ms). (j-m) Spike raster plots of a subgroup of excitatory neurons in the upstream area and downstream area in the weak GBA, IFP and strong GBA regime. The gradient-colored background behind the spike raster reflects the firing rate, with the hue corresponding to the rate at each time point.

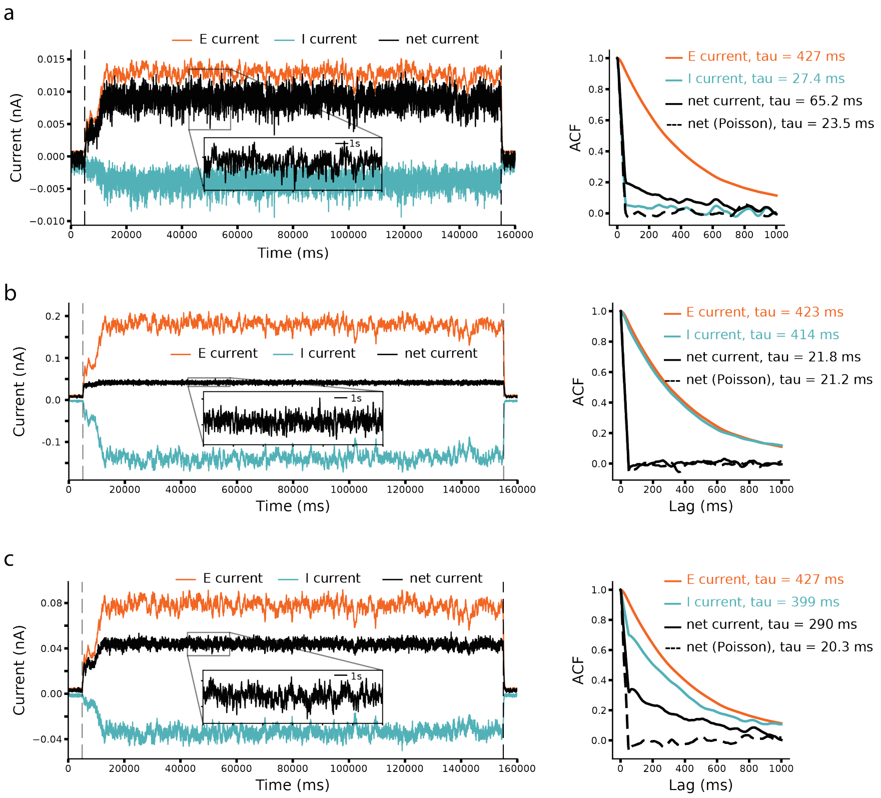


**Fig. S15 Currents of the downstream neuron population in the two-areal SNN.** (a-c) left, the dynamics of inter-areal excitatory current directly from the upstream E population (red), the intra-areal inhibitory current from the downstream I population (blue), and the net current as the sum of the E and I currents (black). Right, Solid curves: the auto-correlation functions of the three currents indicating their timescales. Dash curve: the reference auto-correlation function of the net current when the downstream area receives Poisson inputs with no information of upstream timescale. (a) The weak GBA case. The mean of the net current is small. (b) The IFP case. The mean of the net input is large, meanwhile the timescale of the net input is fast as a result of the complete temporal component cancellation of the E and I currents. (c) The strong GBA case. The mean of the net input is large, but the timescale of the net input is slow as a result of the incomplete temporal component cancellation of the E and I currents.

### 2 Text S1: The two-area models

#### 2.1 Description of minimal two-area models

We begin by presenting two minimal models that capture the dynamics between two connected brain regions, labeled as nodes $i$ and $j$. Each node's dynamics are described by a set of linear ordinary differential equations that model the neural activity within these regions. The internal connections within each node are simplified, allowing us to focus on the interactions between the two nodes. While this setup is simplified compared to realistic brain networks, it effectively conveys the key ideas of this paper and sheds light on how the described motif can be generalized to broader applications.

##### 2.1.1 the E to E model

Our initial model considers a motif in which an excitatory neuronal group in one region sends input to another excitatory group in a different region. This model, referred to as the *E to E model*, serves as a baseline for exploring inter-area neural interactions. The scheme is illustrated in Figure 1 (in the main text) and the model is mathematically formulated as follows:

$$\frac{dr_E^i}{dt} = -\frac{1}{\tau_E^i} r_E^i + \mu_{EE} r_E^j + I^i(t), \tag{1}$$

$$\frac{dr_E^j}{dt} = -\frac{1}{\tau_E^j} r_E^j + I^j(t). \tag{2}$$

In this formulation, $r_E^i$ and $r_E^j$ denote the firing rates of excitatory neuron populations in regions $i$ and $j$, respectively. The parameter $\tau_E^i, \tau_E^j$ represents the effective time constant of firing rate of neuron population $i$ and $j$, respectively and $\mu_{EE}$ defines the coupling strength of the inter-areal excitatory input from region $j$ to region $i$. External inputs to regions $i$ and $j$ are represented as $I^i(t)$ and $I^j(t)$, respectively.

Following the parameter settings from a previous study [3], we set $\tau_E^i = 50\ ms, \tau_E^j = 250\ ms$. The coupling strength $\mu_{EE}$ is varied depending on the scenarios. For Figure 1B, we choose $\mu_{EE} = 0.0005/ms$ for weak connection and $\mu_{EE} = 0.02/ms$ for strong connection. For Figure 1C, $\mu_{EE}$ is varied between 0 to $0.02/ms$.

##### 2.1.2 the E to E and I model

To extend the basic E-to-E model, we introduce an inhibitory neuron group alongside the excitatory group in region $i$, while region $j$ remains exclusively excitatory. The inter-areal connection from region $j$ to region $i$ involves excitatory projections from $j$ to both the excitatory and inhibitory populations in $i$. The scheme is shown in Figure 1D.

The dynamics of this extended model are described by the following equations:

$$\frac{dr_E^i}{dt} = -r_E^i/\tau_E^i - w_{EI} r_I^i + \mu_{EE} r_E^j + I^i(t), \tag{3}$$

$$\frac{dr_I^i}{dt} = -r_I^i/\tau_I^i + \mu_{IE} r_E^j, \tag{4}$$

$$\frac{dr_E^j}{dt} = -r_E^j/\tau_E^j + I^j(t). \tag{5}$$

where $r_E^i, r_I^i$ are the firing rates of excitatory and inhibitory groups in area $i$, respectively and $r_E^j$ is the firing rate of excitatory group in area $j$. The parameter $\tau_E^i, \tau_I^i, \tau_E^j$ represent the effective time constant for firing rate of excitatory and inhibitory population in the node $i$ and $j$, respectively. $w_{EI}$ denotes the coupling strength from the inhibitory population to the excitatory population within the region $i$ and $\mu_{XE}$ represents the coupling strength of the inter-areal input from the excitatory population to the $X$ population in a downstream cortical area. Other parameters retain their definitions from the "E to E model".

Based on [3], we set $\tau_E^i = 50\ ms, \tau_I^i = 10\ ms, \tau_E^i = 250\ ms, w_{EI} = 0.083/ms$ , consistent with empirical findings. For Figure 1E, we set $\mu_{EE} = 0.031/ms, \mu_{IE} = 0.035/ms$ for weak connection and $\mu_{EE} = 0.310/ms, \mu_{IE} = 0.355/ms$ for strong connection (also for Figure 1G). For Figure 1F, the $\mu_{EE}$ is varied between $0/ms$ and $0.763/ms$ and $\mu_{IE}$ changes from $0/ms$ to $0.882/ms$, correspondingly.

#### 2.2 Analysis of minimal two-area models

We now turn to the analysis of the minimal two-area models, with a particular focus on the *E to E-I model*. We start by the asymptotic analysis to derive the two metrics for signal propagation and timescale localization. By solving the linear equations analytically, we demonstrate how the activity in the upstream node influences the activity in the downstream node. Specifically, we show how the mean and temporal fluctuations in the downstream node are controlled by the upstream node activity, and we discuss the two system-defined metrics that govern these quantities. These results reveal a parameter regime where the mean and fluctuations of the downstream node are modulated by the upstream node in distinct ways, which we term as as *interference-free propagation*.

##### 2.2.1 Asymptotic analysis of *E to E-I model*

We start from transforming the dynamics of the *E to E-I model* (3-4) into a second-order ordinary differential equation (ODE) system, focusing solely on the firing rates of the excitatory groups. This simplification eliminates the variables representing the firing rates of the inhibitory group.

First, given equation (4), we can express $w_{EI} r_I^i$ in terms of other variables as follow:

$$w_{EI} r_I^i = -\frac{dr_E^i}{dt} - r_E^i/\tau_E^i + \mu_{EE} r_E^j + I^i, \tag{6}$$

This allows us to reformulate the system (differentiating equation (3) one more time):

$$\frac{d^2 r_E^i}{dt^2} = -\frac{1}{\tau_E^i}\frac{dr_E^i}{dt} - w_{EI}\frac{dr_I^i}{dt} + \mu_{EE}\frac{dr_E^j}{dt} + \frac{dI^i}{dt}$$

$$= -\frac{1}{\tau_E^i}\frac{dr_E^i}{dt} - w_{EI}\left(-\frac{r_I^i}{\tau_I^i} + \mu_{IE}r_E^j\right) + \mu_{EE}\frac{dr_E^j}{dt} + \frac{dI^i}{dt}$$

$$= -\frac{1}{\tau_E^i}\frac{dr_E^i}{dt} - w_{EI}\mu_{IE}r_E^j + \mu_{EE}\frac{dr_E^j}{dt} + \frac{dI^i}{dt} + \frac{1}{\tau_I^i}w_{EI}r_I^i$$

$$= -\frac{1}{\tau_E^i}\frac{dr_E^i}{dt} - w_{EI}\mu_{IE}r_E^j + \mu_{EE}\frac{dr_E^j}{dt} + \frac{dI^i}{dt} + \frac{1}{\tau_I^i}\left(-\frac{dr_E^i}{dt} - r_E^i/\tau_E^i + \mu_{EE}r_E^j + I^i\right)$$

$$= -\left(\frac{1}{\tau_E^i} + \frac{1}{\tau_I^i}\right)\frac{dr_E^i}{dt} - \frac{1}{\tau_E^i\tau_I^i}r_E^i + \left(\mu_{EE}/\tau_I^i - w_{EI}\mu_{IE}\right)r_E^j + \mu_{EE}\frac{dr_E^j}{dt} + \frac{I^i}{\tau_I^i} + \frac{dI^i}{dt} \quad (7)$$

The reformulation results in a second-order ODE for $r_E^i$, capturing the dynamics without explicitly including $r_I^i$:

$$\frac{d^2r_E^i}{dt^2} + \left(\frac{1}{\tau_E^i} + \frac{1}{\tau_I^i}\right)\frac{dr_E^i}{dt} + \frac{1}{\tau_E^i\tau_I^i}r_E^i$$
$$= \left(\mu_{EE}/\tau_I^i - w_{EI}\mu_{IE}\right)r_E^j + \mu_{EE}\frac{dr_E^j}{dt} + \frac{I^i}{\tau_I^i} + \frac{dI^i}{dt} \quad (8)$$

Further defining non-dimensional variables: $R_E^k = (1/\tau_E^i + 1/\tau_I^i)^{-1}r_E^k$ $(k = i, j)$, $\tau = t/(\tau_E^i + \tau_I^i)$, we further transform the second order system (equation (8)) into a dimensionless system as follow (assuming $I^i = 0$):

$$\frac{1}{\tau_E^i\tau_I^i\left(\tau_E^i + \tau_I^i\right)}\frac{d^2R_E^i}{d\tau^2} + \frac{\tau_E^i + \tau_I^i}{\left(\tau_E^i\tau_I^i\right)^2}\frac{dR_E^i}{d\tau} + \frac{\tau_E^i + \tau_I^i}{\left(\tau_E^i\tau_I^i\right)^2}R_E^i$$
$$= \frac{\tau_E^i + \tau_I^i}{\tau_E^i\tau_I^i}\left[\left(\mu_{EE}/\tau_I^i - w_{EI}\mu_{IE}\right)R_E^j + \frac{\mu_{EE}}{\tau_E^i + \tau_I^i}\frac{dR_E^j}{d\tau}\right] \quad (9)$$

which can be simplified into

$$\epsilon\frac{d^2R_E^i}{d\tau^2} - \frac{dR_E^i}{d\tau} - R_E^i = -\tau_E^i\tau_I^i\left[M_{SP}R_E^j + \frac{\mu_{EE}}{\tau_E^i + \tau_I^i}\frac{dR_E^j}{d\tau}\right], \quad (10)$$

where $\epsilon = -1/(\sqrt{\tau_E^i/\tau_I^i} + \sqrt{\tau_I^i/\tau_E^i})^2$, and $M_{SP} = \mu_{EE}/\tau_I^i - w_{EI}\mu_{IE}$.

By viewing the parameter $\epsilon$ in equation (10) as small, we reduce model (10) to an approximated *E to E model* as:

$$\frac{dR_E^i}{d\tau} = -R_E^i + \tau_E^i\tau_I^i\left[M_{SP}R_E^j + \frac{\mu_{EE}}{\tau_E^i + \tau_I^i}\frac{dR_E^j}{d\tau}\right] \quad (11)$$

Transforming $R_E^i, R_E^j, \tau$ back to the original variables, we have the following

$$\frac{dr_E^i}{dt} = -\frac{1}{\tilde{\tau}_E^i} r_E^i + \frac{M_{SP}}{T} r_E^j + \frac{\mu_{EE}}{T} \frac{dr_E^j}{dt}, \tag{12}$$

where the effective time constant $\tilde{\tau}_E^i = \tau_E^i + \tau_I^i$, $T = 1/\tau_E^i + 1/\tau_I^i$. This reduction reveals an additional activity driver of the downstream population in equation (12), represented by the derivative of the upstream population's activity, $\frac{\mu_{EE}}{T}\frac{dr_E^j}{dt}$. In addition, by combining equations (5) and (12), we have

$$\frac{dr_E^i}{dt} = -\frac{1}{\tilde{\tau}_E^i} r_E^i + \frac{M_{TL}}{T} r_E^j + \frac{\mu_{EE}}{T} I^j(t), \tag{13}$$

where $M_{TL} = M_{SP} - \mu_{EE}/\tau_E^j$ (the accuracy of the approximation is numerically validated in Figure S3).

##### 2.2.2 Explicit solutions of *E to E-I model*

We now turn to an explicit solution of the system (equation (3) - (5)) to investigate whether similar derivations can be achieved. As demonstrated below, the same formulas and definitions for the two metrics $M_{SP}, M_{TL}$ , can be derived using this approach.

We begin by decomposing the neuronal activities into their mean components and the temporally varying components as follows:

$$r_E^i = \bar{r}_E^i + \Delta r_E^i(t), \quad r_I^i = \bar{r}_I^i + \Delta r_I^i(t), \quad r_E^j = \bar{r}_E^j + \Delta r_E^j(t), \quad I^x = \bar{I}^x + \Delta I^x(t), x = i, j. \tag{14}$$

Substituting the decomposition into Eq.(5), we obtain:

$$\frac{d\Delta r_E^j}{dt} = -\left(\bar{r}_E^j + \Delta r_E^j\right)/\tau_E^j + \bar{I}^j + \Delta I^j, \tag{15}$$

which leads to that

$$\bar{r}_E^j = \tau_E^j \bar{I}^j, \quad \frac{d\Delta r_E^j}{dt} = -\Delta r_E^j/\tau_E^j + \Delta I^j. \tag{16}$$

Assuming $\Delta r_E^j(0) = 0$, the explicit formula of $\Delta r_E^j$ is:

$$\Delta r_E^j(t) = \int_0^t e^{-(t-s)/\tau_E^j} \Delta I^j(s) ds. \tag{17}$$

Next, substituting these into Eq.(4) gives:

$$\frac{d\Delta r_I^i}{dt} = -\left(\bar{r}_I^i + \Delta r_I^i\right)/\tau_I^i + \mu_{IE}\left(\bar{r}_E^j + \Delta r_E^j\right), \tag{18}$$

which leads to

$$\bar{r}_I^i = \tau_I^i \mu_{IE} \, \bar{r}_E^j = \tau_E^j \tau_I^i \mu_{IE} \bar{I}^j, \tag{19}$$

$$\frac{d\Delta r_I^i}{dt} = -\Delta r_I^i / \tau_I^i + \mu_{IE} \Delta r_E^j. \tag{20}$$

This leads to the following solution for $\Delta r_I^i$:

$$\begin{aligned}
\Delta r_I^i(t) &= \mu_{IE} \int_0^t e^{-(t-s)/\tau_I^i} \Delta r_E^j(s) ds & (21)\\
&= \frac{\mu_{IE}}{1/\tau_I^i - 1/\tau_E^j} \left( \int_0^t e^{-(t-s)/\tau_E^j} \Delta I^j(s) ds - \int_0^t e^{-(t-s)/\tau_I^i} \Delta I^j(s) ds \right) & (22)\\
&= \frac{\mu_{IE}}{1/\tau_I^i - 1/\tau_E^j} \left( \Delta r_E^j(t) - \int_0^t e^{-(t-s)/\tau_I^i} \Delta I^j(s) ds \right) & (23)
\end{aligned}$$

Finally, substituting these into Eq.(3) results in:

$$\frac{d\Delta r_E^i}{dt} = - \left( \bar{r}_E^i + \Delta r_E^i \right) / \tau_E^i - w_{EI} \left( \bar{r}_I^i + \Delta r_I^i \right) + \mu_{EE} \left( \bar{r}_E^j + \Delta r_E^j \right) + \bar{I}^i + \Delta I^i, \tag{24}$$

which leads to the solution of steady state for $\bar{r}_E^i$:

$$\begin{aligned}
\bar{r}_E^i &= \tau_E^i \left( -w_{IE} \bar{r}_I^i + \mu_{EE} \bar{r}_E^j + \bar{I}^i \right) = \tau_E^i \left( -w_{IE} \tau_I^i \mu_{IE} \, \bar{r}_E^j + \mu_{EE} \bar{r}_E^j + \bar{I}^i \right) \\
&= \tau_E^i \tau_I^i \left( \mu_{EE} / \tau_I^i - w_{IE} \mu_{IE} \right) \bar{r}_E^j + \tau_E^i \bar{I}^i, & (25)
\end{aligned}$$

For the temporal fluctuations, we obtain:

$$\begin{aligned}
\frac{d\Delta r_E^i}{dt} &= -\Delta r_E^i / \tau_E^i - w_{EI} \Delta r_I^i + \mu_{EE} \Delta r_E^j + \Delta I^i \\
&= -\Delta r_E^i / \tau_E^i - w_{EI} \frac{\mu_{IE}}{1/\tau_I^i - 1/\tau_E^j} \left( \Delta r_E^j(t) - \int_0^t e^{-(t-s)/\tau_I^i} \Delta I^j(s) ds \right) + \mu_{EE} \Delta r_E^j + \Delta I^i \\
&= -\Delta r_E^i / \tau_E^i + \left( \mu_{EE} - \frac{w_{EI} \mu_{IE}}{1/\tau_I^i - 1/\tau_E^j} \right) \Delta r_E^j \\
&+ \frac{w_{EI} \mu_{IE}}{1/\tau_I^i - 1/\tau_E^j} \int_0^t e^{-(t-s)/\tau_I^i} \Delta I^j(s) ds + \Delta I^i. & (26)
\end{aligned}$$

##### 2.2.3 Metrics for signal integration and propagation and IFP

Drawing from the explicit solution of the systems (Eq.(25), Eq.(26)), we propose two quantitative metrics that characterize timescale localization (the distinct timescales of the different nodes' activities) and signal propagation in this framework.

We define these two metrics as follows (same as the formula using asymptotic analysis):

$$M_{SP} = \mu_{EE}/\tau_I^i - w_{EI}\mu_{IE}, \tag{27}$$
$$M_{TL} = M_{SP} - \mu_{EE}/\tau_E^j. \tag{28}$$

From Eq.(25) and Eq.(26), we can deduce the following:

$$\bar{r}_E^i = \tau_E^i \tau_I^i \; M_{SP} \bar{r}_E^j + \tau_E^i \bar{I}^i. \tag{29}$$
$$\frac{d\Delta r_E^i}{dt} = -\frac{\Delta r_E^i}{\tau_E^i} + \frac{M_{TL}}{1/\tau_I^i - 1/\tau_E^j} \Delta r_E^j + \frac{w_{EI}\mu_{IE}}{1/\tau_I^i - 1/\tau_E^j} \int_0^t e^{-(t-s)/\tau_I^i} \Delta I^j(s) ds + \Delta I^i \tag{30}$$

From the analysis above, we can see that (29) and (30) demonstrate how the metrics $M_{SP}$ and $M_{TL}$ control the signal propagation and timescale localization. (29) illustrates how the magnitude of the response is influenced by the upstream signal $r_E^j$ and external input $I^i$, with the signal propagation being proportionally to the metric $M_{SP}$. Concurrently, (30) reveals how the fluctuations in the downstream node are influenced by the upstream node's activity, modulated by the metric $M_{TL}$. We have provided in the later section numerical results that support these findings. We also provide a more detailed mathematical analysis on how these metrics influence signal propagation and timescale localization in a more comprehensive toy model.

Moreover, given that the signal propagation and timescale localization are controlled by different metrics $M_{SP}$ and $M_{TL}$ separately, our analysis indicates a special parameter regime characterized by a large $M_{SP}$ and a small $M_{TL}$. In this regime, the magnitude of the downstream node's response to the upstream node is significant (large $M_{SP}$), but the temporal fluctuations introduced by the upstream node will be of no effect to the downstream node (small $M_{TL}$). This suggests that the two nodes can operate in their own characteristic timescales while allowing robust signal communication between them. We term this parameter regime as "interference-free propagation" (IFP).

##### 2.2.4 Physical Intuition of IFP mechanism

To provide an intuitive understanding of the IFP mechanism, we compute the input currents received by the downstream E population, from which the metrics $M_{SP}$ and $M_{TL}$ will naturally emerge. In the *E to E-I* model (Fig. 1D), the upstream area influences the downstream excitatory population activity through two pathways: a direct excitation pathway where the upstream activity excites the downstream excitatory population, and an indirect inhibition pathway where the upstream activity first excites the downstream inhibitory population, which then inhibits the downstream excitatory population. By decomposing the activity signals into their mean components and the temporally varying components, i.e., $r_X^k = \bar{r}_X^k + \Delta r_X^k(t), X = E, I, k = i, j$, we express the excitatory input to the downstream excitatory population from the direct pathway as

$$I_{E,syn} = \mu_{EE}\bar{r}_E^j + \mu_{EE}\Delta r_E^j(t).$$

In addition, through solving the downstream inhibitory population response $r_I^i(t)$, we can calculate the inhibitory input to the downstream excitatory population from the indirect pathway as

$$I_{I,syn} = -w_{EI}\mu_{IE}\tau_I^i\bar{r}_E^j - w_{EI}\mu_{IE}e^{-t/\tau_I^i} * \Delta r_E^j(t),$$

where $*$ denotes temporal convolution.

By setting $\Delta r_E^j = e^{-t/\tau_E^j} * \Delta I^j(t)$, i.e., the upstream response to a fluctuating external input $I^j = \bar{I}^j + \Delta I^j(t)$, we can compute that

$$\begin{aligned} e^{-t/\tau_I^i} * \Delta r_E^j(t) &= e^{-t/\tau_I^i} * \left(e^{-t/\tau_E^j} * \Delta I^j(t)\right) = \left(e^{-t/\tau_I^i} * e^{-t/\tau_E^j}\right) * \Delta I^j(t) \\ &= \frac{1}{1/\tau_I^i - 1/\tau_E^j}\left(e^{-t/\tau_E^j} - e^{-t/\tau_I^i}\right) * \Delta I^j(t) \\ &= \frac{1}{1/\tau_I^i - 1/\tau_E^j}\left(\Delta r_E^j(t) - e^{-t/\tau_I^i} * \Delta I^j(t)\right). \end{aligned} \tag{31}$$

It indicates that the total synaptic inputs $I_{total,syn} = I_{E,syn} + I_{I,syn}$ takes the form as

$$I_{total,syn} = M_{SP}\tau_I^i\bar{r}_E^j + \frac{M_{TL}}{1/\tau_I^i - 1/\tau_E^j}\Delta r_E^j(t) + R(t), \tag{32}$$

where the remainder term $R(t) = w_{EI}\mu_{IE}\tau_I^i\tau_E^j/(\tau_E^j - \tau_I^i)e^{-t/\tau_I^i} * \Delta I^j(t)$ operates only with inhibitory population's timescale $\tau_I^i$, independent of upstream timescale.

As seen in equation (32), $M_{SP}$ and $M_{TL}$ appear in the mean and temporal fluctuating components of the total synaptic inputs, respectively. A large value of $M_{SP}$ reflects a large mean current to the downstream excitatory population, which results from the imbalance between the mean of the excitatory inputs from the direct pathway and that of the inhibitory inputs from the indirect pathway. Therefore, $M_{SP}$ quantifies the effectiveness of signal propagation through the mean signals. In addition, a small value of $M_{TL}$ reflects a small fluctuating current component governed by the upstream activity timescale, which results from a substantial cancellation between the temporal fluctuations of the excitatory and inhibitory inputs from the two pathways. Therefore, $M_{TL}$ quantifies the degree of timescale localization.

##### 2.2.5 Other motifs for IFP

We also emphasize that while the minimal two-area model presented here is based on a neuronal network, the fundamental concepts underlying IFP can be realized through various other motifs and can probably generalize to other complex networks, as illustrated in Figure S11.

The key conditions for achieving IFP, as demonstrated in the figure, include the following:

1. Multiple Groups in One Node: A critical requirement is the presence of more than one neuronal group within a single local node. This introduces additional degrees of

freedom, enabling the separation of signal propagation and timescale integration. In contrast, in the simple E-to-E model, both signal propagation and timescale localization are controlled by a single parameter, $\mu_{EE}$.

2. Inhibitory Pathway: Another essential condition is that at least one pathway from the upstream to the downstream node must be effectively inhibitory. This inhibition allows for the suppression or cancellation of temporal fluctuations originating from the upstream node. In an entirely excitatory network, such cancellation is not possible, and the temporal variations would propagate to the downstream nodes without control.

#### 2.3 Comparison of IFP with balanced amplification

Murphy and Miller (2009) [4] introduced a mechanism known as "balanced amplification," as one way to amplify specific patterns of neural activities in cortical circuits, which was later expanded upon by Joglekar et al. (2018) [2] as "global balanced amplification." Notably, the "balanced amplification" initially described exhibits behaviors akin to the global system our work explores—particularly, a significant amplification of signal response with a relatively unchanged timescale, as exemplified in Figure 2, Murphy and Miller (2009). Our analysis reveals that this similarity is not coincidental but stems from underlying principles common to both our model and the balanced amplification mechanism.

The mechanism of balanced amplification can be exemplified through the following set of equations, representing the dynamics of excitatory ($r_E$) and inhibitory ($r_I$) populations:

$$\frac{dr_E}{dt} = -r_E + wr_E - k_I w r_I + I_E, \tag{33}$$

$$\frac{dr_I}{dt} = -r_I + wr_E - k_I w r_I, \tag{34}$$

where $w$ denotes the connection strength and $k_I$ the "balance factor". The "balanced amplification" is achieved when $k_I$ is slightly larger than 1.

Now we take the similar method to transform two-dimensional system into a second-order differential equation solely in terms of $r_E$, and it yields that:

$$(\ddot{r}_E - 2\dot{r}_E + r_E) + (k_I - 1)\, w\, (\dot{r}_E + r_E) = (1 + k_I w)\, I_E. \tag{35}$$

This formulation highlights that the essence of balanced amplification lies in $k_I$ being close to 1, rendering the term $(k_I - 1)\, w\, (\dot{r}_E + r_E)$ negligible, even for large $w$. Consequently, increasing $w$ has minimal impact on the equation's left-hand side, which governs the timescale of $r_E$ dynamics. In contrast, the right-hand side, representing the influence of external input, scales linearly with $w$. This decoupling of dynamic timescale and response amplitude enables the system to amplify responses without disrupting temporal dynamics, corroborating observations reported in the literature.

Our model aligns with the core premise of balanced amplification, positing that a balance between excitatory and inhibitory neuronal populations facilitates a separation between the dynamics' timescale and magnitude. This balance allows for efficient signal integration and propagation. However, our framework extends beyond local connectivity balance, incorporating global connections to enable biologically plausible information routing across different brain regions.

### 3 Text S2: The two-area models with local recurrent connections

#### 3.1 Description of two-area models with local recurrent connections

We start with two simplified, foundational models that encapsulate the dynamics between two interconnected brain regions, labeled as nodes $i$ and $j$. Each node is characterized by a set of linear ordinary differential equations, representing the neural activity within these areas. Compared with the minimal two-area models, the simplified two-area models introduce the connections within the local brain region, which can be applied directly to the neuron cortex modeling.

##### 3.1.1 the E to E model

Our initial focus is on a motif where an excitatory neuronal group in one area sends input to another excitatory group in a different area, henceforth referred to as the *E to E model.* This construct serves as a baseline to investigate the dynamics of neural interactions. The model is mathematically formulated as follows:

$$\frac{dr_E^i}{dt} = -\frac{1}{\tau_E^m} r_E^i + w_{EE} r_E^i + \mu_{EE} r_E^j + I^i(t), \tag{36}$$

$$\frac{dr_E^j}{dt} = -\frac{1}{\tau_E^j} r_E^j + I^j(t). \tag{37}$$

In this formulation, $r_E^i$ and $r_E^j$ denote the firing rates of excitatory neuron populations in areas $i$ and $j$, respectively. The parameter $\tau_E^m$ represents the intrinsic membrane time constant of excitatory neurons, $\tau_E^j$ represents the effective time constant of firing rate of neuron population $j$, $w_{EE}$ signifies the strength of self-recurrent excitation within each region, and $\mu_{EE}$ quantifies the coupling strength of the inter-areal excitatory input from area $j$ to area $i$. External inputs to areas $i$ and $j$ are represented as $I^i(t)$ and $I^j(t)$, respectively.

Adapted from a previous study [3], we set $\tau_E^m = 20\ ms, w_{EE} = 0.03/ms, \tau_E^j = 250ms$ while $\mu_{EE}$ is varied for different scenarios. For Figure 1B, we choose $\mu_{EE} = 0.00055/ms$ for weak connection and $\mu_{EE} = 0.01774/ms$ for strong connection. For Figure 1C, $\mu_{EE}$ is varied between 0 to $0.025/ms$.

##### 3.1.2 the E to E and I model

Expanding on the above basic model, we introduce an inhibitory neuron group alongside the excitatory group in area $i$, while area $j$ remains exclusively excitatory. The directed connection from area $j$ to area $i$ contains excitatory projections from area $j$ to both excitatory and inhibitory neuron groups in area $i$.

Therefore, the dynamics of the system can be described as follows:

$$\frac{dr_E^i}{dt} = -\frac{1}{\tau_E^m} r_E^i + w_{EE} r_E^i - w_{EI} r_I^i + \mu_{EE} r_E^j + I^i(t), \tag{38}$$

$$\frac{dr_I^i}{dt} = -\frac{1}{\tau_I^m} r_I^i + w_{IE} r_E^i - w_{II} r_I^i + \mu_{IE} r_E^j, \tag{39}$$

$$\frac{dr_E^j}{dt} = -\frac{1}{\tau_E^j} r_E^j + I^j(t). \tag{40}$$

where $r_E^i, r_I^i$ are the firing rates of excitatory and inhibitory groups in node $i$, respectively and $r_E^j$ is the firing rate of excitatory group in node $j$, while $\tau_E^m$ and $\tau_I^m$ are the corresponding intrinsic time constants, respectively. $w_{XY}$ symbolizes the coupling strength from $Y$ population to $X$ population within the area ($X, Y$ could be E or I population) and $\mu_{XE}$ represents the coupling strength of the inter-areal input from the excitatory population to the $X$ population in a downstream cortical area. Other parameters retain their definitions from the "E to E model". Following a previous study [3], we set $\tau_E^m = 20\ ms, \tau_I^m = 10\ ms, w_{EE} = 0.087/ms, w_{EI} = 0.083/ms, w_{IE} = 0.428/ms, w_{II} = 0.439/ms$, consistent with empirical findings. For Figure 1E, we set $\mu_{EE} = 0.400/ms, \mu_{IE} = 2.571/ms$ for weak connection and $\mu_{EE} = 1.788/ms, \mu_{IE} = 11.478/ms$ for strong connection (also for Figure 1G). For Figure 1F, the $\mu_{EE}$ is varied between $0.672/ms$ and $6.624/ms$ and $\mu_{IE}$ changes from $4.320/ms$ to $42.589/ms$, correspondingly.

#### 3.2 Asymptotic analysis of simplified two-area model

We now turn to the asymptotic analysis of the *E to E-I model*, showing that under the conditions of balancing between local excitatory and inhibitory currents, the network can be well approximated by an *E to E* only model, providing the foundation for the following analysis for metrics determining signal integration and propagation.

##### 3.2.1 Transformation to a second-order ODE system

We start from transforming the dynamics of the *E to E-I model* into a second-order ordinary differential equation (ODE) system, focusing solely on the firing rates of the excitatory groups. This simplification eliminates the variables representing the firing rates of the inhibitory group. Consider the following system:

$$\frac{dr_E^i}{dt} = a r_E^i - b r_I^i + c r_E^j + I^i(t), \tag{41}$$

$$\frac{dr_I^i}{dt} = d r_E^i - e r_I^i + f r_E^j, \tag{42}$$

$$\frac{dr_E^j}{dt} = \lambda_j r_E^j + I^j(t). \tag{43}$$

The parameters $a, b, c, d, e, f$ relate to the original model's parameters as follows:

$$\begin{aligned}
a &= -1/\tau_E^m + w_{EE} = 0.03712/ms, &(44)\\
b &= w_{EI} = 0.08316/ms, &(45)\\
c &= \mu_{EE}, &(46)\\
d &= w_{IE} = 0.42822/ms, &(47)\\
e &= 1/\tau_I^m + w_{II} = 0.53875/ms, &(48)\\
f &= \mu_{IE}, &(49)\\
\lambda_j &= -1/\tau_E^j. &(50)
\end{aligned}$$

We express $br_I^i$ in terms of other variables based on (41),

$$br_I^i = ar_E^i + cr_E^j + I^i(t) - \frac{dr_E^i}{dt}. \tag{51}$$

This allows us to reformulate the system:

$$\begin{aligned}
\frac{d^2 r_E^i}{dt^2} &= a\frac{dr_E^i}{dt} - b\frac{dr_I^i}{dt} + c\frac{dr_E^j}{dt} + \frac{dI^i(t)}{dt}\\
&= a\frac{dr_E^i}{dt} - b\left(dr_E^i - er_I^i + fr_E^j\right) + c\frac{dr_E^j}{dt} + \frac{dI^i(t)}{dt}\\
&= -bdr_E^i + a\frac{dr_E^i}{dt} - bfr_E^j + c\frac{dr_E^j}{dt} + ebr_I^i + \frac{dI^i(t)}{dt}\\
&= -bdr_E^i + a\frac{dr_E^i}{dt} - bfr_E^j + c\frac{dr_E^j}{dt} + e\left(ar_E^i + cr_E^j + I^i(t) - \frac{dr_E^i}{dt}\right) + \frac{dI^i(t)}{dt}\\
&= (ae - bd)\, r_E^i + (a - e)\frac{dr_E^i}{dt} + (ce - bf)\, r_E^j + c\frac{dr_E^j}{dt} + eI^i(t) + \frac{dI^i(t)}{dt} \qquad (52)
\end{aligned}$$

The reformulation results in a second-order ODE for $r_E^i$, capturing the dynamics without explicitly including $r_I^i$:

$$\frac{d^2 r_E^i}{dt^2} - (a - e)\frac{dr_E^i}{dt} + (bd - ae)\, r_E^i = (ce - bf)\, r_E^j + c\frac{dr_E^j}{dt} + eI^i(t) + \frac{dI^i(t)}{dt}. \tag{53}$$

##### 3.2.2 Non-dimensionalization leads to the initial layer problem

We now proceed to do the non-dimensionalization to (53) as the standard prepossessing required for the asymptotic analysis with the following transformation of variables:

$$R_E^i = (e - a)^{-1} r_E^i := \gamma^{-1} r_E^i, \tau = (ae - bd)(a - e)^{-1} t := \kappa t, \tag{54}$$

where

$$\gamma = e - a = 0.50163/ms, \quad \kappa = (ae - bd)(a - e)^{-1} = 0.03112/ms. \tag{55}$$

Now we have the dimensionless scalar $R_E^i, \tau$ satisfying the following second-order dynamics:

$$\kappa^2\gamma\frac{d^2R_E^i}{d\tau^2} - \kappa\gamma\left(a-e\right)\frac{dR_E^i}{d\tau} + \left(bd-ae\right)\gamma R_E^i = \left(ce-bf\right)r_E^j + c\kappa\frac{dr_E^j}{d\tau} + eI^i(\tau/\kappa) + \kappa\frac{dI^i(\tau/\kappa)}{d\tau}. \tag{56}$$

Dividing $\kappa\gamma(a-e)$ on both sides with some math simplification, we have

$$\frac{\kappa}{a-e}\frac{d^2R_E^i}{d\tau^2} - \frac{dR_E^i}{d\tau} + \frac{(bd-ae)}{\kappa(a-e)}R_E^i = \frac{1}{\kappa\gamma(a-e)}\left[\left(ce-bf\right)r_E^j + c\kappa\frac{dr_E^j}{d\tau} + eI^i(\tau/\kappa) + \kappa\frac{dI^i(\tau/\kappa)}{d\tau}\right]. \tag{57}$$

which equals to

$$\epsilon\frac{d^2R_E^i}{d\tau^2} - \frac{dR_E^i}{d\tau} - R_E^i = \frac{1}{\kappa\gamma(a-e)}\left[\left(ce-bf\right)r_E^j + c\kappa\frac{dr_E^j}{d\tau} + e\left(I^i(\tau/\kappa) + \xi\frac{dI^i(\tau/\kappa)}{d\tau}\right)\right]. \tag{58}$$

where

$$\epsilon = \frac{\kappa}{a-e} = -0.06204, \quad \xi = \frac{\kappa}{e} = 0.05777. \tag{59}$$

These non-dimensionalization leads to the classical "initial layer" asymptotic formula where the second order derivative has a very small coefficient, which restricts the role of high order derivatives to the early stages of the dynamics, with no significant affects on long-term behavior. An initial layer problem refers to a type of singularly perturbed differential equation in the temporal domain characterized by a large jump over a short interval to a smooth solution, analogous to the boundary layer problem in the spatial domain [5].

##### 3.2.3 An interpretation of dimensionless quantity $\epsilon$

Before delving into the further asymptotic analysis, we turn our attention to the dimensionless quantity $\epsilon$ and consider its implications when it assumes a small value. Starting from (59) and revisiting the original definitions of $\kappa, a, e$, we express $\epsilon$ as:

$$\epsilon = \frac{\kappa}{a-e} = \frac{(w_{EE} - 1/\tau_E^m)(w_{II} + 1/\tau_I^m) - w_{EI}w_{IE}}{\left[(w_{EE} - 1/\tau_E^m) - (w_{II} + 1/\tau_I^m)\right]^2}. \tag{60}$$

For an isolated downstream brain area $i$, devoid of projections from other areas, the dynamics are described by:

$$\frac{dr_E^i}{dt} = -\frac{1}{\tau_E^m}r_E^i + w_{EE}r_E^i - w_{EI}r_I^i + I^i(t), \tag{61}$$

$$\frac{dr_I^i}{dt} = -\frac{1}{\tau_I^m}r_I^i + w_{IE}r_E^i - w_{II}r_I^i. \tag{62}$$

The eigenvalues from this isolated system fulfill the characteristic equation:

$$[\lambda - (w_{EE} - 1/\tau_E^m)]\,[\lambda + (w_{II} + 1/\tau_I^m)] + w_{IE}w_{EI} = 0 \tag{63}$$

which leads to the following relationships between them (Vieta's formulas):

$$\begin{aligned} \lambda_1 + \lambda_2 &= (w_{EE} - 1/\tau_E^m) - (w_{II} + 1/\tau_I^m)\,, \\ \lambda_1\lambda_2 &= w_{IE}w_{EI} - (w_{EE} - 1/\tau_E^m)\,(w_{II} + 1/\tau_I^m)\,. \end{aligned} \tag{64}$$

Accordingly, $\epsilon$ can be recast as:

$$\epsilon = -\frac{\lambda_1\lambda_2}{(\lambda_1+\lambda_2)^2} = -\frac{1}{\left(\sqrt{r_\lambda} + \sqrt{1/r_\lambda}\right)^2} \tag{65}$$

where $r_\lambda = \lambda_1/\lambda_2$ represents the ratio of the two eigenvalues and, by extension, the ratio of the two intricate timescales governing the downstream area's isolated dynamics. It is important to note that when $\epsilon$ is close to 0, it implies that either $r_\lambda$ or $1/r_\lambda$ is considerably large, indicating a marked difference between the two timescales.

##### 3.2.4 Approximation of E to E-I model to E to E model

Based on the asymptotic analysis, the leading-order approximation of (58) is as follows:

$$-\frac{dR_E^i}{d\tau} - R_E^i = \frac{1}{\kappa\gamma(a-e)}\left[(ce - bf)\,r_E^j + c\kappa\frac{dr_E^j}{d\tau} + eI^i(\tau/\kappa)\right]. \tag{66}$$

Transforming $R_E^i, \tau$ back to the original variables, we have the following

$$-(a-e)\,\frac{dr_E^i}{dt} + (bd - ae)\,r_E^i = (ce - bf)\,r_E^j + c\frac{dr_E^j}{dt} + eI^i(t). \tag{67}$$

which equals to the first-order dynamics

$$\frac{dr_E^i}{dt} = -\frac{bd-ae}{e-a}r_E^i + \frac{ce-bf}{e-a}r_E^j + \frac{c}{e-a}\frac{dr_E^j}{dt} + \frac{e}{e-a}I^i(t). \tag{68}$$

By further denoting the coefficients as

$$T = e - a = w_{II} - w_{EE} + 1/\tau_E^m + 1/\tau_I^m, \tag{69}$$

$$\lambda_i = -\frac{bd-ae}{e-a} = -\frac{w_{EI}w_{IE} - (w_{EE} - 1/\tau_E^m)\,(w_{II} + 1/\tau_I^m)}{w_{II} - w_{EE} + 1/\tau_E^m + 1/\tau_I^m}, \tag{70}$$

$$\tau_E^i = -\frac{1}{\lambda_i} = \frac{w_{II} - w_{EE} + 1/\tau_E^m + 1/\tau_I^m}{w_{EI}w_{IE} - (w_{EE} - 1/\tau_E^m)\,(w_{II} + 1/\tau_I^m)}, \tag{71}$$

we have the approximated first-order dynamics of $r_E^i$ as follows:

$$\frac{dr_E^i}{dt} = -\frac{1}{\tau_E^i} r_E^i + \frac{1}{T}\left[\left(\left(w_{II} + 1/\tau_I^m\right)\mu_{EE} - w_{EI}\mu_{IE}\right) r_E^j + \mu_{EE}\frac{dr_E^j}{dt} + \left(w_{II} + 1/\tau_I^m\right) I^i(t)\right]. \tag{72}$$

#### 3.3 Metrics for signal integration and propagation

Drawing from the asymptotic analysis of the 'E-to-EI' model (Eq. (72)), we propose two quantitative metrics specifically designed to characterize timescale localization and signal propagation within this framework. This approach not only provides a more compact representation of the system but also offers deeper insights into the underlying behavioral dynamics of the neural model.

##### 3.3.1 Definition of two metrics

We first define **two metrics** $M_{SP}, M_{TL}$ as

$$M_{SP} = \left(w_{II} + 1/\tau_I^m\right)\mu_{EE} - w_{EI}\mu_{IE}, \tag{73}$$
$$M_{TL} = M_{SP} - \mu_{EE}/\tau_E^j. \tag{74}$$

Taking (73) and (74) back into the dynamics of $r_E^j$ (72), we have the following simplified representations of $r_E^i$ dynamics:

$$\frac{dr_E^i}{dt} = -\frac{1}{\tau_E^i} r_E^i + \frac{M_{SP}}{T} r_E^j + \frac{\mu_{EE}}{T}\frac{dr_E^j}{dt}, \tag{75}$$
$$= -\frac{1}{\tau_E^i} r_E^i + \frac{1}{T}\left(M_{SP} - \frac{1}{\tau_E^j}\mu_{EE}\right) r_E^j + \frac{\mu_{EE}}{T} I^j(t)$$
$$= -\frac{1}{\tau_E^i} r_E^i + \frac{M_{TL}}{T} r_E^j + \frac{\mu_{EE}}{T} I^j(t). \tag{76}$$

By decomposing the neuron's activity into its average components, $\bar{r}_E^i, \bar{r}_E^j$, and the temporally varying components, $\Delta r_E^i, \Delta r_E^j$, we establish the following relationships governed by $M_{SP}$ and $M_{TL}$:

$$\bar{r}_E^i = \frac{\tau_E^i M_{SP}}{T}\bar{r}_E^j, \tag{77}$$
$$\frac{d\Delta r_E^i}{dt} = -\frac{1}{\tau_E^i}\Delta r_E^i + \frac{M_{TL}}{T}\Delta r_E^j + \frac{\mu_{EE}}{T}\left(I^j(t) - \frac{1}{\tau_E^j}\bar{r}_E^j\right). \tag{78}$$

In the following discussion, we examine how (77) elucidates the relationship between changes in the magnitude of the signal $r_E^j$ and corresponding changes in $r_E^i$, a process indicative of signal propagation. Concurrently, (78) reveals the influence of fluctuations in the activity of $r_E^j$ on $r_E^i$, shedding light on the mechanism of timescale localization.

##### 3.3.2 $M_{SP}$ as a metric for signal propagation

For signal propagation, (77) directly points out that if we hold $r_E^j$ at a constant value $\bar{r}_E^j$, the steady state of $r_E^i$ would satisfy that:

$$\bar{r}_E^i = \frac{\tau_E^i M_{SP}}{T} \bar{r}_E^j, \tag{79}$$

which increases proportionally with $M_{SP}$. This further indicates that an increase of $r_E^j$ by $\Delta r_{E,i}$ results in an amplified response in the postsynaptic groups, approximately by $O\left(M_{SP} \Delta r_{E,i}\right)$.

##### 3.3.3 $M_{TL}$ as a metric for timescale localization

Our analysis now turns to timescale localization, specifically examining its role in two key scenarios: the resting state, characterized by white noise input, and the decay phase following stimulus presentation. We demonstrate that in both instances, the extent of timescale localization — reflected in the degree of independence among neural activities across varying timescales — can be effectively quantified by the metric $M_{TL}$.

***Signal integration during the resting state***

For the resting state, we consider (76) (with white noise input) such that

$$\frac{dr_E^i}{dt} = \lambda_i r_E^i + \frac{M_{TL}}{T} r_E^j + \frac{\mu_{EE}}{T} \sigma_j N^j(t) + \sigma_i N^i(t), \tag{80}$$

$$\frac{dr_E^j}{dt} = \lambda_j r_E^j + \sigma_j N^j(t). \tag{81}$$

where $N^i(t), N^j(t)$ are the white noise input and $\lambda_{i/j} = -\frac{1}{\tau_E^{i/j}}$. The formula of the system gives out the following theorem of the auto-correlation function of activity of brain area $i$.

**Theorem:** The auto-covariance function of activity of brain area $i$ satisfies the equation:

$$ACF^i(t) = \left( Var(r_E^i(0)) + \frac{M_{TL} Cov(r_E^i(0), r_E^j(0))}{T(\lambda_i - \lambda_j)} \right) e^{\lambda_i t} - \frac{M_{TL} Cov(r_E^i(0), r_E^j(0))}{T(\lambda_i - \lambda_j)} e^{\lambda_j t}. \tag{82}$$

**Proof:** Denoting

$$A = \begin{pmatrix} \lambda_i & \frac{M_{TL}}{T} \\ 0 & \lambda_j \end{pmatrix}, \quad \Sigma = \begin{pmatrix} \sigma_i & \frac{\mu_{EE}}{T} \sigma_j \\ 0 & \sigma_j \end{pmatrix}, \tag{83}$$

we know that

$$d \begin{pmatrix} r_E^i \\ r_E^j \end{pmatrix} / dt = A \begin{pmatrix} r_E^i \\ r_E^j \end{pmatrix} + \Sigma \begin{pmatrix} N^i(t) \\ N^j(t), \end{pmatrix} \tag{84}$$

Based on the knowledge of OU process, we know that the stationary distribution is a Gaussian distribution with

$$E(r_E^i) = 0, \quad E(r_E^j) = 0, \tag{85}$$

and covariance matrix $S$ satisfies

$$AS + SA^t = -\Sigma\Sigma^t. \tag{86}$$

It leads to the solution of $S$:

$$S = \begin{pmatrix} -\frac{\sigma_i^2}{2\lambda_i} - \frac{M_{TL}\sigma_j^2}{2T\lambda_i\lambda_j(\lambda_i+\lambda_j)} + \frac{M_{TL}\mu_{EE}\sigma_j^2}{T^2\lambda_i(\lambda_i+\lambda_j)} - \frac{\mu_{EE}^2\sigma_j^2}{2T^2\lambda_i} & \frac{(M_{TL}/2-\lambda_j\mu_{EE})\sigma_j^2}{T\lambda_j(\lambda_i+\lambda_j)} \\ \frac{(M_{TL}/2-\lambda_j\mu_{EE})\sigma_j^2}{T\lambda_j(\lambda_i+\lambda_j)} & -\frac{\sigma_j^2}{2\lambda_j} \end{pmatrix} \tag{87}$$

Hence we know

$$Var(r_E^i(0)) = \frac{\sigma_i^2}{2\lambda_i} - \frac{M_{TL}\sigma_j^2}{2T\lambda_i\lambda_j(\lambda_i+\lambda_j)} + \frac{M_{TL}\mu_{EE}\sigma_j^2}{T^2\lambda_i(\lambda_i+\lambda_j)} - \frac{\mu_{EE}^2\sigma_j^2}{2T^2\lambda_i}, \quad Var(r_E^j) = -\frac{\sigma_j^2}{2\lambda_j}. \tag{88}$$

and

$$Cov(r_E^i(0), r_E^j(0)) = \frac{(M_{TL}/2 - \lambda_j\mu_{EE})\,\sigma_j^2}{T\lambda_j\,(\lambda_i+\lambda_j)}. \tag{89}$$

Besides, by solving the OU equations, we have

$$\begin{aligned} r_E^i(t) &= e^{\lambda_i t} r_E^i(0) + \frac{e^{\lambda_i t} - e^{\lambda_j t}}{\lambda_i - \lambda_j}\frac{M_{TL}}{T} r_E^j(0) \\ &+ \int_0^t e^{\lambda_i(t-t')}\sigma_i dW_{t'}^i + \int_0^t \left[\frac{\mu_{EE}}{T}\sigma_j e^{\lambda_i(t-t')} + \frac{e^{\lambda_i(t-t')} - e^{\lambda_j(t-t')}}{\lambda_i-\lambda_j}\frac{M_{TL}}{T}\sigma_j\right] dW_{t'}^j \end{aligned} \tag{90}$$

Hence

$$\begin{aligned} Cov(r_E^i(t), r_E^i(s)) &= e^{\lambda_i(t+s)} Var(r_E^i(0)) + \frac{e^{\lambda_i t} - e^{\lambda_j t}}{\lambda_i - \lambda_j}\frac{e^{\lambda_i s} - e^{\lambda_j s}}{\lambda_i - \lambda_j}\frac{M_{TL}^2}{T^2} Var(r_E^j(0)) \\ &+ \left(e^{\lambda_i t}\frac{e^{\lambda_i s} - e^{\lambda_j s}}{\lambda_i - \lambda_j} + e^{\lambda_i s}\frac{e^{\lambda_i t} - e^{\lambda_j t}}{\lambda_i - \lambda_j}\right)\frac{M_{TL}}{T} Cov(r_E^i(0), r_E^j(0)) \\ &+ Cov\left(\int_0^t e^{\lambda_i(t-t')}\sigma_i dW_{t'}^i, \int_0^s e^{\lambda_i(s-t')}\sigma_i dW_{t'}^i\right) \\ &+ Cov\left(\int_0^t \left[\frac{\mu_{EE}}{T}\sigma_j e^{\lambda_i(t-t')} + \frac{e^{\lambda_i(t-t')} - e^{\lambda_j(t-t')}}{\lambda_i - \lambda_j}\frac{M_{TL}}{T}\sigma_j\right] dW_{t'}^j,\right. \\ &\left.\int_0^s \left[\frac{\mu_{EE}}{T}\sigma_j e^{\lambda_i(s-t')} + \frac{e^{\lambda_i(s-t')} - e^{\lambda_j(s-t')}}{\lambda_i - \lambda_j}\frac{M_{TL}}{T}\sigma_j\right] dW_{t'}^j\right). \end{aligned}$$

Thus

$$\begin{aligned} ACF^i(t) &= Cov(r_E^i(t), r_E^i(0)) = e^{\lambda_i t} Var(r_E^i(0)) + \frac{e^{\lambda_i t} - e^{\lambda_j t}}{\lambda_i - \lambda_j} \frac{M_{TL}}{T} Cov(r_E^i(0), r_E^j(0)) \\ &= \left( Var(r_E^i(0)) + \frac{M_{TL} Cov(r_E^i(0), r_E^j(0))}{T(\lambda_i - \lambda_j)} \right) e^{\lambda_i t} - \frac{M_{TL} Cov(r_E^i(0), r_E^j(0))}{T(\lambda_i - \lambda_j)} e^{\lambda_j t}. \end{aligned}$$

According to Theorem (82), $M_{TL}$ primarily determines the correlation between the two nodes, with a particular emphasis on the contribution from the timescale of $r_E^j$ (expressed as $e^{\lambda_j t}$). In the scenario where $M_{TL}$ approaches zero, the auto-correlation function (ACF) of $r_E^i$ converges to $ACF^i(t) \to Var(r_E^i(0)) e^{\lambda_i t} = Var(r_E^i(0)) e^{-t/\tau_E^i}$. This convergence signifies a state of perfect timescale localization.

***Signal integration after stimulus presentation***

For the post-stimulus scenario, we still consider (76) with $I^i(t) = I^j(t) = 0, r_E^j(t) = r_E^j(0) e^{\lambda_j t}$, we can analytically solve the system to find:

$$r_E^i(t) = \left[ r_E^i(0) + \frac{M_{TL} r_E^j(0)}{T(\lambda_i - \lambda_j)} \right] e^{\lambda_i t} - \frac{M_{TL} r_E^j(0)}{T(\lambda_i - \lambda_j)} e^{\lambda_j t}. \tag{91}$$

It is apparent that the metric $M_{TL}$ significantly affects the relative contributions of the two exponentials, thereby influencing the timescale localization. As $M_{TL}$ approaches zero, the behavior of $r_E^i(t)$ tends towards $r_E^i(0) e^{\lambda_i t} = r_E^i(0) e^{-t/\tau_E^i}$, indicating perfect timescale localization.

##### 3.3.4 Summary of two metrics

With the two metrics $M_{SP}$ and $M_{TL}$ defined above, we can now clearly describe both signal propagation and timescale localization between the two areas, as well as how these processes can be influenced by other parameters in the model. One key composite parameter is $T$, which in the simplified two-area model is defined as $T = w_{II} - w_{EE} + \frac{1}{\tau_E^m} + \frac{1}{\tau_I^m}$, and is reduced to $T = \frac{1}{\tau_E^i} + \frac{1}{\tau_I^i}$ in the minimal two-area model. Notably, in both cases, $T$ is governed by local parameters of the downstream node and remains independent of the upstream node, making it a node-specific parameter for the downstream node. To clarify how the upstream node influences the downstream dynamics, we summarize the three effects as follows:

1. The mean amplitude of the upstream node affects the mean amplitude of downstream activity, as described by the metric $M_{SP}$. The relationship is given by

$$\bar{r}_E^i = \frac{\tau_E^i M_{SP}}{T} \bar{r}_E^j, \tag{92}$$

   where the magnitude additionally depends on $\tau_E^i$ and $T$, both of which are local parameters of the downstream node.

2. The fluctuation of the upstream node adds an additional fluctuation component (with the upstream node's timescale) to the downstream node fluctuation, with its magnitude controlled by the metric $M_{TL}$. The exact formula is given as

$$\frac{d\Delta r_E^i}{dt} = -\frac{1}{\tau_E^i}\Delta r_E^i + \frac{M_{TL}}{T}\Delta r_E^j + \frac{\mu_{EE}}{T}\left(I^j(t) - \frac{1}{\tau_E^j}\bar{r}_E^j\right). \tag{93}$$

Again, the exact magnitude will only additionally depend on the local parameter $T$.

3. The upstream node effectively transmits the external input it receives to downstream node's fluctuation, even if the downstream node does not directly receive that input. This is captured by the term $\frac{\mu_{EE}}{T}\left(I^j(t) - \frac{1}{\tau_E^j}\bar{r}_E^j\right)$ in the following formula

$$\frac{d\Delta r_E^i}{dt} = -\frac{1}{\tau_E^i}\Delta r_E^i + \frac{M_{TL}}{T}\Delta r_E^j + \frac{\mu_{EE}}{T}\left(I^j(t) - \frac{1}{\tau_E^j}\bar{r}_E^j\right). \tag{94}$$

which is additionally determined by $T$ and the connection strength $\mu_{EE}$ between the upstream node and downstream excitatory group. Note that, while the upstream node's mean amplitude and timescale additionally contribute a *constant* drive $\frac{\mu_{EE}}{T}\frac{\bar{r}_E^j}{\tau_E^j}$ to the downstream activity, this *constant* drive does not influence the timescale of the downstream activity fluctuations.

### 4 Text S3: Extensive Scenarios for the two-area models

In this section, we explore the simplified two-area model under various conditions, particularly on the asymptotic assumptions (different $\epsilon$ values) and the system's behavior in the frequency domain.

#### 4.1 Model approximation and IFP for different epsilon values

We begin by examining the necessity of the asymptotic condition in the simplified two-area model, where the parameter $\epsilon$ (defined in Eq.(59)) is assume to be close to 0.

To investigate whether violating the asymptotic conditions invalidates the approximation used to reduce the system (Eq. (58) to Eq.(72)) , we varied the local connection strengths to increase the magnitude of $\epsilon$. From our analysis, we have shown that a small magnitude of $\epsilon$ corresponds to a large ratio between the two eigenvalues of the local node system when it receives no upstream input (Eq.(65)). As $\epsilon$ increases, the two local timescales become less distinct. To ensure comparability with other numerical results, we adjusted the parameters so that the eigenvalue corresponding to the slow timescale remains unchanged, while the eigenvalue corresponding to the fast timescale decreases, resulting in a slower fast timescale compared to the conditions where the asymptotic assumption holds.

As shown in Figure S12A, when we varied the local connection strengths such that $\epsilon$ approaches it maximum magnitude ($\epsilon = -0.24$, while Eq.(65) leads to $\epsilon = -1/\left(\sqrt{r_\lambda} + \sqrt{1/r_\lambda}\right)^2 \geq -1/\left(2\sqrt{\sqrt{r_\lambda}\sqrt{1/r_\lambda}}\right)^2 = -0.25$, suggesting the maximum magnitude to be $-0.25$), we observed a small deviation between the approximation and the full system. However, this deviation remained minor. This observation is further supported by Figure S12B, where the relative error (in L2 norm) increases with the magnitude of $\epsilon$, but remains small ($2\% \sim 6\%$) over a wide range of $\epsilon$ values.

Additionally, we emphasize that while the reduction of the model to a simple first-order E-E model relies on the assumption of a small $\epsilon$, the core finding of our work—that upstream and downstream areas can operate on different timescales—is independent of this assumption.

Starting from Eq.(53), which is equivalent to the original two-dimensional ODE systems, we use the coefficients defined in Eq.(71), which leads to:

$$\frac{1}{T}\frac{d^2 r_E^i}{dt^2} + \frac{dr_E^i}{dt} + \frac{1}{\tau_E^i} r_E^i$$
$$= \frac{1}{T}\left[\left(\left(w_{II} + 1/\tau_I^m\right)\mu_{EE} - w_{EI}\mu_{IE}\right) r_E^j + \mu_{EE}\frac{dr_E^j}{dt} + \left(w_{II} + 1/\tau_I^m\right) I^i(t)\right]. \quad (95)$$

with

$$T = w_{II} - w_{EE} + 1/\tau_E^m + 1/\tau_I^m. \quad (96)$$

We then incorporate the definitions of the two metrics $M_{SP}, M_{TL}$ (Eq.(73), Eq.(74)), which yield:

$$\frac{1}{T}\frac{d^2 r_E^i}{dt^2} + \frac{dr_E^i}{dt} + \frac{1}{\tau_E^i} r_E^i = \frac{M_{SP}}{T} r_E^j + \frac{\mu_{EE}}{T}\frac{dr_E^j}{dt}, \tag{97}$$

$$= \frac{M_{TL}}{T} r_E^j + \frac{\mu_{EE}}{T} I^j(t). \tag{98}$$

Similarly, by decomposing the neuron's activity into its average components, $\bar{r}_E^i, \bar{r}_E^j$, and the temporally varying components, $\Delta r_E^i, \Delta r_E^j$, we establish the following relationships governed by $M_{SP}$ and $M_{TL}$:

$$\bar{r}_E^i = \frac{\tau_E^i M_{SP}}{T} \bar{r}_E^j, \tag{99}$$

$$\frac{1}{T}\frac{d^2 \Delta r_E^i}{dt^2} + \frac{d\Delta r_E^i}{dt} + \frac{1}{\tau_E^i}\Delta r_E^i = \frac{M_{TL}}{T}\Delta r_E^j + \frac{\mu_{EE}}{T}\left(I^j(t) - \frac{1}{\tau_E^j}\bar{r}_E^j\right). \tag{100}$$

As seen in Eq.(99), the relationship between $\bar{r}_E^i, \bar{r}_E^j$ and $M_{SP}$ remains unchanged, regardless of the value of $\epsilon$. We can also observe that the influence of the upstream node on the downstream node's timescale is governed by the right-hand side term in Eq.(100). The asymptotic parameter $\epsilon$, along with the second-order derivative term, only appears on the left-hand side, affecting the intrinsic temporal dynamics of the downstream activity.

Our numerical simulations further support this finding. As shown in Figure S12C, when the magnitude of $\epsilon$ is varied from small to large, there is only a minor change in the temporal behavior of the downstream node. Despite this, the temporal dynamics remain characterized by a double-exponential decay, governed by its two local timescales—both of which are shorter than the upstream node's timescale—with no intermingling of the upstream node's timescale. Furthermore, Figure S12D demonstrates that even with an almost maximal magnitude of $\epsilon$, the system still achieves both reliable signal propagation with large magnitude, when maintaining nearly perfect timescale localization (characterized by a near-zero interference metric). It is important to note that in this context, the interference metric is computed by fitting the response of the downstream area with a triple exponential decay function:

$$f(t) = A_1 \exp(-t/\tau_1) + A_2 \exp(-t/\tau_2) + A_3 \exp(-t/\tau_3). \tag{101}$$

Here $\tau_1$, $\tau_2$ represent the two timescales corresponding to the eigenvalues of the downstream node when receiving no upstream input, while $\tau_3$ is the effective characteristic timescales of the upstream area. The interference metric is then defined as the ratio $A_3/(A_1 + A_2 + A_3)$, which ranges from 0 (minimal interference) to 1 (maximal interference).

#### 4.2 The dynamic behavior of the model under oscillatory inputs

We now investigate the dynamic behavior of the model under oscillatory inputs, focusing on the system's response in the frequency domain. Similar to the timescale localization phenomenon observed in the temporal domain (independence of the nodes' timescales), we observe a significant independence in the frequency domain. Specifically, the system can be effectively described as two low-pass filters that independently process external inputs.

We start from the general solution of a first-order dynamic equation with characteristic timescale $\tau$ driven by an oscillatory external input with frequency $\omega$, which obeys the following equation:

$$\frac{dr}{dt} = -r/\tau + A\sin(\omega t), \quad r(0) = 0. \tag{102}$$

The solution to this equation is:

$$r(t) = \frac{A\tau}{\sqrt{1 + (\omega\tau)^2}} \sin(\omega t - \arctan(\omega\tau)), \tag{103}$$

where the phase shift $\arctan(\omega\tau)$ increases monotonically with $\tau$. Besides, the magnitude $\frac{A\tau}{\sqrt{1+(\omega\tau)^2}}$ indicates that $r(t)$ behaves as a low-pass filter of the oscillatory signal, where the frequency response decays with increasing frequency $\omega$. This indicates that $r(t)$ attenuates higher frequency components more than lower ones. The *cutoff frequency* $\omega_c$ (where the magnitude drops to $1/\sqrt{2}$ of its maximum value) can be computed as

$$\omega_c = \frac{1}{\tau}, \tag{104}$$

which indicates that the frequencies higher than $\omega_c$ will be significantly attenuated.

In the IFP regime, although the downstream node receives input exclusively from the upstream node (which itself acts as a low-pass filter), the IFP motif can reduce the filtering effect, potentially decreasing it to zero. This can be observed by both the reconstruction of high-frequency responses and the reduction of phase shift (temporal lag), as we will illustrate with the following numerical results.

First, we show how the downstream node can reconstruct the high-frequency components of the external signal, even after significant filtering by the upstream node. As shown in Figure S2A, we apply an external input composed of five sinusoidal functions with identical amplitude but different frequencies to the upstream node. The downstream node's activity closely tracks the external input and exhibits larger oscillation amplitudes compared to the upstream node. This observation is further supported by estimating the power spectrum density (PSD) of the activities. As shown in Figure S2B, even though the oscillatory external input is significantly filtered by the upstream node, the downstream node reconstructs these high-frequency components and shows a stronger response at higher frequencies than the upstream node. This is non-trivial, as it would not be possible if the downstream node were merely a low-pass filter of the upstream node's activity.

Next, we investigate whether the downstream node can also compensate for the temporal lag due to the filtering effect of the upstream node. As shown in Figure S2A, the downstream node tracks the external input faster than the upstream node. The observation is quantitatively verified by applying a single sinusoidal input to the system and systematically examining the relationship between the phase shift (temporal lag) of the downstream node and the value of $M_{TL}$. Figure S2C shows that decreasing $M_{TL}$ can directly decrease the temporal lag of downstream node activity, and this relationship is robust across a wide range of input frequencies.

In summary, the IFP mechanism allows the downstream node to overcome the filtering effect introduced by the upstream node, functioning effectively as a low-pass filter directly receiving the external input. This occurs despite the fact that the external input must propagate through the upstream node to reach the downstream node.

#### 4.3 The simplified two-area model with reciprocal connections

We now extend the simplified two-areas model to include reciprocal connections between the two nodes, as shown in Figure S5A. Using similar analytic techniques, we illustrate how the system can be reduced approximately to an E-E system with reciprocal connections, and define corresponding $M_{SP}, M_{TL}$ governing the signal propagation and timescale localization in this scenario. Further numerical results are presented to validate the analysis.

##### 4.3.1 Analysis of the simplified two-area model with reciprocal connections

The dynamics of the system are described by the following equations:

$$\frac{dr_E^i}{dt} = -\frac{1}{\tau_E^m} r_E^i + w_{EE}^{ii} r_E^i - w_{EI}^{ii} r_I^i + \mu_{EE}^{ij} r_E^j + I^i(t), \tag{105}$$

$$\frac{dr_I^i}{dt} = -\frac{1}{\tau_I^m} r_I^i + w_{IE}^{ii} r_E^i - w_{II}^{ii} r_I^i + \mu_{IE}^{ij} r_E^j, \tag{106}$$

$$\frac{dr_E^j}{dt} = -\frac{1}{\tau_E^m} r_E^j + w_{EE}^{jj} r_E^j - w_{EI}^{jj} r_I^j + \mu_{EE}^{ji} r_E^i + I^j(t), \tag{107}$$

$$\frac{dr_I^j}{dt} = -\frac{1}{\tau_I^m} r_I^j + w_{IE}^{jj} r_E^j - w_{II}^{jj} r_I^j + \mu_{IE}^{ji} r_E^i. \tag{108}$$

Similarly, we can define the parameters $a, b, c, d, e, f$, which are related to model's parameters, as follow:

$$a^i = -1/\tau_E^m + w_{EE}^{ii}, \qquad b^i = w_{EI}^{ii}, \qquad c^j = \mu_{EE}^{ij},$$
$$d^i = w_{IE}^{ii}, \qquad e^i = 1/\tau_I^m + w_{II}^{ii}, \qquad f^{ij} = \mu_{IE}^{ij},$$

$$a^j = -1/\tau_E^m + w_{EE}^{jj}, \qquad b^j = w_{EI}^{jj}, \qquad c^i = \mu_{EE}^{ji},$$
$$d^j = w_{IE}^{jj}, \quad e^j = 1/\tau_I^m + w_{II}^{jj}, \quad f^{ji} = \mu_{IE}^{ji}. \tag{109}$$

We then proceed to do the non-dimensionalization similar to the unidirectional case, as the standard prepossessing required for the asymptotic analysis with the following transformation of variables:

$$R_E^x = (e^x - a^x)^{-1} r_E^x := (\gamma^x)^{-1} r_E^x, \tau^x = (a^x e^x - b^x d^x)(a^x - e^x)^{-1} t := \kappa^x t, \quad x = i, j, \tag{110}$$

where

$$\gamma^x = e^x - a^x, \quad \kappa^x = (a^x e^x - b^x d^x)(a^x - e^x)^{-1}, \quad x = i, j. \tag{111}$$

Using this non-dimensionalization and applying the same technique in the previous section to transform a two-dimensional first-order ODEs to one second-order ODE, we can transform the original system into a pair of second-order ODEs:

$$\begin{aligned}
\epsilon^i \frac{d^2 R_E^i}{d\left(\tau^i\right)^2} - \frac{dR_E^i}{d\tau^i} - R_E^i &= \frac{1}{\kappa^i \gamma^i (a^i - e^i)} \left[ \left(c^j e^i - b^i f^j\right) r_E^j + c^j \kappa^i \frac{dr_E^j}{d\tau^i} + e^i \left( I^i(\tau^i/\kappa^i) + \xi^i \frac{dI^i(\tau^i/\kappa^i)}{d\tau^i} \right) \right], \\
\epsilon^j \frac{d^2 R_E^j}{d\left(\tau^j\right)^2} - \frac{dR_E^j}{d\tau^j} - R_E^j &= \frac{1}{\kappa^j \gamma^j (a^j - e^j)} \left[ \left(c^i e^j - b^j f^i\right) r_E^i + c^i \kappa^j \frac{dr_E^i}{d\tau^j} + e^j \left( I^j(\tau^j/\kappa^j) + \xi^j \frac{dI^j(\tau^j/\kappa^j)}{d\tau^j} \right) \right],
\end{aligned} \tag{112}$$

where

$$\epsilon^x = \frac{\kappa^x}{a^x - e^x}, \quad \xi^x = \frac{\kappa^x}{e^x}, \quad x = i, j. \tag{113}$$

Assuming both $\epsilon^i, \epsilon^j$ are small, the leading-order approximation of (112) can be obtained as follows:

$$\begin{aligned}
-\frac{dR_E^i}{d\tau^i} - R_E^i &= \frac{1}{\kappa^i \gamma^i (a^i - e^i)} \left[ \left(c^j e^i - b^i f^j\right) r_E^j + c^j \kappa^i \frac{dr_E^j}{d\tau^i} + e^i I^i(\tau^i/\kappa^i) \right], \\
-\frac{dR_E^j}{d\tau^j} - R_E^j &= \frac{1}{\kappa^j \gamma^j (a^j - e^j)} \left[ \left(c^i e^j - b^j f^i\right) r_E^i + c^i \kappa^j \frac{dr_E^i}{d\tau^j} + e^j I^j(\tau^j/\kappa^j) \right].
\end{aligned} \tag{114}$$

Transforming $R_E^i, \tau$ back to the original variables and reorganizing the formulas, we obtain the approximated first-order dynamics of $r_E^i, r_E^j$ as follows:

$$\begin{aligned}
\frac{dr_E^i}{dt} &= -\frac{1}{\tau_E^i} r_E^i + \frac{1}{T^i} \left[ \left( \left(w_{II}^{ii} + 1/\tau_I^m\right) \mu_{EE}^{ij} - w_{EI}^{ii} \mu_{IE}^{ij} \right) r_E^j + \mu_{EE}^{ij} \frac{dr_E^j}{dt} + \left(w_{II}^{ii} + 1/\tau_I^m\right) I^i(t) \right]. \\
\frac{dr_E^j}{dt} &= -\frac{1}{\tau_E^j} r_E^j + \frac{1}{T^j} \left[ \left( \left(w_{II}^{jj} + 1/\tau_I^m\right) \mu_{EE}^{ji} - w_{EI}^{jj} \mu_{IE}^{ji} \right) r_E^i + \mu_{EE}^{ji} \frac{dr_E^i}{dt} + \left(w_{II}^{jj} + 1/\tau_I^m\right) I^j(t) \right].
\end{aligned}$$

(115)

where

$$T^x = e^x - a^x = w_{II}^{xx} - w_{EE}^{xx} + 1/\tau_E^m + 1/\tau_I^m, \tag{116}$$

$$\tau_E^x = \frac{w_{II}^{xx} - w_{EE}^{xx} + 1/\tau_E^m + 1/\tau_I^m}{w_{EI}^{xx} w_{IE}^{xx} - \left(w_{EE}^{xx} - 1/\tau_E^m\right)\left(w_{II}^{xx} + 1/\tau_I^m\right)}, \quad x = i, j. \tag{117}$$

Similarly, we can then define **two metrics** $M_{SP}, M_{TL}$ as

$$M_{SP}^{ij} = \left(w_{II}^{ii} + 1/\tau_I^m\right)\mu_{EE}^{ij} - w_{EI}^{ii}\mu_{IE}^{ij}, \quad M_{SP}^{ji} = \left(w_{II}^{jj} + 1/\tau_I^m\right)\mu_{EE}^{ji} - w_{EI}^{jj}\mu_{IE}^{ji}, \tag{118}$$

$$M_{TL}^{ij} = M_{SP}^{ij} - \mu_{EE}^{ij}/\tau_E^j, \quad M_{TL}^{ji} = M_{SP}^{ji} - \mu_{EE}^{ji}/\tau_E^i, \tag{119}$$

Taking (118) and (119) back into the dynamics of $r_E^j$ (115), we have the following simplified representations of $r_E^i, r_E^j$ dynamics:

$$\begin{aligned} \frac{dr_E^i}{dt} &= -\frac{1}{\tau_E^i} r_E^i + \frac{M_{SP}^{ij}}{T^i} r_E^j + \frac{\mu_{EE}^{ij}}{T^i}\frac{dr_E^j}{dt} + \frac{w_{II}^{ii} + 1/\tau_I^m}{T^i} I^i(t), \\ \frac{dr_E^j}{dt} &= -\frac{1}{\tau_E^j} r_E^j + \frac{M_{SP}^{ji}}{T^j} r_E^i + \frac{\mu_{EE}^{ji}}{T^j}\frac{dr_E^i}{dt} + \frac{w_{II}^{jj} + 1/\tau_I^m}{T^j} I^j(t), \end{aligned} \tag{120}$$

Furthermore, we have for $r_E^i$,

$$\begin{aligned} \frac{dr_E^i}{dt} &= -\frac{1}{\tau_E^i} r_E^i + \frac{1}{T^i}\left(M_{SP}^{ij} - \frac{1}{\tau_E^j}\mu_{EE}^{ij}\right) r_E^j + \frac{w_{II}^{ii} + 1/\tau_I^m}{T^i} I^i(t) \\ &+ \frac{\mu_{EE}^{ij}}{T^i}\left(\frac{M_{SP}^{ji}}{T^j} r_E^i + \frac{\mu_{EE}^{ji}}{T^j}\frac{dr_E^i}{dt} + \frac{w_{II}^{jj} + 1/\tau_I^m}{T^j} I^j(t)\right) \\ &= -\frac{1}{\tau_E^i} r_E^i + \frac{M_{TL}^{ij}}{T^i} r_E^j + \frac{w_{II}^{ii} + 1/\tau_I^m}{T^i} I^i(t) \\ &+ \frac{\mu_{EE}^{ij}\left(M_{TL}^{ji} + \mu_{EE}^{ji}/\tau_E^i\right)}{T^i T^j} r_E^i + \frac{\mu_{EE}^{ij}\mu_{EE}^{ji}}{T^i T^j}\frac{dr_E^i}{dt} + \frac{\mu_{EE}^{ij}\left(w_{II}^{jj} + 1/\tau_I^m\right)}{T^i T^j} I^j(t) \end{aligned} \tag{121}$$

The simplification leads to that

$$\frac{dr_E^i}{dt} = \frac{1}{1 - \frac{\mu_{EE}^{ij}\mu_{EE}^{ji}}{T^i T^j}}\left[-\left(\frac{1}{\tau_E^i} - \frac{\mu_{EE}^{ij}\left(M_{TL}^{ji} + \mu_{EE}^{ji}/\tau_E^i\right)}{T^i T^j}\right) r_E^i + \frac{M_{TL}^{ij}}{T^i} r_E^j \right. \tag{122}$$

$$\left. + \frac{w_{II}^{ii} + 1/\tau_I^m}{T^i} I^i(t) + \frac{\mu_{EE}^{ij}\left(w_{II}^{jj} + 1/\tau_I^m\right)}{T^i T^j} I^j(t)\right]. \tag{123}$$

Similarly for $r_E^j$, we then have

$$\frac{dr_E^j}{dt} = \frac{1}{1-\frac{\mu_{EE}^{ij}\mu_{EE}^{ji}}{T^iT^j}}\left[-\left(\frac{1}{\tau_E^j}-\frac{\mu_{EE}^{ji}\left(M_{TL}^{ij}+\mu_{EE}^{ij}/\tau_E^j\right)}{T^iT^j}\right)r_E^j+\frac{M_{TL}^{ji}}{T^j}r_E^i\right. \tag{124}$$

$$\left.+\frac{w_{II}^{jj}+1/\tau_I^m}{T^j}I^j(t)+\frac{\mu_{EE}^{ji}\left(w_{II}^{ii}+1/\tau_I^m\right)}{T^iT^j}I^i(t)\right]. \tag{125}$$

By decomposing the population's activity and external inputs into its average components, $\bar{r}_E^i, \bar{r}_E^j, \bar{I}^i, \bar{I}^j$, and the temporally varying components, $\Delta r_E^i, \Delta r_E^j, \Delta I^i, \Delta I^j$, we establish the following relationships governed by $M_{SP}^{ij}, M_{SP}^{ji}$ and $M_{TL}^{ij}, M_{TL}^{ji}$:

$$\bar{r}_E^i = \frac{\tau_E^i}{T^i}\left[M_{SP}^{ij}\bar{r}_E^j+\left(w_{II}^{ii}+1/\tau_I^m\right)\bar{I}^i\right], \quad \bar{r}_E^j = \frac{\tau_E^j}{T^j}\left[M_{SP}^{ji}\bar{r}_E^i+\left(w_{II}^{jj}+1/\tau_I^m\right)\bar{I}^j\right], \tag{126}$$

$$\frac{d\Delta r_E^i}{dt} = -\frac{1}{1-\psi}\left[-\frac{1}{\tilde{\tau}_E^i}\Delta r_E^i+\frac{M_{TL}^{ij}}{T^i}\Delta r_E^j+\frac{w_{II}^{ii}+1/\tau_I^m}{T^i}\Delta I^i+\frac{\mu_{EE}^{ij}\left(w_{II}^{jj}+1/\tau_I^m\right)}{T^iT^j}\Delta I^j\right],$$

$$\frac{d\Delta r_E^j}{dt} = -\frac{1}{1-\psi}\left[-\frac{1}{\tilde{\tau}_E^j}\Delta r_E^j+\frac{M_{TL}^{ji}}{T^j}\Delta r_E^i+\frac{w_{II}^{jj}+1/\tau_I^m}{T^j}\Delta I^j+\frac{\mu_{EE}^{ji}\left(w_{II}^{ii}+1/\tau_I^m\right)}{T^iT^j}\Delta I^i\right], \tag{127}$$

where $\psi = \mu_{EE}^{ij}\mu_{EE}^{ji}/\left(T^iT^j\right)$ and

$$\tilde{\tau}^x = \tau_E^x/\left[1-\tau_E^x\cdot\frac{\mu_{EE}^{xy}\left(M_{TL}^{yx}+\mu_{EE}^{yx}/\tau_E^x\right)}{T^xT^y}\right] = \tau_E^x/\left[1-\psi-\frac{\mu_{EE}^{xy}\tau_E^x}{T^xT^y}M_{TL}^{yx}\right], \quad (x,y)=(i,j) \text{ or } (j,i). \tag{128}$$

As we can see, (126) elucidates the relationship between changes in the magnitude of the signal $r_E^j$ and corresponding changes in $r_E^i$. Similarly, (127) reveals the influence of fluctuations in the activity of $r_E^j$ on $r_E^i$. Specifically, the metric $M_{TL}$ can affect the timescale localization in three key ways (which are slightly more complex than in the unidirectional case): (1) Similar to the uni-directional case, $M_{TL}$ controls the magnitude of the fluctuating activity component from the opposite node within the activity of the local node, as show in (127). (2) Additonaly, $M_{TL}$ can directly modify the characteristic time constant of each node, deviating from its value when there are no connections, as shown in (128). (3) The reciprocal connections create a mixture of two activity components in both nodes, introducing a second-order effect due to the circularity of the feedback. However, these effects can be negligible when $M_{TL}$ is small, as required in the IFP regime.

##### 4.3.2 Numerical results of the simplified two-area model with reciprocal connections

Next, we turn to the numerical validation of our analysis under the reciprocal condition. We first show that the E-to-E-I model with reciprocal connections can still achieve both intra-areal signal integration with distinct timescales and reliable inter-areal signal propagation. In Figure S5B, we present two scenarios where the nodes have weak and strong coupling strengths, both receiving external input. In both cases, the neuronal populations maintain distinct timescales, independent of the response magnitude.

We also examine the utility of the two metrics $M_{SP}, M_{TL}$ in quantifying signal propagation and timescale localization, respectively. It is noticeable that for each metric, we now have two values corresponding to the two directed connection ($i \to j$ and $j \to i$ ) that describe the behavior for each direction. Figure S5C shows that the amplitude of the node $i$ ($j$) response increases linearly with $M_{SP}^{ij}$ ($M_{SP}^{ji}$). Figure S5D shows that increasing the opposite population activity by $\eta$ proportionally enhances the activity component matching the corresponding timescale by $M_{TL}^{ij}\eta$ ($M_{TL}^{ji}\eta$) in the post-stimulus conditions.

#### 4.4 The simplified model with multiple upstream areas

We now extend the simplified model to include multiple upstream nodes (Figure S6A), each operating with a distinct timescale, connected to a single downstream node. Applying similar analytic techniques, we illustrate how the asymptotic assumptions remain valid and define corresponding metrics $M_{SP}$ and $M_{TL}$ governing the timescale localization and signal propagation in this scenario. Further numerical results are presented to validate the analysis.

##### 4.4.1 Analysis of the simplified model with multiple upstream areas

The dynamics of the system are now described by:

$$\frac{dr_E^i}{dt} = -\frac{1}{\tau_E^m} r_E^i + w_{EE} r_E^i - w_{EI} r_I^i + \sum_{j=1}^{N} \mu_{EE}^{ij} r_E^j + I^i(t), \tag{129}$$

$$\frac{dr_I^i}{dt} = -\frac{1}{\tau_I^m} r_I^i + w_{IE} r_E^i - w_{II} r_I^i + \sum_{j=1}^{N} \mu_{IE}^{ij} r_E^j, \tag{130}$$

$$\frac{dr_E^j}{dt} = -\frac{1}{\tau_E^j} r_E^j + I^j(t), \quad 1 \le j \le N. \tag{131}$$

We can then define the following parameters $a, b, c, d, e, f$, which relate to the model's parameters, as follows:

$$a = -1/\tau_E^m + w_{EE} = 0.03712/ms, \qquad b = w_{EI} = 0.08316/ms, \qquad c^j = \mu_{EE}^{ij}, \tag{132}$$

$$d = w_{IE} = 0.42822/ms, \qquad e = 1/\tau_I^m + w_{II} = 0.53875/ms, \qquad f^j = \mu_{IE}^{ij}, \tag{133}$$

$$\lambda_j = -1/\tau_E^j, \quad 1 \le j \le N. \tag{134}$$

Substituting these into the governing equations, we have:

$$\frac{dr_E^i}{dt} = ar_E^i - br_I^i + \sum_{j=1}^N c^j r_E^j + I^i(t), \tag{135}$$

$$\frac{dr_I^i}{dt} = dr_E^i - er_I^i + \sum_{j=1}^N f^j r_E^j, \tag{136}$$

$$\frac{dr_E^j}{dt} = \lambda_j r_E^j + I^j(t), \quad 1 \le j \le N. \tag{137}$$

We can transform these equations into a second-order ODE, similar to the single upstream node case, (substituting $cr_E^j$ as $\sum_{j=1}^N c^j r_E^j$ and $fr_I^j$ as $\sum_{j=1}^N f^j r_E^j$ for Eq.(53))

$$\frac{d^2 r_E^i}{dt^2} - (a-e)\frac{dr_E^i}{dt} + (bd - ae)\, r_E^i = \sum_{j=1}^N \left[ \left(c^j e - bf^j\right) r_E^j + c^j \frac{dr_E^j}{dt} \right] + eI^i(t) + \frac{dI^i(t)}{dt}. \tag{138}$$

We now proceed to do the non-dimensionalization to (138) as the standard prepossessing required for the asymptotic analysis with the following transformation of variables:

$$R_E^i = (e-a)^{-1} r_E^i := \gamma^{-1} r_E^i, \tau = (ae - bd)(a-e)^{-1} t := \kappa t, \tag{139}$$

where

$$\gamma = e - a = 0.50163/ms, \quad \kappa = (ae - bd)(a-e)^{-1} = 0.03112/ms. \tag{140}$$

This gives us the dimensionless scalar $R_E^i, \tau$ satisfying the following second-order dynamics:

$$\begin{aligned} &\kappa^2 \gamma \frac{d^2 R_E^i}{d\tau^2} - \kappa\gamma\,(a-e)\frac{dR_E^i}{d\tau} + (bd - ae)\,\gamma R_E^i \\ &= \sum_{j=1}^N \left[ \left(c^j e - bf^j\right) r_E^j + c^j \kappa \frac{dr_E^j}{d\tau} \right] + eI^i(\tau/\kappa) + \kappa \frac{dI^i(\tau/\kappa)}{d\tau}. \end{aligned} \tag{141}$$

which leads to

$$\epsilon \frac{d^2 R_E^i}{d\tau^2} - \frac{dR_E^i}{d\tau} - R_E^i = \frac{1}{\kappa\gamma(a-e)} \left[ \sum_{j=1}^N \left( \left(c^j e - bf^j\right) r_E^j + c^j \kappa \frac{dr_E^j}{d\tau} \right) + e\left( I^i(\tau/\kappa) + \xi \frac{dI^i(\tau/\kappa)}{d\tau} \right) \right]. \tag{142}$$

where

$$\epsilon = \frac{\kappa}{a-e} = -0.06204, \quad \xi = \frac{\kappa}{e} = 0.05777. \tag{143}$$

Based on the asymptotic analysis, the leading-order approximation of (142) is as follows:

$$-\frac{dR_E^i}{d\tau} - R_E^i = \frac{1}{\kappa\gamma(a-e)}\left[\sum_{j=1}^{N}\left(\left(c^j e - bf^j\right) r_E^j + c^j \kappa \frac{dr_E^j}{d\tau}\right) + eI^i(\tau/\kappa)\right]. \tag{144}$$

Transforming $R_E^i, \tau$ back to the original variables and reorganizing the formulas, we have the following first-order dynamics

$$\frac{dr_E^i}{dt} = -\frac{bd-ae}{e-a} r_E^i + \sum_{j=1}^{N}\left(\frac{c^j e - bf^j}{e-a} r_E^j + \frac{c^j}{e-a}\frac{dr_E^j}{dt}\right) + \frac{e}{e-a} I^i(t). \tag{145}$$

We similarly define the coefficients:

$$T = e - a = w_{II} - w_{EE} + 1/\tau_E^m + 1/\tau_I^m, \tag{146}$$

$$\lambda_i = -\frac{bd-ae}{e-a} = -\frac{w_{EI}w_{IE} - \left(w_{EE} - 1/\tau_E^m\right)\left(w_{II} + 1/\tau_I^m\right)}{w_{II} - w_{EE} + 1/\tau_E^m + 1/\tau_I^m}, \tag{147}$$

$$\tau_E^i = -\frac{1}{\lambda_i} = \frac{w_{II} - w_{EE} + 1/\tau_E^m + 1/\tau_I^m}{w_{EI}w_{IE} - \left(w_{EE} - 1/\tau_E^m\right)\left(w_{II} + 1/\tau_I^m\right)}, \tag{148}$$

we have the approximated first-order dynamics of $r_E^i$ as follows:

$$\frac{dr_E^i}{dt} = -\frac{1}{\tau_E^i} r_E^i + \frac{1}{T}\sum_{j=1}^{N}\left[\left(\left(w_{II} + 1/\tau_I^m\right)\mu_{EE}^{ij} - w_{EI}\mu_{IE}^{ij}\right) r_E^j + \mu_{EE}^{ij}\frac{dr_E^j}{dt}\right] + \frac{w_{II} + 1/\tau_I^m}{T} I^i(t). \tag{149}$$

Finally, we define **two metrics** $M_{SP}, M_{TL}$ as

$$M_{SP}^{ij} = \left(w_{II} + 1/\tau_I^m\right)\mu_{EE}^{ij} - w_{EI}\mu_{IE}^{ij}, \tag{150}$$

$$M_{TL}^{ij} = M_{SP}^{ij} - \mu_{EE}^{ij}/\tau_E^j, \quad 1 \le j \le N. \tag{151}$$

Substituting (150) and (151) back into the dynamics of $r_E^j$ (149), we obtain the following simplified representations of $r_E^i$ dynamics:

$$\begin{aligned}\frac{dr_E^i}{dt} &= -\frac{1}{\tau_E^i} r_E^i + \sum_{j=1}^{N}\left[\frac{M_{SP}^{ij}}{T} r_E^j + \frac{\mu_{EE}^{ij}}{T}\frac{dr_E^j}{dt}\right], \\ &= -\frac{1}{\tau_E^i} r_E^i + \sum_{j=1}^{N}\left[\frac{1}{T}\left(M_{SP}^{ij} - \frac{1}{\tau_E^j}\mu_{EE}^{ij}\right) r_E^j + \frac{\mu_{EE}^{ij}}{T} I^j(t)\right]\end{aligned} \tag{152}$$

$$= -\frac{1}{\tau_E^i} r_E^i + \sum_{j=1}^{N} \frac{M_{TL}^{ij}}{T} r_E^j + \sum_{j=1}^{N} \frac{\mu_{EE}^{ij}}{T} I^j(t). \tag{153}$$

By decomposing the neuron's activity and external inputs into its average components, $\bar{r}_E^i, \bar{r}_E^j, \bar{I}^j$, and the temporally varying components, $\Delta r_E^i, \Delta r_E^j, \Delta I^j$, we establish the following relationships governed by $M_{SP}$ and $M_{TL}$ (neglecting the external input to the downstream node $I^i$) :

$$\bar{r}_E^i = \frac{\tau_E^i}{T} \sum_{j=1}^{N} M_{SP}^{ij} \bar{r}_E^j, \quad \bar{r}_E^j = \tau_E^j \bar{I}^j, \tag{154}$$

$$\frac{d\Delta r_E^i}{dt} = -\frac{1}{\tau_E^i} \Delta r_E^i + \sum_{j=1}^{N} \frac{M_{TL}^{ij}}{T} \Delta r_E^j + \sum_{j=1}^{N} \frac{\mu_{EE}^{ij}}{T} \Delta I^j(t). \tag{155}$$

Similarly, (154) elucidates the relationship between changes in the magnitude of the signal $r_E^j$ and corresponding changes in $r_E^i$. Concurrently, (155) reveals the influence of fluctuations in the activity of $r_E^j$ on $r_E^i$. Compared to the case of a single upstream node, the signal propagation and the timescale localization now will controlled by $\sum_{j=1}^{N} M_{SP}^{ij}$ and $\sum_{j=1}^{N} M_{TL}^{ij}$ , respectively. These quantities are approximately proportional to the in-degree of the downstream node, $N$.

##### 4.4.2 Numerical results of the simplified model with multiple upstream areas

Next, we numerically validate our analysis under the condition of multiple upstream nodes. First, we demonstrate that the E-to-E-I model, when connected to multiple upstream nodes, still achieves both intra-areal signal integration with distinct timescales and reliable inter-areal signal propagation. The timescale of the upstream nodes are chosen from 100 $ms$ to 500 $ms$ uniformly. Figure S6B shows the two scenarios where the the downstream node is connected weakly and strongly to the upstream nodes, respectively. In both cases, the downstream node maintains markedly distinct timescales, regardless of the response magnitude.

We then numerically verify how signal propagation and timescale localization depend on the in-degree of the downstream node. Figure S6C shows that the amplitude of the downstream node response increases linearly with the in-degree $N$, while Figure S6D demonstrates that increasing the upstream population activity by $\eta$ proportionally increase the interference by $N\eta$ in the post-stimulus condition, while the interference remains small given each $M_{TL}^{ij}$ is small. It is noteworthy that this relationship is not perfectly linear, which is probably attributed to the instability in fitting multiple exponential decay functions when computing the interference metric.

### 5 Text S4: The multi-regional network model

We now further extend our analysis from the two-area model to a larger network-level model, employing similar dynamic equations. Concurrently, we expand the scope of our metrics to encompass system-level characteristics.

#### 5.1 Model formulation and parameters

We built the multi-regional macaque cortical model using a set of ordinary differential equations, adapted from previous works of modeling macaque cortical network [1, 2]. Each cortical area was modeled with one excitatory and one inhibitory neuron groups, which is governed by the following dynamics:

$$\tau_E \frac{d}{dt} r_E^i = -r_E^i + \beta_E \left[ (1+\eta_E h_i)\, w_{EE} r_E^i - w_{EI} r_I^i + (1+\eta_E h_i)\, \mu_{EE} \sum_{j=1}^{N} FLN_{ij} r_E^j + I_{\text{ext,E}}^i \right]_+ \quad (156)$$

$$\tau_I \frac{d}{dt} r_I^i = -r_I^i + \beta_I \left[ (1+\eta_I h_i)\, w_{IE} r_E^i - w_{II} r_I^i + (1+\eta_I h_i)\, \mu_{IE} \sum_{j=1}^{N} FLN_{ij} r_E^j + I_{\text{ext,I}}^i \right]_+ \quad (157)$$

The external input to the $X$ population in area $i$ is expressed as $I_{ext,X}^i$. We used a threshold linear function $f(x) = \beta[x]_+$ for the gain function for both E and I populations, with gain factors $\beta = \beta_E$ and $\beta_I$, respectively. The gradient $h_i$ is a factor modulating both local intra-areal and long-range inter-areal excitatory inputs and varies across different areas. The scaling factor $\eta_X$ governs the magnitude of the modulation effect of the gradient on the $X$ population. Adapted from a previous study [3] to obtain desired $M_{TL}$ and $M_{SP}$, we set $\tau_E = 20\ ms, \tau_I = 10\ ms, \beta_E = 0.066\ Hz/pA, \beta_I = 0.351\ Hz/pA, \eta_E = 0.685, \eta_I = 0.745, w_{EE} = 24.4\ pA/Hz, w_{EI} = 19.7\ pA/Hz, w_{IE} = 11.66\ pA/Hz, w_{II} = 12.5\ pA/Hz, \mu_{EE} = 132.58\ pA/Hz, \mu_{IE} = 98.92\ pA/Hz$.

#### 5.2 Extension of two metrics to multi-region model

Expanding the metrics in the simplified two-area model ((73), (74)) to a scenario with multiple nodes, the formulae for the two metrics can be naturally generalized as:

$$M_{SP} = \frac{1}{N_{\text{area}}} \sum_{i=1}^{N_{\text{area}}} \frac{\beta_E}{\tau_E} \frac{\beta_I}{\tau_I} \left( \left[ w_{II} + \frac{1}{\beta_I} \right] (1+\eta_E h_i)\, \mu_{EE} - w_{EI}\, (1+\eta_I h_i)\, \mu_{IE} \right),$$

$$M_{TL} = \frac{1}{N_{\text{area}}} \sum_{i=1}^{N_{\text{area}}} \left| \frac{\beta_E}{\tau_E} \frac{\beta_I}{\tau_I} \left( \left[ w_{II} + \frac{1}{\beta_I} \right] (1+\eta_E h_i)\, \mu_{EE} - w_{EI}\, (1+\eta_I h_i)\, \mu_{IE} \right) + \bar{\lambda} \frac{\beta_E}{\tau_E} (1+\eta_E h_i)\, \mu_{EE} \right|$$

$$\bar{\lambda} = \frac{1}{N_{\text{area}}} \sum_{i=1}^{N_{\text{area}}} \lambda_i.$$

where $\lambda_i$ is the eigenvalue corresponding to node $i$ where all brain areas are isolated (i.e., the inverse of the intrinsic timescale of node $i$).

In general, a large-scale network model can be decomposed into basic motifs, including unidirectional connections, bidirectional connections, and convergent connections from multiple upstream nodes to a downstream node, etc. Having demonstrated that the metrics $M_{SP}$ and $M_{TL}$ remain effective when generalized from the unidirectional two-area model to both the bidirectional two-area model and the multi-upstream-area model, we expect these metrics can be effectively extended to a large-scale model. This extension can be represented as the total summation or, equivalently, the average of pairwise $M_{SP}$ and $M_{TL}$ values across all pairs of nodes. We note that these metrics represent averages across all cortex areas thus they do not fully capture the signal propagation or timescale localization between specific brain areas, which additionally depend on the connectome and the stimulus patterns.

#### 5.3 Discussion on signal attenuation in the multi-region model

The simulation of the model shows that signals can reliably propagate from sensory areas to association areas in the IFP regime, in contrast to the weak GBA regime. However, we note that the reduction of signal strength within the cortical network is about 2 orders of magnitude, which is much larger than that in the two-area model. This is because, in the cortical network, signals must traverse a series of interconnected areas to move from V1 to 24c. In addition, the connectivity strength between these areas is highly heterogeneous, spanning five orders of magnitude according to anatomy [6]. As a result, signal propagation through certain intermediate brain regions with extremely weak connection may experience much greater attenuation than in the simplified model, leading to a cumulative effect that further amplifies this attenuation.

In addition, we find that the drastic signal reduction of 2 orders of magnitude could be attributed to the limited coverage of the connectome we use in the model. As the complete connectome of the macaque monkey has not been measured in experiment yet, here we only consider a subgraph of the full cortex in the model. Therefore, some existing signal propagation paths from V1 to 24c are missing. We expect that the input signal will be less attenuated once we add more areas in the model.

#### 5.4 Multi-region model receiving oscillatory inputs

To further justify timescale localization in each area, here we drive the multi-region network model with a periodic signal applied to area V1, with input frequencies ranging from 0.25 to 16 Hz to cover the timescales of all areas. We then compute the gains from the population response of each area at different frequencies, finding that the gains closely followed the form

$$p(w) \propto \frac{1}{\sqrt{w_c^2 + w^2}}.$$

It indicates that the activity of each area can be well described by a first-order dynamics, with a constant characteristic timescale $\tau$ independent of input frequency, i.e.

$$\frac{dr}{dt} = -\frac{1}{\tau} r + A \sin(wt).$$

The characteristic cut-off frequency of this ”low-pass filter” is predicted by $w_c = 1/\tau$, where $\tau$ is estimated from the autocorrelation function of the population activity under white noise drive. The accuracy of this low-pass filter prediction across different input frequencies is shown in Figure S13. Results for V1 and 24c are provided in Figure S13A as two examples. To quantify the prediction accuracy, we used the root mean squared error (RMSE) between each area's response gain and its theoretical prediction. As shown in Figure S13B, the low RMSE values across all areas indicate that the characteristic frequencies of local network nodes are largely invariant to the changes of input period.

### 6 Text S5: The Multi-regional spiking neuron network model

#### 6.1 Description of the multi-regional spiking neuron network model

Here we build a spiking neural network (SNN) model to investigate the existence of the IPF regime. The spiking network is constructed with the parameters adapted from Ref. [7].

##### 6.1.1 LIF neuron

For each area, there are 800 excitatory neurons and 200 inhibitory neurons. The membrane potential dynamics of the leaky integrate-and-fire (LIF) neuron model are governed by the equation:

$$\tau \frac{dV}{dt} = -(V(t) - V_{\text{rest}}) + RI(t), \tag{158}$$

where the resting potential $V_{\text{rest}} = -70$ mV, reset potential $V_{\text{reset}} = -55$ mV, and threshold potential $V_{\text{th}} = -50$ mV. For excitatory (E) neurons, the membrane time constant $\tau = 20$ ms, membrane resistance $R = 40$ MΩ, and refractory period $\tau_{\text{ref}} = 2$ ms. For inhibitory (I) neurons, the membrane time constant $\tau = 10$ ms, membrane resistance $R = 50$ MΩ, and refractory period $\tau_{\text{ref}} = 1$ ms.

##### 6.1.2 Synapse

We use NMDA and AMPA for excitatory synapse projection and $\text{GABA}_A$ for inhibitory synapse projection. For NMDA receptors, the synaptic conductance dynamics are described by the following system of equations:

$$\frac{dg_{\text{NMDA}}}{dt} = -\frac{g_{\text{NMDA}}}{\tau_{\text{decay}}} + ax(1 - g_{\text{NMDA}}), \tag{159}$$

$$\frac{dx}{dt} = -\frac{x}{\tau_{\text{rise}}} + \sum_k \delta(t - t_j^k), \tag{160}$$

and the synaptic current is given by:

$$I_{\text{NMDA}} = g_{\text{NMDA}} \cdot w_{\text{NMDA}} \cdot (E_{\text{NMDA}} - V(t)) \cdot \left(1 + e^{-\alpha V} \frac{[Mg^{2+}]_o}{\beta}\right)^{-1}, \tag{161}$$

where the parameters are as follows: $\tau_{\text{decay}} = 100$ ms, $\tau_{\text{rise}} = 2$ ms, $a = 0.5$ ms$^{-1}$, $E_{\text{NMDA}} = 0$ mV, $[Mg^{2+}]_o = 1$ mM, $\alpha = 0.062$, and $\beta = 3.57$.

For AMPA receptors, the synaptic conductance follows

$$\frac{dg_{\text{AMPA}}}{dt} = -\frac{g_{\text{AMPA}}}{\tau_{\text{decay}}} + \sum_k \delta(t - t_j^k), \tag{162}$$

and the synaptic current is

$$I_{\text{AMPA}} = g_{\text{AMPA}} \cdot w_{\text{AMPA}} \cdot (E_{\text{AMPA}} - V(t)), \tag{163}$$

where $\tau_{\text{decay}} = 2$ ms and $E_{\text{AMPA}} = 0$ mV.

For GABA$_A$ receptors, the synaptic conductance is given by

$$\frac{dg_{\text{GABA}}}{dt} = -\frac{g_{\text{GABA}}}{\tau_{\text{decay}}} + \sum_k \delta(t - t_j^k), \tag{164}$$

and the synaptic current is

$$I_{\text{GABA}} = g_{\text{GABA}} \cdot w_{\text{GABA}} \cdot (E_{\text{GABA}} - V(t)), \tag{165}$$

with $\tau_{\text{decay}} = 5$ ms and $E_{\text{GABA}} = -70$ mV.

##### 6.1.3 Neuron connection

E neurons project to E and I neurons through both NMDA and AMPA synapses with a probability 0.1. I neurons project to E and I neurons through GABA$_A$ synapse with a probability 0.1. When the SNN includes more than one area, I neurons only project to the neurons within the same area. Each neuron receives Poisson background input through AMPA synapse.

##### 6.1.4 Two-areal case

In the two-area scenario, we consider the motif that the upstream area projects to the downstream area without feedback. We denote the excitatory neurons in the upstream and downstream areas as $E_u$ and $E_d$ respectively, and $I_u$ and $I_d$ for inhibitory neurons. We set the synapse weight (in the unit of nS): $w^u_{EE,\text{NMDA}} = 0.3168$, $w^u_{EE,\text{AMPA}} = w^u_{EE,\text{NMDA}}/3.3$, $w^u_{IE,\text{NMDA}} = 0.208$, $w^u_{IE,\text{AMPA}} = w^u_{IE,\text{NMDA}}/3.3$, $w^u_{EI,\text{GABA}} = 2.6$, $w^u_{II,\text{GABA}} = 2$, $w^d_{EE,\text{NMDA}} = 0.105336$, $w^d_{EE,\text{AMPA}} = w^d_{EE,\text{NMDA}}/3.3$, $w^d_{IE,\text{NMDA}} = 0.06916$, $w^d_{IE,\text{AMPA}} = w^d_{IE,\text{NMDA}}/3.3$, $w^d_{EI,\text{GABA}} = 2.6$, $w^d_{II,\text{GABA}} = 2$, $\mu_{EE,\text{NMDA}} = 0.33k_E$, $\mu_{EE,\text{AMPA}} = \mu_{EE,\text{NMDA}}/3.3$, $\mu_{IE,\text{NMDA}} = 0.26k_I$, $\mu_{IE,\text{AMPA}} = \mu_{IE,\text{NMDA}}/3.3$. The frequencies of the background inputs to the upstream E and I neurons are 1.8kHz and 1.6kHz, respectively; and those for the downstream E and I neurons are 1.9kHz and 1.6kHz, respectively. The synapse weights of background input to the E and I neurons are 2.1 nS and 1.62 nS, respectively. During stimulus period, the E neurons in the upstream area receive 76Hz external Poisson inputs with synapse weight 2.1 nS. For the weak GBA case, we set $k_E = 0.154$, $k_I = 0.35$; for the IFP case, we set $k_E = 2.124$, $k_I = 3$; for the strong GBA case, we set $k_E = 0.913$, $k_I = 1.1$.

##### 6.1.5 Multi-regional case

For the multi-regional case, we set the following synapse weights (in the unit of nS): $w^i_{EE,\text{NMDA}} = 0.10296 \cdot (1 + \eta h_i)$, $w^i_{EE,\text{AMPA}} = w^i_{EE,\text{NMDA}}/3.3$, $w^i_{IE,\text{NMDA}} = 0.0676 \cdot$

$(1+\eta h_i)$, $w^i_{IE,\mathrm{AMPA}} = w^i_{IE,\mathrm{NMDA}}/3.3$, $w^i_{EI,\mathrm{GABAa}} = 2.6$, $w^i_{II,\mathrm{GABAa}} = 2$, $\mu^{ij}_{EE,\mathrm{NMDA}} = 2.855952 \cdot FLN_{ij} \cdot (1+\eta h_i)$, $\mu^{ij}_{EE,\mathrm{NMDA}} = \mu^{ij}_{EE,\mathrm{NMDA}}/3.3$, $\mu^{ij}_{IE,\mathrm{NMDA}} = 3.12 \cdot FLN_{ij} \cdot (1+\eta h_i)$, $\mu^{ij}_{IE,\mathrm{AMPA}} = \mu^{ij}_{IE,\mathrm{NMDA}}/3.3$, $\eta = 1.5$. Each area receives background Poisson inputs, and the synapse weights of the background input to the E and I neurons are 2.1 nS and 1.62 nS, respectively. During stimulus period, the E neurons in area V1 receives 9kHz external Poisson input with synapse weights 2.1 nS.

##### 6.1.6 Poisson network

We also construct a Poisson network to evaluate the intrinsic timescale of a specific area in the spiking neural network. For a given area, we replace the inputs from each upstream area with Poisson spike trains, ensuring that these inputs does not contain any inherent timescales. In addition, the Poisson input rate for each upstream area is set to match the original firing rate of that area in the network. Consequently, this network effectively removes temporal information from the upstream areas while preserving the network state, enabling the measure of intrinsic timescale of the specific area.

#### 6.2 Data analysis and simulation

##### 6.2.1 Population firing rate

After simulating the network, the population firing rate is calculated using sliding time windows. For each 50 ms time window [8], sliding in 0.1 ms steps, the total spike count for each neuron groups is calculated. This count is then divided by both the number of neurons and the duration of the window. For each phase of the simulation—pre-stimulus, stimulus, and delay period—the mean firing rate is estimated after the firing rate has stabilized.

##### 6.2.2 Current

For the analyses of signal propagation and timescale localization, the mean and temporal fluctuating components of synaptic input currents are primarily relevant. For a group of neurons in a specific area, we calculate the group-averaged current across all neurons. To be consistent, we apply the same sliding window process used for the firing rate to the current time series. The total current to E neurons in a given area can be decomposed into several components: excitatory inter-areal current, excitatory intra-areal current, inhibitory intra-areal current, and external/background current. In particular, we focus on the excitatory inter-areal current (E current) and the inhibitory intra-areal current (I current) that reflect the timescales of upstream areas, and the net current as the sum of the E current and I currents to investigate the temporal cancellation of the two currents.

##### 6.2.3 Signal propagation and timescale localization metrics

Although the metrics of $M_{SP}$ and $M_{TL}$ derived from the linear rate model cannot be directly applied to the nonlinear spiking network, we identified signatures of the IFP regime in the spiking network by leveraging the mechanistic insights into IFP established from the rate model. To be specific, our analysis in the main text shows that a robust signal propagation depends on a high mean total current to the downstream E population, resulting from an imbalance between the mean E current and the mean I current. Additionally, an effective timescale localization is associated with minimal filtered fluctuating current driven by upstream timescales, due to substantial cancellation of temporal fluctuations between the E and I currents. Therefore, we use the mean of net current change received by E neurons in a given area to quantify signal propagation ability, and we calculate the coverage under the autocorrelation function of the net current to quantify timescale localization. Moreover, this coverage is then normalized by the maximum time lag used in the autocorrelation function calculation. The difference between this normalized coverage and that of the "Poisson" network indicates the extent of timescale cancellation for the SNN case. A value close to zero suggests that the temporal component of inter-areal projections is effectively canceled by the inhibitory neuron group.

##### 6.2.4 Simulation

We use BrainPy 2.6.0 [9] to simulate the SNN, with a 0.1 ms timestep. The simulation consists of a 5 s pre-stimulus period, a 150 s stimulus period (to ensure the stability of the autocorrelation function), and a 5 s delay period.

#### 6.3 Numerical Results of the two-areal spiking neuron network model

We first construct a simplified two-areal model, with an upstream area projecting to a downstream area without feedback projection. As shown in Figure S14A, during the pre-stimulus period, the upstream excitatory (E) neurons only receive background noise input without external input. Upon stimulus presentation, these neurons are subjected to a specific level of external Poisson input, which is subsequently removed during the delay period. The firing rate of the upstream E neurons is thus modulated by the external input, as shown in Figure S14B. We calculate the autocorrelation function (ACF) for the firing rate reaching steady state, and adopt an exponential fit to obtain the upstream timescale as 386 ms, as illustrated in Figure S14C. With different projection strengths, the downstream E neurons fire with different patterns. For the weak GBA regime (Figure S14D-E), the downstream neurons' firing rate is only slightly influenced by the upstream neurons ($\sim$1 Hz), and the timescale is independent for these two areas (upstream: 386 ms, downstream: 34.9 ms). For the IFP regime (Figure S14F-G), the signal reliably propagates from the upstream group to the downstream group ($\sim$10 Hz), while keeping the timescales independent (upstream: 386 ms, downstream: 42.5 ms). For the strong GBA regime (Figure S14H-I), the signal reliably propagates ($\sim$10 Hz), but the downstream timescale is significantly influenced by the upstream group (upstream: 386 ms, downstream: 359 ms).

We next analyze the group-averaged input current to the downstream E neurons in Figure S15. In the weak GBA regime (Figure S15A), where inter-areal connectivity strength is very low, the interaction from the upstream to the downstream area is small.

This leads to both a small mean of total current and small temporal fluctuations in the total input current, even when E and I synaptic inputs do not cancel. In the IFP regime (Figure S15B), the net current is strong indicating reliable signal propagation, while the timescale of the net current is close to zero, despite the E current and I current carry large timescales from upstream activities. The cancellation between the E and I currents indicates the timescale of the downstream area is minimally influenced by the upstream area. In the strong GBA regime (Figure S15C), the net current remains strong, but the net current carries the timescale from the upstream group, which breaks the timescale localization. For each regime, we also present the auto-correlation function of the net current of the Poisson network as the control case. Comparison with the Poisson network further demonstrates that the temporal components of the E and I currents are canceled in the IFP regime, but not in the strong GBA regime.

##### 6.4 Numerical Results of the multi-regional spiking neuron network model

We further construct a multi-regional SNN with experimentally measured connection strengths (FLN) and hierarchy values, and set the parameters that lead to the IFP regime. For the pre-stimulus period, there is no external input. During the stimulus period, V1 receives a certain level of external Poisson input. This input is then removed during the delay period. In Figure S9A, we measure the firing rate change for each area, showing that the signal reliably propagates from V1 to other areas. In Figure S9B, we use the same procedure as that in the two-areal case to obtain the timescale of each area from the auto-correlation function of its activity. The timescale is hierarchically organized, consistent with experimental findings [8]. The gradient values of excitation used in the model are shown in Figure S9C. As a spiking version of the rate model, the magnitude of areal activity and timescale are highly correlated with that of the rate model (Figure S9D-E). In addition, the timescale of each area in the network is highly correlated with its intrinsic timescale measured in the Poisson network, as shown in Figure S9F.

We next show the firing rates during the stimulus period of nine selected areas (in hierarchical order) in Figure S10A. As depicted in Figure S10B, the the temporal components of the E and I currents effectively cancel out, resulting in a net current with a small timescale. To quantitatively analyze this cancellation, we compute the coverage under the ACF curve of the net current and normalize it by the maximum time lag. The difference between this value and that of the ”Poisson” network is shown in Figure S10C (top panel). For most areas, this difference is close to zero, indicating that the multi-regional SNN operates in the IFP regime. Additionally, we show the mean net current change in Figure S10C (bottom panel). The significantly non-vanishing value of the mean current for most areas indicates the effective signal propagation in the network.
